# Targeted KRAS^G12V^ degradation elicits efficient and durable lung adenocarcinoma regression *in vivo*

**DOI:** 10.1101/2024.12.13.627539

**Authors:** Alberto Martín, Inés M. García-Pérez, Sonia San José, Pep Rojo, Carlos Riego-Mejías, Cristina Teodosio, Bárbara MG. Barbosa, Carolina Sánchez-Zarzalejo, Ignasi Folch-I-Casanovas, Antonia Odena Caballol, Sònia Jarió, Rodrigo Entrialgo, Marie-Julie Nokin, Diana Loa, Elizabeth Guruceaga, Camille Stephan-Otto Attolini, Chiara Ambrogio, Alberto Villanueva, Silvestre Vicent, Antoni Riera, David Santamaría, Cristina Mayor-Ruiz

## Abstract

Recent drug discovery breakthroughs led to the approval of KRAS^G12C^ inhibitors in lung adenocarcinoma (LUAD). Unfortunately, clinical responses remain limited due to rapid resistance onset. Proteolysis-targeting chimeras (PROTACs) have emerged as promising alternatives to traditional inhibition. However, there is limited mechanistic understanding of KRAS degradation *in vivo*. Here, we developed a preclinical LUAD mouse model and demonstrated that targeted oncogenic KRAS degradation induces rapid tumor regression. Transcriptional, histological, and immunophenotypic analyses revealed a substantial remodeling of the tumor microenvironment. Notably, disease relapse observed during long-term degrader treatment stems from proteolysis machinery dysregulation, indicating resistance mechanisms distinct from those reported upon KRAS inhibition. Our findings highlight the therapeutic potential of KRAS degradation in LUAD, offering insights into cell-intrinsic and extrinsic mechanisms driving durable antitumor responses and supporting further clinical exploration.

**SIGNIFICANCE:** Given the short duration of the clinical responses to KRAS inhibitors, complementary therapies are a dire medical need. Our preclinical findings endorse KRAS degradation as a therapeutic alternative in LUAD, where cell-intrinsic and extrinsic mechanisms drive tumor regression and durable therapeutic responses.

## INTRODUCTION

RAS GTPases are among the most frequently mutated oncoproteins in cancer with ∼30% of all human tumors harboring a mutation in either HRAS, NRAS, or KRAS isoforms^1^. The most common activating mutations occur at codons 12, 13 and 61^1,2^. Functionally, these mutations stabilize the active, GTP-bound (ON) state of RAS proteins, thereby increasing oncogenic flux through downstream effectors such as the MAPK pathway^3^. KRAS mutations are also found in nearly a quarter of all lung adenocarcinomas (LUAD), the deadliest type of cancer worldwide^1^. Although KRAS was one of the first oncogenes identified in human tumors, efforts at developing efficient inhibitors stalled for around four decades^4,5^. However, recent drug development breakthroughs led to the approval of the first KRAS^G12C^ inhibitors for LUAD patients, namely sotorasib (AMG510) and adagrasib (MRTX849), greatly renewing the hope of inhibiting oncogenic KRAS through pharmacological tools^6–11^. Unfortunately, the clinical efficacy of KRAS^G12C^ inhibitors rapidly declines due to the development of resistance^12–14^.

Furthermore, these drugs spare non-G12C mutants, which constitute most KRAS alterations in cancer. As such, many drug discovery programs are ongoing to identify inhibitors targeting of other KRAS mutants. For instance, a non-covalent KRAS^G12D^ inhibitor (MRTX1133) is already undergoing early clinical testing^15,16^. In addition, multi-KRAS inhibitors such as BI- 2865, capable of engaging a broad spectrum of KRAS alleles have been reported^17^.

Despite extensive advances in the treatment of cancer using inhibitors, resistance to this type of pharmacology is still a major medical problem, as observed with sotorasib and adagrasib. Indeed, preclinical data with additional mutant-KRAS inhibitors suggest extent and rate of response are far from complete^18,19^.

Intense efforts are devoted to exploit on- and off-target drug combinations. The combination of KRAS inhibitors with immunotherapy, with inhibitors of upstream and downstream effectors, or with other actionable vulnerabilities are expected to increase the efficacy of anti-KRAS therapies in the clinic^20–22^.

In addition to combination therapies, novel approaches of direct KRAS targeting are actively being pursued. The development of RAS therapeutics that operate via different mechanisms of action compared to inhibitors could help realize the full potential of direct RAS targeting for cancer treatment. In particular, hijacking cellular circuits with drugs that induce protein-protein proximity offers interesting therapeutic opportunities. RMC-4998 and RMC-6291, two covalent tri-complex inhibitors of active KRAS^G12C^, exploit the concept of chemically-induced protein proximity^23^. They are macrocyclic compounds derived from sanglifehrin A, a natural product that binds to cyclophilin A (CYPA) with high affinity. Upon binding CYPA, these tri-complex inhibitors remodel the surface of CYPA to create a binary complex with high affinity for active KRAS. The ternary complex sterically blocks KRAS– effector interactions and disrupts downstream signaling. RMC-6291 is currently in clinical trials (NCT05462717). Recently, the reversible RAS(ON) multi-selective inhibitor RMC-7977 was also reported, and the related compound RMC-6236 has also entered clinical testing (NCT05379985)^24^.

Another promising RAS therapeutic concept enabled by proximity-inducing pharmacology is “targeted protein degradation” (TPD). TPD depends on small molecules called ‘*degraders*’ that prompt the destruction of disease-relevant proteins by inducing proximity to effectors of the cellular proteolytic machinery —often an E3 ubiquitin ligase^25–27^. While inhibitors typically target a specific enzymatic function, degraders affect protein abundance comparably to a genetic perturbation. The event-driven mechanism of action of degraders compared with occupancy-based traditional inhibitors imparts several unique advantages related to target scope, target fate, potency, and selectivity^28^. For instance, degraders do not require high binding affinity for the target protein in order to achieve effective depletion, thus holding the promise to widening the druggable proteome. Two are the prototypical types of degraders: proteolysis-targeting chimeras (PROTACs) and molecular glue degraders. PROTACs feature a dedicated ligand for the target protein and a dedicated E3 binder, both connected by a linker. In contrast, molecular glue degraders are monovalent linker-less compounds and connect target and E3 in a highly cooperative manner, typically by binding only one of them in isolation. The availability of KRAS inhibitors quickly resulted in the development of initial PROTACs against KRAS^G12C^ and KRAS^G12D29–31^. The most clinically advanced PROTACs are ASP3082 and ASP4396, both KRAS^G12D^ degraders with phase I studies ongoing (NCT05382559 and NCT06364696 respectively)^32^. Recently, a multi-KRAS PROTAC showed degradation against 13 out of 17 of the most prevalent oncogenic KRAS alleles, including those not yet tractable by inhibitors^33^. Unfortunately, this chemical tool is unsuitable for long-term studies *in vivo*.

In summary, PROTACs have garnered great interest as means against KRAS-driven tumors, suggesting that they may be one of the next long sought-after therapeutic modalities tested in the clinic. Recent elegant studies with LUAD mouse models revealed that while *Kras^G12C^* ablation induced massive tumor regression and prevented resistance *in vivo*, KRAS^G12C^ inhibition caused a limited antitumor response similar to that observed in the clinic, including the rapid onset of resistance^20^. The prospect of chemical KRAS degradation mimicking the superior efficacy of genetic ablation compared to inhibition is certainly appealing.

Although the emphasis of TPD has been typically placed on expanding the druggable proteome, degraders can also help tackle resistance to current anticancer inhibitors^34^. The critical opportunity lies in the fact that different pharmacology exhibits distinct resistance mechanisms. Inhibitors typically lead to target-related resistance such as secondary mutations at the binding pocket that decrease inhibitor affinity. The resistance to degraders instead is mainly driven by dysregulated activity of the proteolytic machinery^35^. Leveraging these differences presents a strategic chance to defeat resistance^36–38^. From the perspective of overcoming resistance, and besides other interesting complementary capabilities, the availability of different pharmacological modalities against KRAS is expected to help achieve durable responses. There is, however, limited mechanistic understanding of KRAS degradation *in vivo*.

Here, we set out to evaluate the therapeutic potential of PROTAC-induced degradation of oncogenic KRAS *in vivo*. Given the utmost unmet clinical need, we focused on LUAD. Particularly, on KRAS^G12V^-driven LUAD, a KRAS mutant with no selective inhibitors or degraders available. To this end, we developed a first-in-class KRAS^G12V^-degradable LUAD mouse model by leveraging a tag-PROTAC chemogenetic approach. Tag-based technologies of protein degradation offer a loss-of-function chemical solution to resolve target biology in cells and in mice even before having the required chemistry in hand^39–43^. Chemical, tumor-specific modulation of KRAS^G12V^ protein abundance served here as a versatile strategy to probe the therapeutic potential of oncoprotein degradation and to selectively dissect the biological outcomes.

## RESULTS

### Novel LUAD KRAS^G12V^-degradable syngeneic mouse models enable PROTAC-dependent phenotyping *in vivo*

To explore the *in vivo* outcomes of acute KRAS^mut^ degradation in LUAD, we took advantage of the degradation TAG (dTAG) approach (Fig. 1A). In brief, this chemogenetic system uses a 12-kDa cytosolic prolyl isomerase engineered variant (FKBP12^F36V^) as a tag to make fusion proteins amenable to targeted degradation^39,40^. This tag-based approach leverages the potency of standardized PROTACs such as dTAG-13 and dTAG^V^-1, which hijack the endogenous E3 ubiquitin ligases CRL4^CRBN^ and CRL2^VHL^, respectively^39,40^.

**Figure 1.**
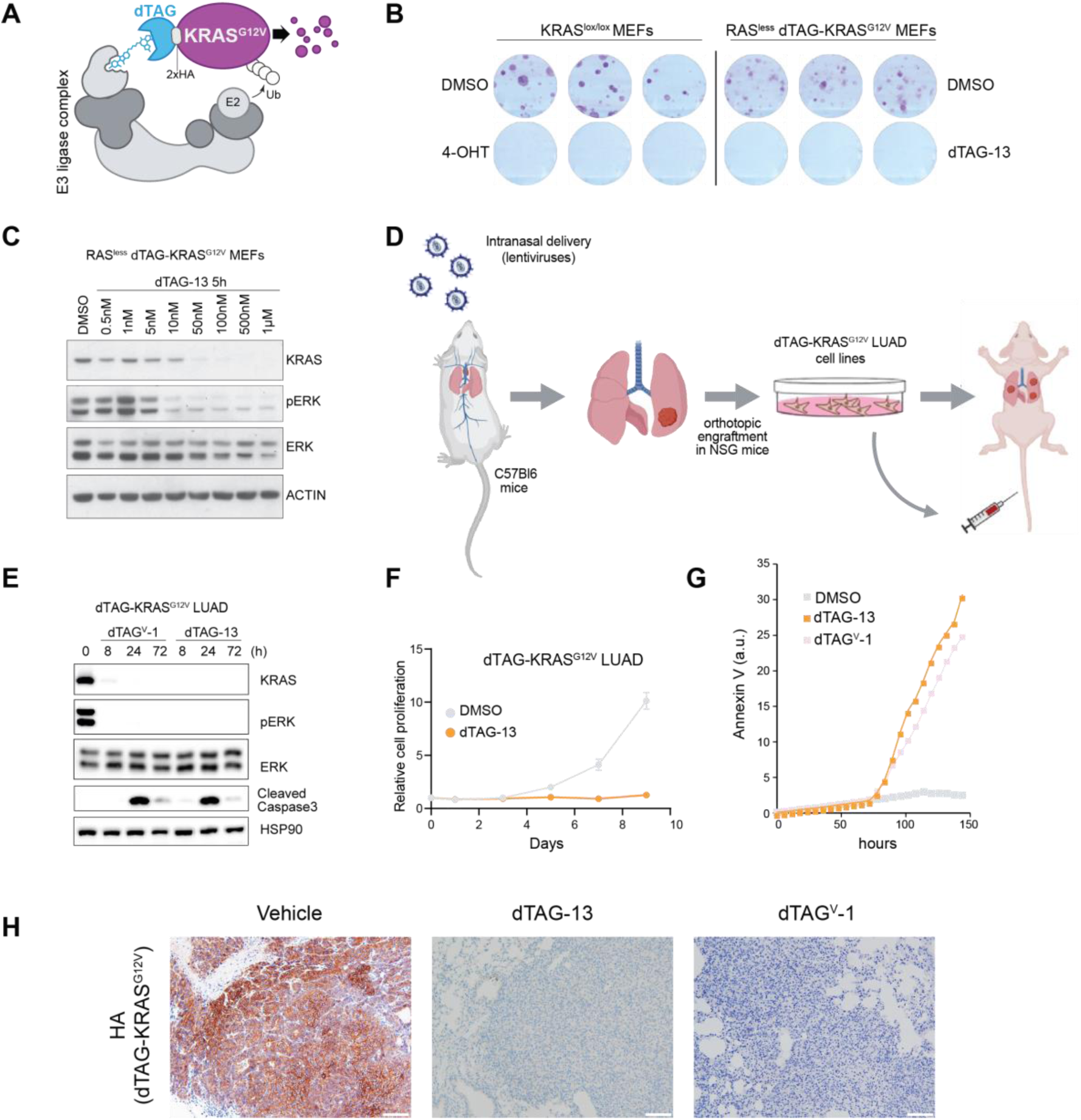
A novel LUAD mouse model driven by degradable KRAS^G12V^ enables PROTAC- dependent phenotyping *in vivo*. **A,** Depiction of the dTAG sytem. **B,** Colony forming assay of RASless MEFs without or with dTAG-KRAS^G12V^ expression. OHT (1 µg/mL); dTAG-13 = 0.3 µM. **C,** Degradation of dTAG-KRAS^G12V^ in RASless MEFs at the indicated doses and time. **D,** Depiction of the dTAG- KRAS^G12V^–driven LUAD model generation based on intranasal administration of lentiviruses carrying the dTAGged oncogene. **E,** Time-resolved degradation of dTAG-KRAS^G12V^ in LUAD cell lines established from tumors induced as depicted in (D). (dTAG-13 and dTAG^V^-1 = 0.25 µM). **F,** Growth curves of tumor-derived dTAG-KRAS^G12V^ LUAD lines (dTAG-13 = 0.25 µM). **G,** Time-resolved Annexin V staining upon degradation of dTAG-KRAS^G12V^ in LUAD cell lines (dTAG-13 and dTAG^V^-1 = 0.25 µM). **H,** Representative images of HA-stained (surrogate marker for dTAG-KRAS^G12V^) paraffin-embedded sections of LUADs from C57BL/6J mice treated with vehicle, dTAG-13, or dTAG^V^-1 for 7 days at 40 mg/Kg.

We focused on KRAS^G12V^, a variant with no selective inhibitors or PROTACs available. The C-terminal portion of KRAS contains the hypervariable region that cannot be modified as it is essential for KRAS trafficking and function. KRAS has been N-terminally fused to a variety of tags without any obvious biological alteration. Thus, we opted for the N-terminal dTAG fusion. The chimeric fusion also included two HA tags to facilitate detection (Fig. 1A). Of note, FKBP12^F36V^ and HA tags have been previously expressed *in vivo* with no immunogenicity reported^44–46^. First, we validated *in celullo* the ability of dTAG-KRAS^G12V^ to rescue the proliferation of cells devoid of all RAS isoforms^47,48^, thus confirming the biological activity of the tagged oncoprotein and the overall cellular efficiency of the system (Fig. 1B, C and Supplementary Fig. S1A).

To establish mouse models amenable to targeted KRAS degradation, we innovated a LUAD- induction approach based on intranasal inhalation of dTAG-KRAS^G12V^-carrying lentiviruses into fully immunocompetent C57BL/6J mice (Fig. 1D). Our approach confirmed the oncogenic activity of dTAG-KRAS^G12V^ by inducing the development of LUAD with an average latency of 10 months. Tumors were subsequently harvested and orthotopically implanted into the lungs of immunodeficient mice as this approach greatly enhances the development of primary cancer cell lines^49^. We next established several cancer cell lines derived from independent lesions and evaluated the efficiency of the dTAG system *in cellulo* upon exposure to the dTAG-13 and dTAG^V^-1 PROTACs. As expected, acute degradation of KRAS^G12V^ resulted in downstream signaling blockade accompanied by concomitant proliferation arrest and apoptosis (Fig. 1E- G).

Tail vein injection of the established cell lines consistently enabled the generation of syngeneic mouse models of LUAD. To test the efficacy of dTAG-KRAS^G12V^ degradation *in vivo* we first optimized the scalable synthesis of the compatible PROTACs dTAG-13 and dTAG^V^-1 (Supplementary Fig. S1B; see Methods and Supplementary information for detailed synthetic and purification procedures). In addition, we designed and synthesized an additional PROTAC analogue (IGP002; Supplementary Fig. S1B) aimed at improving the metabolic stability and solubility of the formers. However, we observed concomitant cellular degradation of the endogenous FKBP12 (Supplementary Fig. S1C). This finding with IGP1.002 was particularly surprising given that it only differs from dTAG-13 in one atom. We assume that subtle differences in the ternary complex formation drive the lack of FKBP12^F36V^-selective degradation. Henceforth, we focused on dTAG-13 and dTAG^V^-1, and observed efficient degradation *in vivo* by intraperitoneal administration at 40 mg kg^−1^ (Fig. 1H).

Collectively, we established an efficient approach to study tumor-specific chemical modulation of KRAS^G12V^ protein abundance in LUAD syngeneic mice.

### Degradation of oncogenic KRAS inhibits downstream signaling resulting in efficient tumor regression *in vivo*

To assess the *in vivo* potential of a KRAS^G12V^ degradation-based therapy, we further modified the dTAG-KRAS^G12V^ cell lines described above by concomitantly introducing a luciferase reporter and fluorescent marker (ZsGreen) to allow both longitudinal assessment of tumor burden and purification of cancer cells upon tissue dissociation, respectively. To this end, we delivered the dTAG-KRAS^G12V^ LUAD cells to the lungs of isogenic C57BL/6J recipient mice via caudal vein injection. This procedure resulted in efficient engraftment followed by tumor development with a short latency period (3-4 weeks). In order to induce degradation of KRAS^G12V^ we utilized dTAG-13 given than most of the PROTACs currently in clinical trials rewire the E3 CRL4^CRBN^ ^25^. Daily intraperitoneal injection of dTAG-13 significantly suppressed tumor growth inducing tumor regression as early as 3 days post-treatment, assessed both by luminescence signal (Fig. 2A) and histopathological examination (Fig. 2B). Two independent cohorts were sacrificed following 3 and 7 days of continuous dTAG-13 treatment. Immunohistochemical detection of the ZsGreen cancer cell reporter confirmed the efficient tumor regression phenotype anticipated by the decline in the luminescence signal. Likewise, quantification of HA-tag immunostaining (a KRAS^G12V^ surrogate marker) demonstrated the acute degradation of the driver oncoprotein (Fig. 2C). Furthermore, immunostaining analyses of lung tumor sections showed that dTAG-13 treatment resulted in a significant reduction of proliferative cells and enhanced induction of apoptosis when compared to untreated counterparts (Fig. 2C). As in the *in cellulo* data, oncoprotein degradation resulted in a marked downregulation of ERK/MAPK and PI3K signaling pathways (Fig. 2C). To gain deeper mechanistic insights, bulk RNA sequencing (RNAseq) was performed on LUAD cells (ZsGreen+/CD45– population) directly isolated upon dissociation and FACS separation of tumor-bearing lungs. The biological significance of KRAS^G12V^ degradation was reinforced by an overall transcriptional profile almost identical to a recently reported gene-signature obtained following chemical inhibition of the oncogene^50^ (Fig. 2D). A more detailed scrutiny using gene set enrichment analysis (GSEA) identified 7 gene sets significantly enriched in dTAG-13- treated tumor cells, including immune-related categories such as those associated with type I and type II interferon (IFN) alpha and gamma responses (Fig. 2E), previously reported in the context of oncogenic KRAS inhibition^21^. Consistent with this, we observed increased expression of IFN target genes (Fig. 2F). On the other hand, 12 gene sets were found significantly downregulated when compared to untreated tumor cells with a remarkable presence of metabolic and proliferative related functions (Fig. 2E). Collectively these data indicate that targeted degradation of KRAS^G12V^ is a highly effective and rapid anti-tumoral approach.

**Figure 2.**
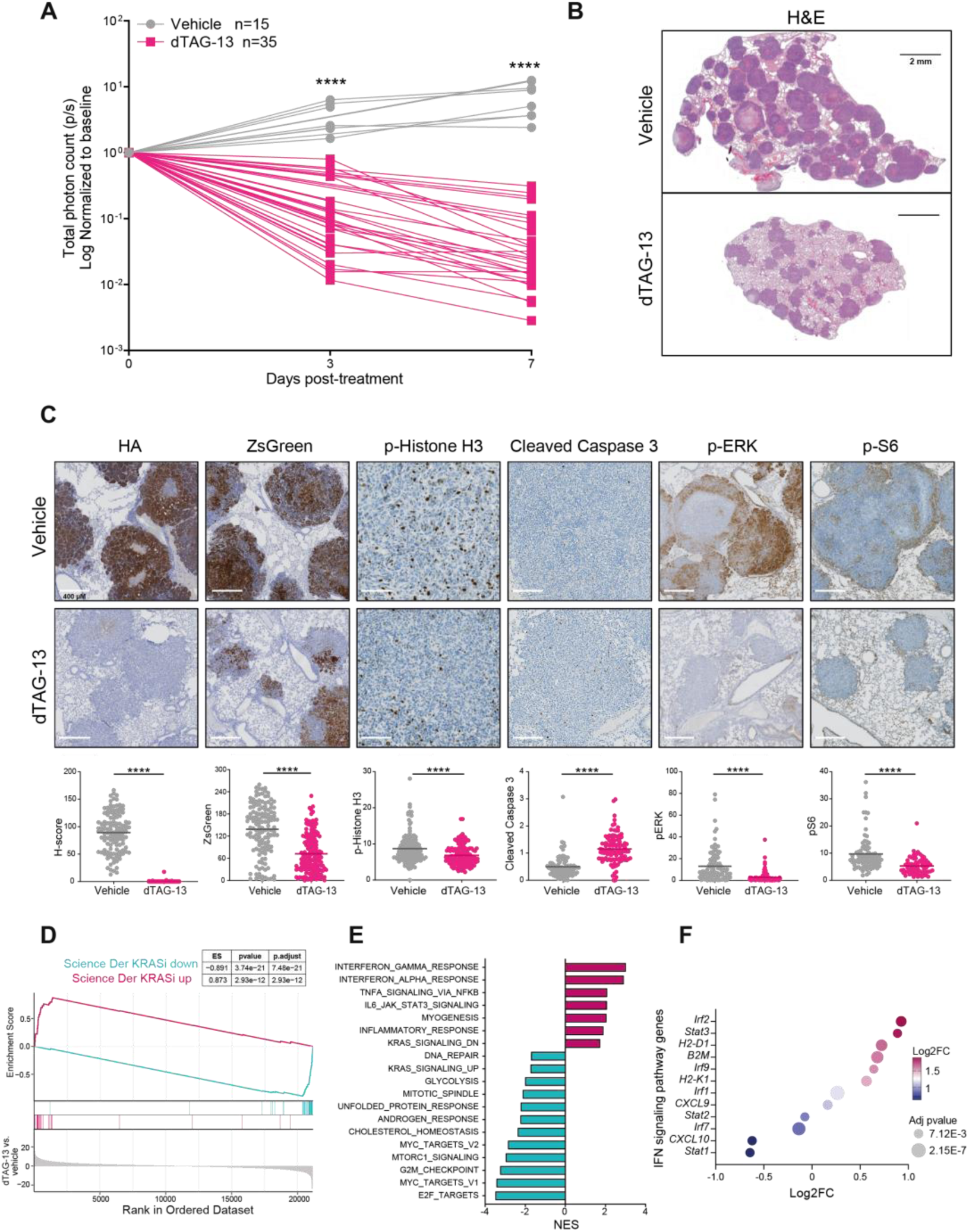
KRAS^G12V^ targeted degradation induces tumor regression associated with changes in cell-autonomous and non-autonomous processes. **A,** Longitudinal quantification of LUAD tumor evolution by IVIS imaging (LUAD bioluminescence signal) in C57BL/6J mice treated daily with vehicle or dTAG-13 (40 mg/kg). Measurements were performed at days 3 and 7 post-treatment initiation. **B,** Representative images of hematoxylin and eosin (H&E)-stained, paraffin-embedded lung sections of tumor-bearing C57BL/6J mice treated daily with vehicle or dTAG-13 (40 mg/kg, day 7). **C,** Representative images of the indicated immunohistochemistry markers (upper panels) and the quantification (bottom graphs) from tumors of C57BL/6J mice treated with vehicle or dTAG-13 (40 mg/kg). **D,** Gene signature associated with KRAS inhibition with enrichment scores among the corresponding up and downregulated genes differentially expressed in dTAG-13 vs. vehicle-treated FACS-sorted LUAD cells (ZsGreen+/CD45-). **E,** Hallmark gene set enrichment analysis (GSEA) with positive and negative enrichment scores from RNAseq data of 7-day dTAG-13 vs. vehicle treated FACS-sorted LUAD cells. **F,** Dysregulated canonical IFN signaling pathway genes upon KRAS^G12V^ degradation. Statistical differences in A, B and C were analyzed using Mann-Whitney test. ****, p < 0.0001. FDR Q values in D, E and F were calculated using GSEA software.

### Role of the adaptive immunity in the immediate therapeutic response triggered by KRAS^G12V^ degradation

The results presented above, highlighting the increased enrichment of several pro-inflammatory pathways following KRAS oncoprotein degradation, prompted us to investigate the tumor microenvironment (TME) composition. Comprehensive immunophenotypic characterization of both dTAG-13 treated and untreated tumor-bearing lungs, revealed that the tumor shrinkage observed as soon as 3 days after the degrader administration occurred concomitantly with an expansion of immune (CD45+) and, to a lesser degree, non-immune (CD45–) cell components (Fig. 3A). To more precisely quantify the immune-related phenotypic changes, all immune cell subtypes were normalized relative to the tumor cellularity (ZsGreen+/CD45– population). As shown in Fig. 3B, the elimination of KRAS oncoprotein significantly increased both myeloid and lymphoid major lineages. Since cytotoxic populations, such as CD8+ T cells, are pivotal in mediating immunological rejection, we focused our initial analyses on T cell compartments. Most notably, the relative abundance of all major T cell subtypes was significantly increased upon oncogenic KRAS^G12V^ degradation, with a concomitant decrease of the respective peripheral blood populations, likely indicative of a strong pulmonary recruitment (Fig. 3C and Supplementary Fig. S2A). Indeed, immunostaining of lung sections revealed enhanced infiltration of CD8+ T cells in dTAG-13-treated tumors (Fig. 3D), a process potentially linked to the upregulation of CXCL9, CXCL10, and CXCL11 chemokines and interferon-stimulated genes upon KRAS degradation (Fig. 2F). Adaptive immunity, driven by priming and activation of T cells, critically depends on dendritic cells (DCs), which phagocyte and present antigens loading them on plasma membrane embedded MHC proteins as an initial step. In response to KRAS^G12V^ degradation, increased infiltration of the two major DC subtypes (cDC2 and CD103+ DC cells) was detected (Fig. 3E) alongside with reduced overall numbers in peripheral blood (Supplementary Fig. S2B), again potentially indicating increased DC recruitment to tumor sites. Among them, the CD103+ DC subset displaying co-expression of the co-stimulatory molecule CD80 showed a more remarkable expansion, suggesting a more effective antigen cross-presentation to CD8+ cells. Accordingly, the activated cytotoxic CD8+CD4- T cell subset appeared significantly amplified according to the CD25 activation marker (Fig. 3F). Likewise, the activated CD4+_Th subtypes also displayed a similar amplification pattern (Fig. 3F), probably due to the increased DC2 infiltration and the MHC-II presentation capacity of both cDC2 and CD103+ DCs. Moreover, KRAS^G12V^ degradation also promoted the acquisition of a differentiation phenotype with both memory and effector properties (Fig. 3G). This robust adaptive cytotoxic response was potentially enhanced both by increased MHC-I expression of LUAD cells (Fig. 2F), leading to higher antigen presentation capacity, as well as by elevated numbers of innate cytotoxic NK cells (Fig. 3B). Altogether, these evidences highlight the prominent role played by oncogenic KRAS in the suppression of several anti-tumor immune responses that appeared significantly increased following degrader treatment.

**Figure 3.**
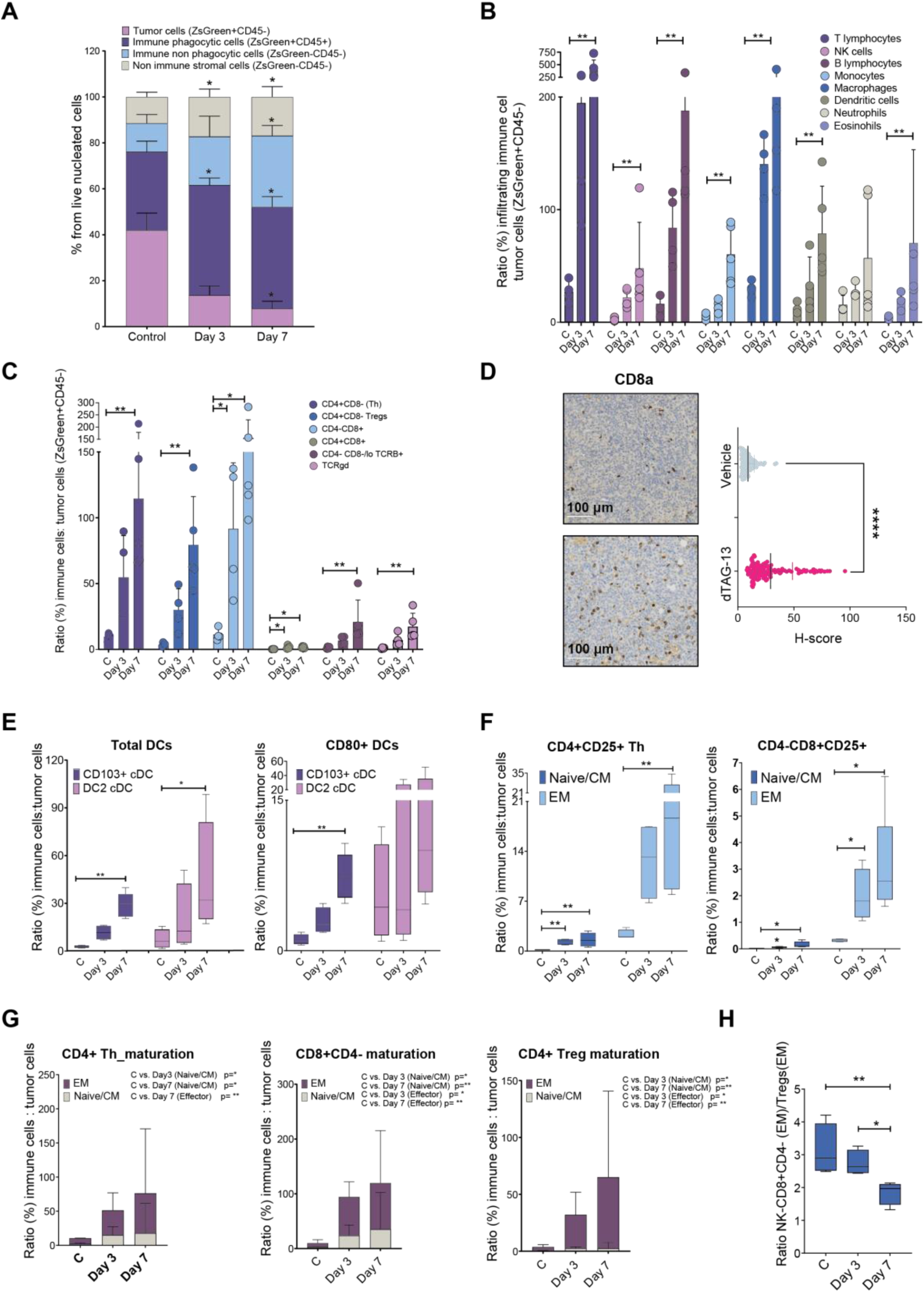
Adaptive immune phenotypes driven by KRAS^G12V^ degradation. **A,** Immune and non-immune cell population summary from the flow cytometric analysis following lung tissue dissociation. Percentage of live cells for vehicle- (n = 4) and dTAG-13-treated (40 mg/kg) C57BL/6J mice at 3 (n = 4) or 7 (n = 5) days post-treatment initiation. **B,** Normalized abundance of the indicated immune cell populations with respect to the remaining tumor cells in each of the treatment conditions (vehicle and dTAG-13 treatment). **C,** Relative abundance, considering both lung immune infiltration and tumor cellularity, of the indicated major T cell subtypes in response to vehicle or dTAG-13 treatment. **D,** Representative immunostainings of tumor-bearing lung sections showing CD8+ T cell infiltration levels in lung tumors of 7 day treated and untreated mice (left). Quantification of CD8+ cells in all tumors identified in both vehicle and dTAG-13-treated cohorts is depicted (right graph). **E, F,** Quantitative analysis as in (B) and (C) of total and CD80+ dendritic cells (DCs) (E) and evaluation of T cell activation (CD25+) of CD4+Th and CD8+CD4- T cells (F) in vehicle, 3 day- and 7 day-dTAG-13 treated C57BL/6J mice. **G,** Maturation stage distribution including Naïve/Central Memory (CM) and Effector Memory (EM), is shown for the 3 major T cell populations at each of the indicated time points in vehicle or dTAG-13 treated mice. **H,** Evaluation of the combined contribution of cytotoxic cell populations (mature NK and CD8+CD4- T cells) compared to immunosuppressive regulatory T cell (Tregs) infiltrates in vehicle and dTAG-13 treated cells at 3 (n = 4) or 7 (n = 5) days post-treatment initiation. Statistical differences were analyzed using non-parametric Mann-Whitney test in (D) and non-parametric One-way ANOVA followed by False Discovery Rate multiple comparison tests in the rest of the panels. In panel (A) all the statistical differences correspond to the comparisons between treated and vehicle control conditions. *, p< 0.05; **, 0.05<p<0.01; ****, p < 0.0001. Data are indicated as the mean ± sd.

Finally, the ratio between the sum of cytotoxic lymphoid CD8+CD4- and NK cells and the counteracting immunosuppressive Treg population, was found significantly diminished at day 7, suggesting a shift towards inflammation resolution likely due to the drastic reduction of the tumor burden during dTAG-13 treatment (Fig. 3H).

### KRAS^G12V^ degradation enhances tumor regression mechanisms potentially dependent on the phagocytic ability of the myeloid innate immunity

Next, we evaluated the role of the innate immune response as a potential contributor in the tumor regression process described above. To determine the importance of both the adaptive and innate immune components in the short-term antineoplastic response, we replicated the orthotopic engraftment procedure using NSG mice, a strain characterized by lacking functional B, T and NK cells. Remarkably, tumor regression (measured by the decline in bioluminescence signal) occurred with similar kinetics to that observed in an immunocompetent background (Fig. 4A). Accordingly, histological examination showed a significant decrease in tumor area as well as in the amount of HA and ZsGreen positive LUAD cells upon dTAG-13 administration (Supplementary Fig. S3A and 3B). Likewise, immunohistochemical analysis revealed an impairment of ERK/MAPK and PI3K signaling pathways accompanied by cleaved caspase 3 induction and compromised cell proliferation (Supplementary Fig. S3B). These anti-tumor phenotypes, also observed upon dTAG13 treatment in the C57BL/6J background, are likely the consequence of tumor-intrinsic processes and are in good agreement with the substantial GSEA intersection observed between both genetic backgrounds (Supplementary Fig S3C).

**Figure. 4.**
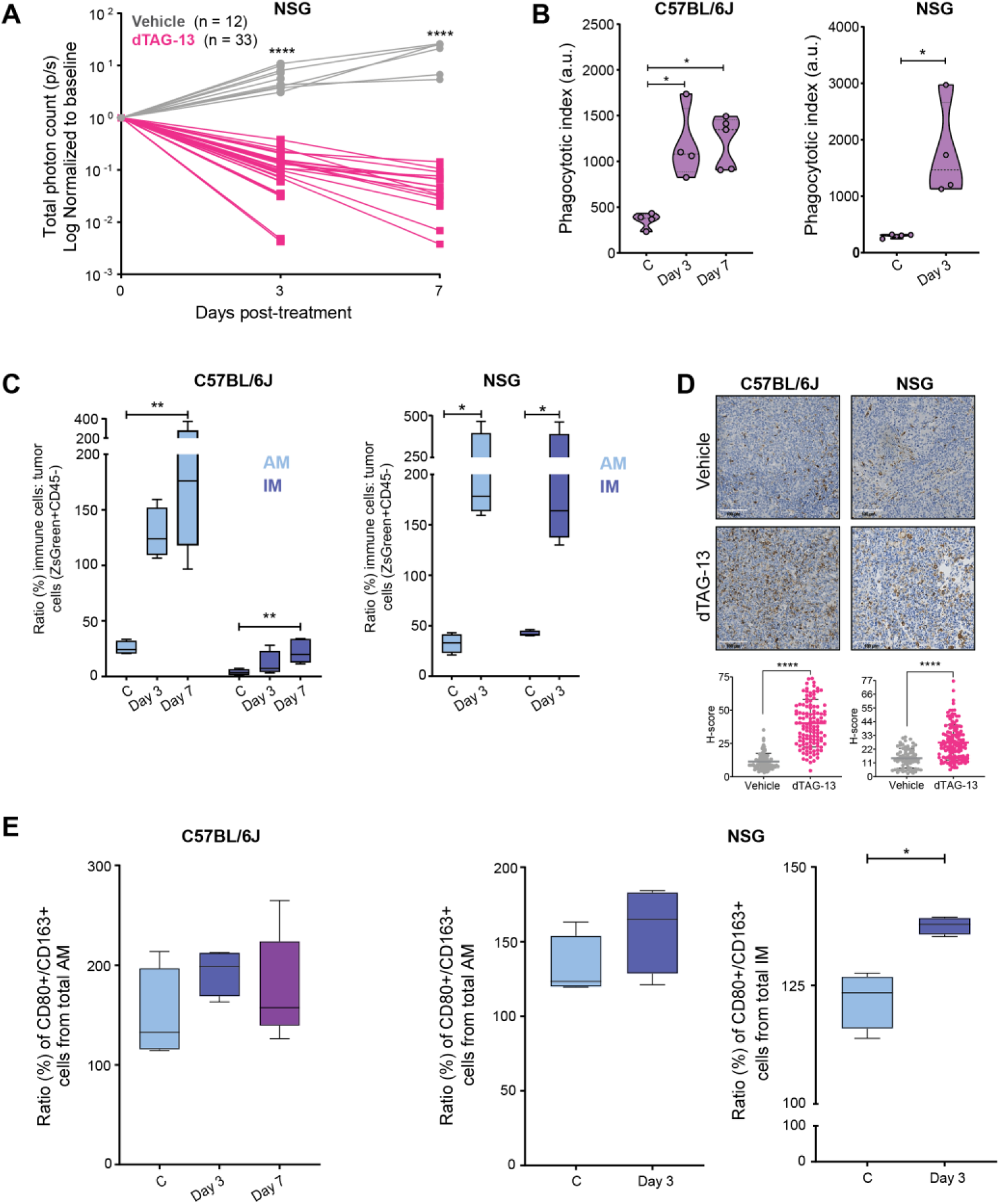
Prominent anti-tumor response persists despite the absence of a functional adaptive immune system following oncogenic KRAS^G12V^ protein degradation. **A,** Longitudinal quantification of LUAD tumor evolution by IVIS imaging (LUAD bioluminescence signal) in NSG mice treated daily with vehicle or dTAG-13 (40 mg/kg) at days 3 and 7 post-treatment initiation. **B,** Comparative analysis (vehicle vs. dTAG-13 treatment) in NSG and C57BL/6J mice of the respective phagocytic indexes (see Methods). **C,** Relative abundance of AM (alveolar macrophages) and IM (interstitial macrophages) upon vehicle or dTAG-13 treatment in NSG and C57BL/6J mice. **D,** Representative images of lung tumor macrophage infiltration (F4/80 pan-macrophage marker) from NSG and C57BL/6J cohorts (upper). Immunohistochemical scoring (H-score) values for F4/80 expression in all tumor bearing samples obtained from C57BL/6J and NSG mice and subjected to the indicated treatment conditions (bottom). **E,** Representation of the pro-/anti-inflammatory macrophage ratio based on the expression of the respective polarization markers CD80 and CD163 is shown for NSG and C57BL/6J cohorts upon vehicle or dTAG-13 treatment (40 mg/kg). Statistical differences were analyzed using non-parametric Mann-Whitney test and non-parametric One-way ANOVA followed by False Discovery Rate multiple comparison tests, in NSG and C57BL/6J mice, respectively. *, p< 0.05; **, 0.05<p<0.01; ****, p < 0.0001. Data are indicated as the mean ± sd.

Next, we decided to directly address the potential implication of innate immune responses in both mouse strains. Considering the key role of the phagocytic response as an anti-tumor mechanism^51^, we took again advantage of the ZsGreen reporter expressed by tumor cells to perform a more precise characterization of this cellular response. Noteworthy, degradation of KRAS oncoprotein greatly exacerbated the overall phagocytic activity in both immunocompetent and immunocompromised tumor bearing lungs (Fig. 4B). Among the potential immune populations with phagocytic capacity, alveolar macrophages (AM) emerged as the most prominent cell type able to phagocyte tumor cells in the C57BL/6J background. In NSG mice, interstitial macrophages (IM) were also actively implicated in this process (Fig. 4C). In good agreement, we detected massive infiltration in both models as measured by the F4/80 pan-macrophage marker in dTAG-13 treated tumors (Fig. 4D). Interestingly, both AM and IM populations displayed an increased proportion of CD80+ cells indicative of enhanced anti-tumoral M1 phenotype (Fig. 4E).

All these findings suggest that the combination of cancer-cell intrinsic together with innate anti-tumor immune responses contribute to the tumor regression phenotype observed at early stages following KRAS^G12V^ degradation.

### Resistance to KRAS^G12V^ degradation *in vivo* is mostly driven by dysregulation of the proteolytic machinery

Despite the rapid and efficient tumor regression elicited by KRAS^G12V^ degradation, we decided to prolong the dTAG-13 treatments *in vivo* to evaluate the emergence of resistance to this therapeutic approach. Patterns of PROTAC resistance *in vivo* remain largely unknown. We therefore followed C57BL/6J and NSG cohorts until detecting tumor relapse (see Methods). Mice were subsequently maintained on dTAG-13 treatment until reaching humane end-point to increase the selective pressure potentially maximizing the onset of full therapy resistance. dTAG-13 resistance appeared with accelerated kinetics in NSG mice resulting in shorter median time to progression and overall reduced survival when compared to C57BL/6J mice (Fig. 5A-D). Immunohistochemical studies of the resistant tumors revealed that the vast majority of proliferative lesions retained KRAS^G12V^ protein expression (Fig. 5E, F).

**Figure 5.**
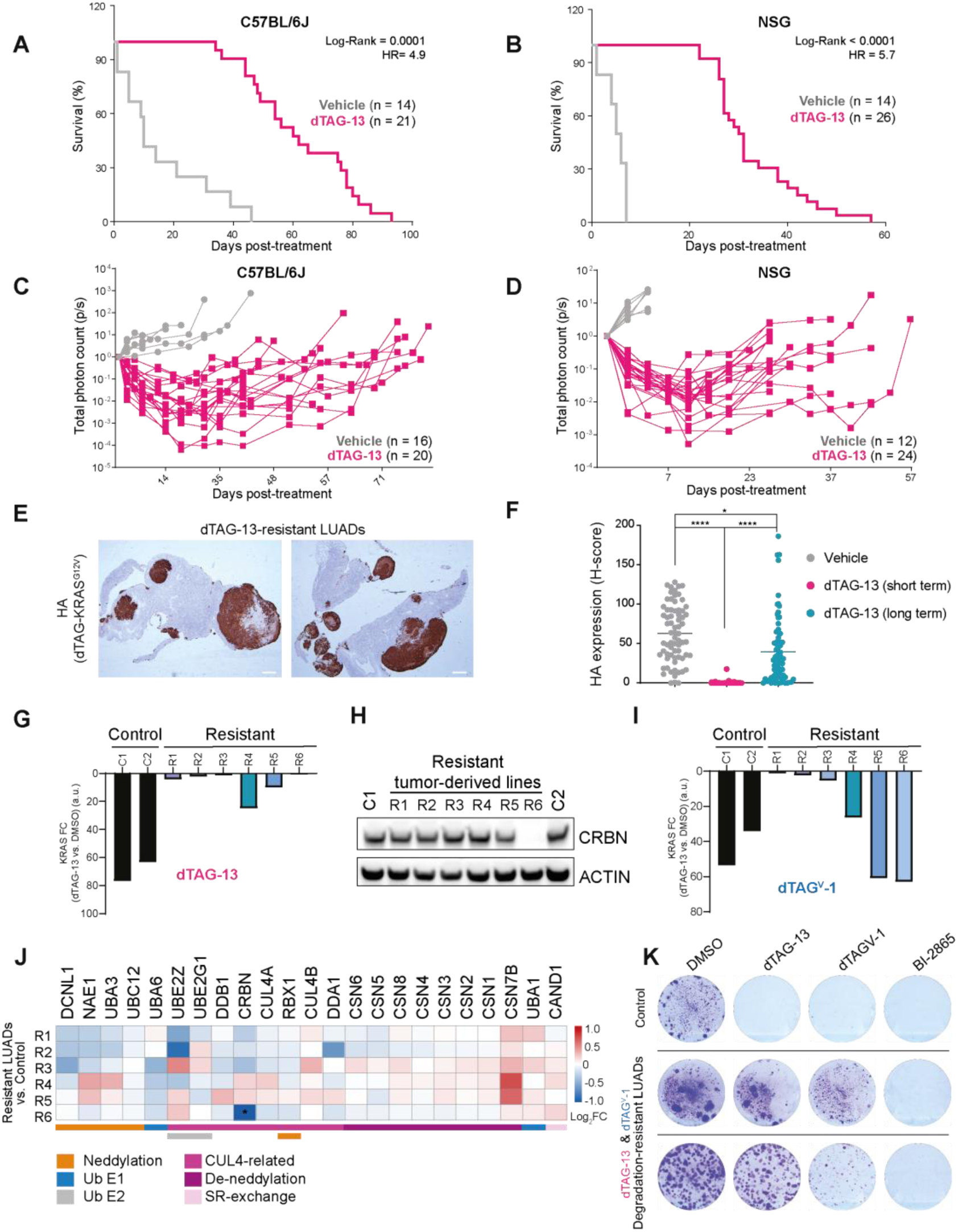
Resistance to KRAS degradation *in vivo* is mainly driven by dysregulation of the proteolytic machinery. **A-D,** Longitudinal assessment of dTAG-KRAS^G12V^-driven LUAD tumor evolution by IVIS imaging (LUAD bioluminescence signal in C57BL/6J (C) or NSG (D) mice treated with vehicle or dTAG-13 at 40 mg/kg 4/days/week. The log rank test was used to evaluate survival differences between both groups, and the HR indicates an increased risk of LUAD related-death in the untreated group. **E,** Representative images of HA (dTAG-KRAS^G12V^) paraffin-embedded sections of dTAG-13- resistant LUADs from C57BL/6J mice treated with dTAG-13 (40 mg/kg) 4/days/week until humane end-point. **F,** Quantification of HA (dTAG-KRAS^G12V^) expression from paraffin-embedded sections of LUADs from C57BL/6J mice treated with vehicle, or dTAG-13 (short treatment = 7 days; long treatment ≥ 45 days). **G,** Quantification of dTAG-KRAS^G12V^ degradation (dTAG-13 vs. DMSO) in LUAD murine cells (dTAG-13: 0.5 µM, 24h) from western blot in Supplementary Fig. S4C. Cell lines established from treatment-naïve and dTAG-13 resistant LUAD tumors. **H,** CRBN levels in the cell lines shown in (D). **I,** Quantification of dTAG-KRAS^G12V^ degradation (dTAG^V^-1 vs. DMSO) in LUAD murine cells (dTAG^V^-1: 0.5 µM, 24h from western blot in Supplementary Fig. S4D). **J,** Targeted proteomics via parallel reaction monitoring (PRM) of a selection of E3-related proteins. Log2FC (resistant LUADs vs. control) is shown in the heatmap (n=3). Functional information is color-coded as indicated in the legend. Ub: ubiquitin; SR: substrate receptor. *CRBN in sample R6: lower than minimum color-coded scale. **K,** Colony forming assays in LUAD lines resistant to dTAG-13 and dTAG^V^-1 (I). dTAG-13 = 0.3 µM; BI-2865 (panKRAS inhibitor) = 8 µM; dTAG^V^-1 = 0.3 µM.

Cell lines were established from resistant tumors for further phenotyping, together with control lines derived from treatment-naïve animals. Time-resolved immunoblotting confirmed that dTAG-13-driven KRAS^G12V^ degradation was compromised in the resistant lines compared to control cells (Fig. 5G and Supplementary Fig. S4C). We then focused on understanding the mechanisms driving the lack of targeted degradation. To discard potential technical issues related to the dTAG system, we first sequenced the dTAG-KRAS^G12V^ fusion in the cell lines established from the resistant LUAD tumors and revealed no on-target secondary mutations (Supplementary Table 1). Efflux pump overexpression has been reported as a mechanism of resistance to PROTACs^52^. However, these models were sensitive to paclitaxel, a common substrate of efflux pumps^53^, and an efflux pump inhibitor did not restore dTAG-13-induced degradation of KRAS (Supplementary Fig. S4A,B).

The loss of function of the E3 hijacked by degraders is the most common resistance mechanism *in cellulo*^35,54–58^. To our surprise, only one of the tumor-resistant cell lines showed lack of CRBN expression (Fig. 5H). We treated our dTAG-13 resistant models with dTAG^V^-1, which recruits CRL2^VHL^ instead of CRL4^CRBN^. In addition to the cell line without CRBN, some others were sensitive to this alternative PROTAC (Fig. 5I and Supplementary Fig. S4D), suggesting that the lack of KRAS degradation was driven by factors related to the CRL4^CRBN^ complex. For instance, we observed some downregulation of the adaptor protein DDB1 in some clones (Supplementary Fig. S4E), previously shown to confer resistance to degraders *in cellulo*^35,59^. When challenged with other KRAS-unrelated PROTACs that co-opt the same E3s, similar compromised targeted degradation was observed (Supplementary Fig. S4F). These results suggested dysregulation of proteolytic machinery upstream or downstream of the E3 ligases as the main drivers of resistance *in vivo*. To query changes in the protein plasticity of E3-related factors, we defined a panel of interesting proteins related to the Cullin-RING ligase (CRL) family of ubiquitin E3s, and performed targeted proteomics via parallel reaction monitoring (PRM). We could capture a trend in the tumor-derived resistant cell lines towards dampening the levels of important activation machinery (Fig. 5J). For instance, downregulation of DCNL1 and NAE1, both part of the neddylation cascade to activate CRLs, was observed and might contribute to the defective targeted degradation of KRAS (Fig. 5J). We also mined our RNAseq data to search for potential coding point mutations that might compromise ligase activity in this panel of E3-related genes. In addition to the resistant cell lines, control and dTAG-13 resistant tumors from immunocompetent and immunodeficient mice were analyzed. We only found one point mutation (50% of reads, which would be equivalent to a heterozygote tumor) corresponding to a single-nucleotide insertion predicted to cause a frameshift in the *Nae1* gene (Supplementary Fig. S4G and Supplementary Table 2). Overall, we could not find any other point mutations predicted to affect E3 activity in the resistant cell lines or tumors, suggesting the observed dysregulation at the protein level as the most probable contribution towards resistance (Fig. 5J).

Given that the KRAS oncoprotein was expressed, we decided to test whether our degradation-resistant LUAD models were sensitive to KRAS inhibitors. There are no selective KRAS^G12V^ inhibitors available, but the panKRAS inhibitor BI-2865 engages this mutant. dTAG-13 resistant cells and the cell lines also resistant to dTAG^V^-1 were sensitive to BI-2865 (Fig. 5K) reinforcing their dependency on the driver oncoprotein. These results support the rational use of different, orthogonal pharmacology against the same target to defeat resistance.

In conclusion, impaired KRAS^G12V^ degradation was the predominant resistance pattern observed *in vivo*. Mechanistically, our studies suggest that resistance *in vivo* mainly occurs as a consequence of dysfunctional E3-dependent protein degradation, a mechanism that can be overcome with KRAS chemical inhibitors.

## DISCUSSION

Given the limited duration of the clinical responses to FDA-approved KRAS inhibitors, finding alternative therapies is a pressing medical need. In this context, the development of KRAS PROTACs, with some already under early clinical evaluation, could represent a significant breakthrough in the treatment of KRAS-driven cancers. Yet, there is currently very limited information regarding the mechanistic phenotypes brought about by oncogenic KRAS targeted degradation *in vivo* and no data whatsoever concerning the potential resistance mechanisms.

In this study, we focused on modeling targeted degradation of KRAS^G12V^ in LUAD using the dTAG system. Leveraging this PROTAC-based chemogenetic strategy, we enabled *in vivo* mechanistic and follow-up studies that could inform the efficacy of the first recently reported pan-KRAS degraders^33^. Overall, our preclinical findings provide proof-of-principle of KRAS^G12V^ degradation as a therapeutic strategy in LUAD, potentially leading to durable anti-tumor responses through a combination of cell-intrinsic and extrinsic mechanisms. The transcriptional profile obtained from LUAD tumors following KRAS^G12V^ degradation displayed substantial overlapping with recently reported signatures upon oncogene inhibition using small molecules^50^, suggesting that dampened downstream and pro-survival signaling are major contributors leading to early tumor regression.

KRAS-derived neo-peptides have been identified as efficient immunological targets^60^. Peptides derived from targeted proteasomal degradation display an average size compatible with MHC-mediated presentation. It is therefore tempting to speculate that, in addition to the enhanced inflammatory response driven by the abrogation of KRAS-driven immune evasion programs, acting on both innate and adaptive compartments as previously reported upon inhibition with specific inhibitors^61^, PROTACs directed against KRAS may also fuel anti-tumor immunity by means of neo-antigens directly derived from its targeted degradation. While some of these notions will require additional experimental support, altogether our observations suggest a potential synergy between KRAS degradation and tailored immunotherapy to achieve more durable responses.

Currently, the patterns of PROTAC resistance *in vivo* remain largely unknown. However, potential resistance mechanisms can be inferred from studies with cell lines. We and others have mapped through functional genomic studies how the repertoire of E3 ligases and their regulators shape the resistance mechanisms to targeted protein degradation^35,54–56,62^. Indeed, these mechanisms largely differ from the target-centric resistance associated with traditional inhibitors, and mainly involve the loss or mutation of key ubiquitin-proteasome-system components such as E3 ligases, E2 enzymes, or other regulatory factors^35,62^. Given that several PROTACs and molecular glue degraders are currently undergoing clinical trials, valuable lessons in these clinical settings are expected to complement our current understanding of how cancer cells cope with degrader modalities.

Our *in vivo* studies revealed that long-term treatment eventually resulted in some disease relapse due mainly to KRAS-independent mechanisms. We observed that tumor progression occurred with delayed dynamics in immunocompetent hosts, suggesting that effector/memory components of the adaptive immunity might be key to enhance therapeutic responses. Interestingly, resistance predominantly emerged from dysfunctional E3-dependent protein degradation, and not from adaptive or acquired KRAS-related mechanisms as observed in patients progressing on KRAS specific inhibitors. Our preclinical study is based on tag-based PROTACs, thus, direct KRAS degraders might have divergent resistance mechanisms and/or toxicity profiles. Nevertheless, we hypothesize that the resistance patterns we observed with dTAG-13 are expected to be similar for PROTACs currently in clinical trials against different targets, given that most of them also hijack the E3 CRL4^CRBN^ via similar thalidomide-inspired ligands. Thus, our KRAS-focused work might provide valuable insights into potential therapeutic challenges of PROTACs as a pharmacological modality. Our findings suggest a valuable opportunity to explore on-target combination therapies with different pharmacological modalities. Indeed, the use of a KRAS inhibitor was able to overcome degrader resistance *in cellulo*, suggesting that the sequential use of PROTACs and inhibitors could become a promising strategy to achieve more sustained therapeutic responses in KRAS-driven LUAD. Ongoing research with engineered mouse models will further elucidate the systemic effects triggered by KRAS degradation and help refine therapeutic strategies.

## Supporting information

Supplementary Material Chemistry

## SUPPLEMENTARY FIGURES

**Supplementary Figure S1.**
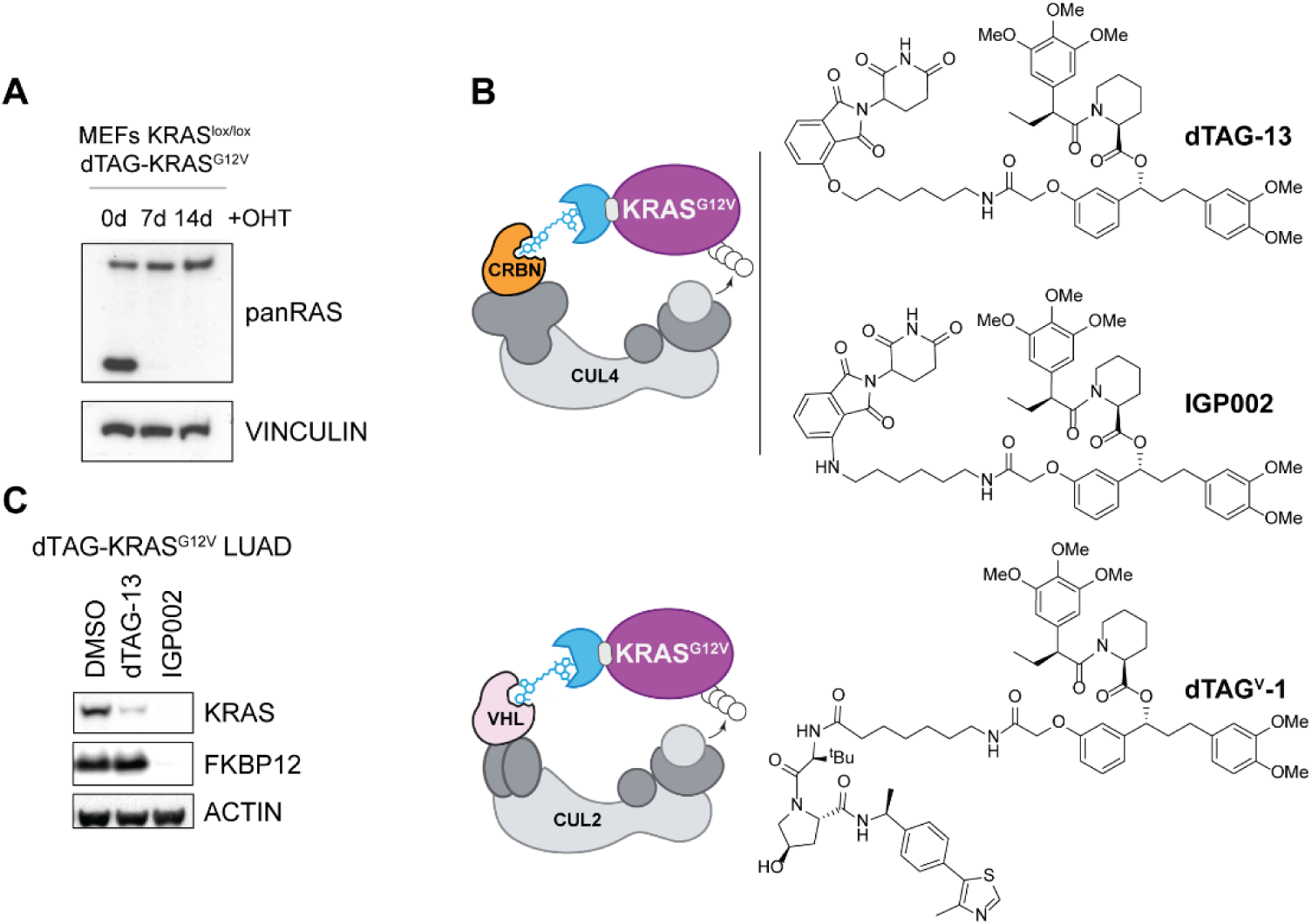
Selection of PROTACs compatible with the dTAG-KRAS^G12V^ LUAD model. **A,** RASless MEFs with dTAG-KRAS^G12V^ expression generated from H/NRAS^-/-^ KRAS^lox/lox^ MEFs. (OHT: 1 µg/mL). **B,** Structures of the dTAG PROTACs tested in (C) and depictions of the E3 recruited. IGP002 was synthesized aiming to improve the metabolic stability and solubility of the previously reported PROTAC dTAG-13. **C,** Degradation of dTAG-KRAS^G12V^ and endogenous FKBP12 in LUAD murine cells with DMSO or the indicated PROTACs (0.3 µM, 24h).

**Supplementary Figure S2.**
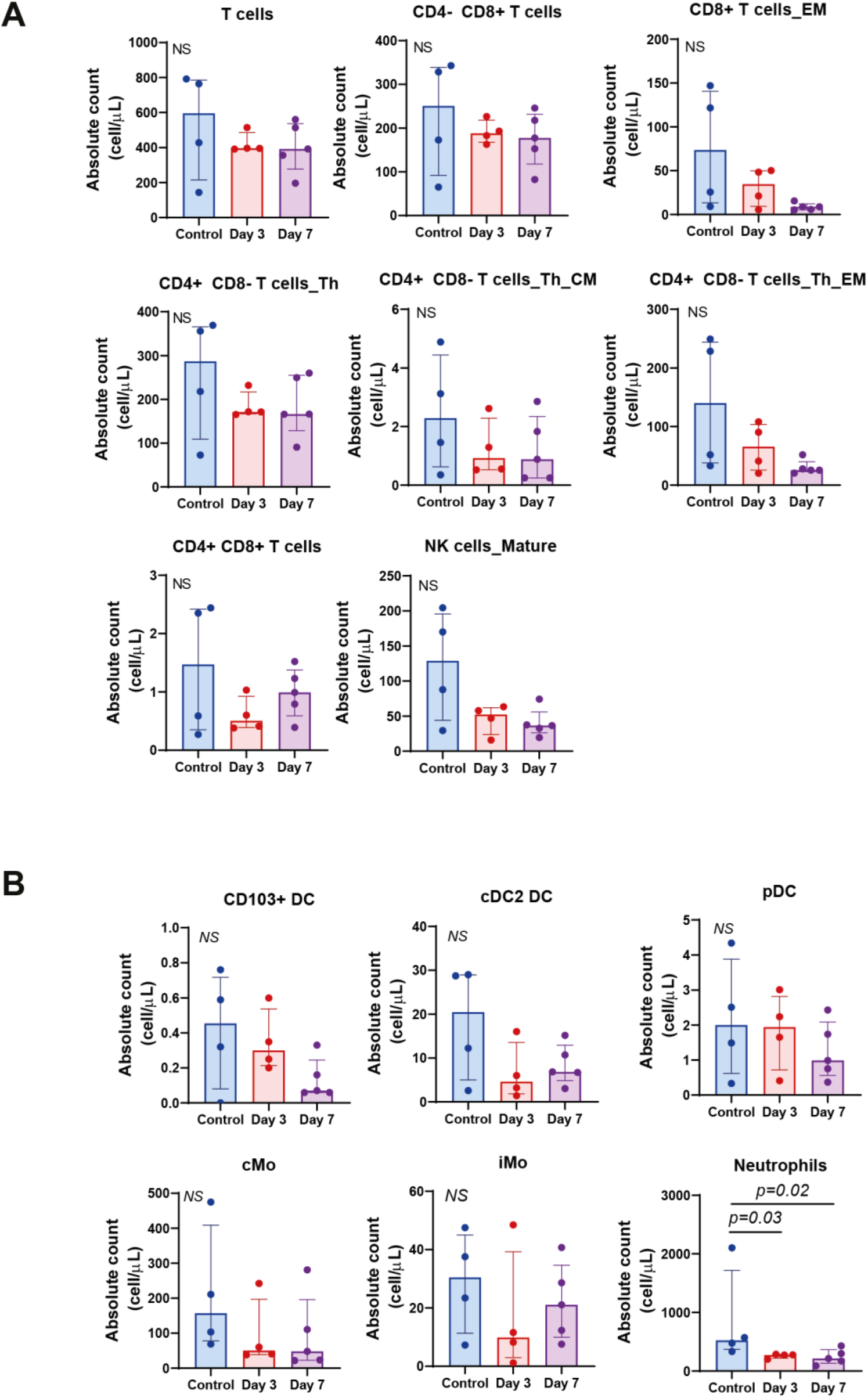
Reduced peripheral blood cell counts suggest that dTAG-13 treatment induces recruitment of immune cells to LUAD tumors. **A,B,** Quantification of peripheral blood immune cell populations at 3 and 7 days post dTAG-13 treatment initiation. Lymphoid cell types are shown in (A) and myeloid types in (B).

**Supplementary Figure S3.**
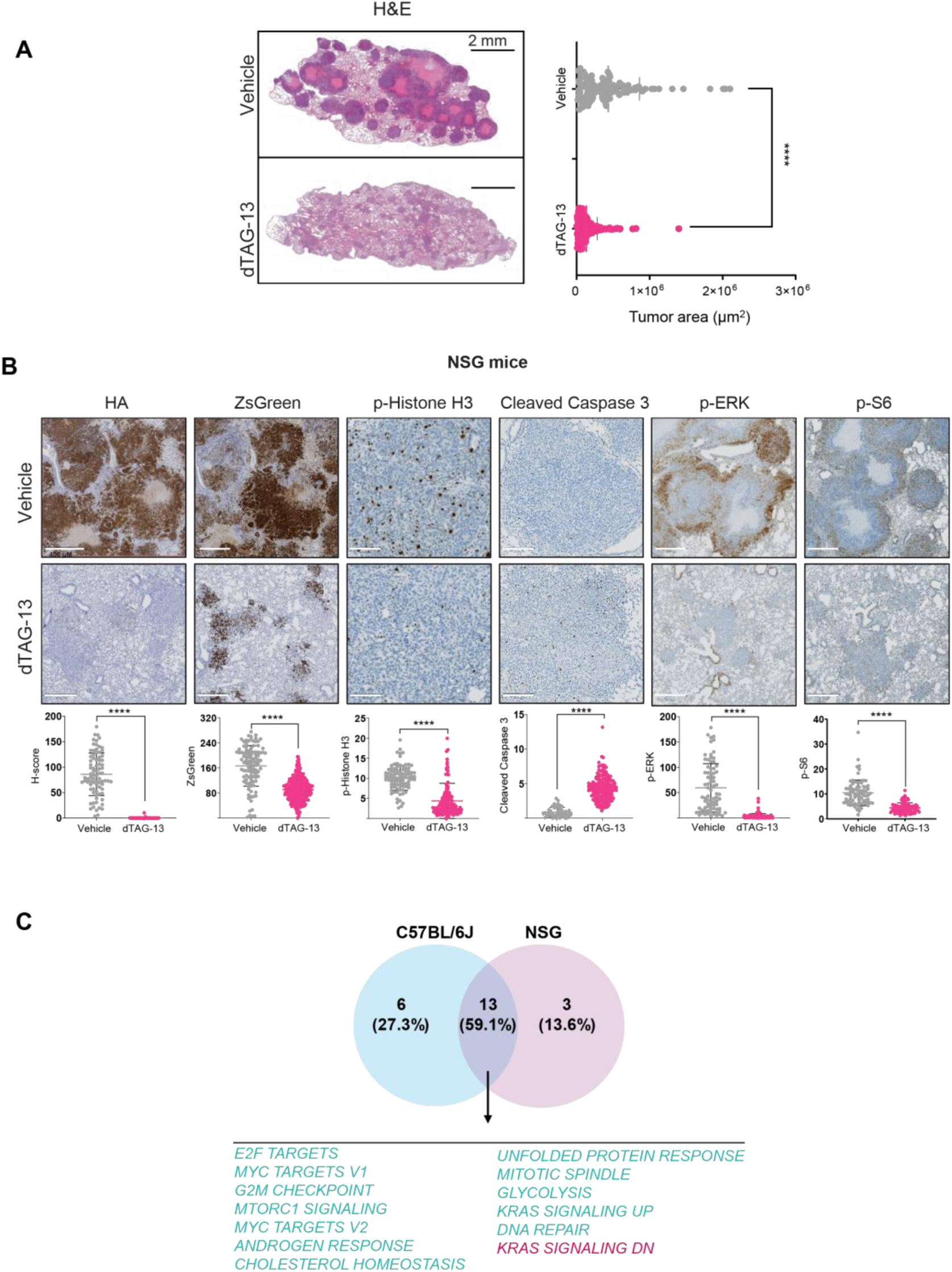
The anti-tumor response elicited by KRAS^G12V^ degradation in immunocompromised NSG mice closely resembles the effects observed in immunocompetent C57BL/6J mice. **A,** Representative example of H&E- stained sections of tumor-bearing lungs collected from C57BL/6J and NSG mice together with the respective tumor area quantifications (right). **B,** Representative images of the indicated immunohistochemistry markers (upper panels) and the quantification (bottom graphs) from tumors of NSG mice treated with vehicle or dTAG-13 (40 mg/kg) at day 7 post treatment initiation. Quantification data correspond to individual lung tumors derived from all the animals included in each of the mouse cohorts (vehicle and dTAG-13 treated). Statistical differences were analyzed using non-parametric Mann-Whitney test. ****, p < 0.0001. **C,** Positively (red) and negatively-enriched (green) biological processes shared in LUAD cells from dTAG-13- treated tumor-bearing mice (C57BL/6J and NSG). GSEAs of RNAseq data from the tumor-derived LUAD cells were applied.

**Supplementary Figure S4.**
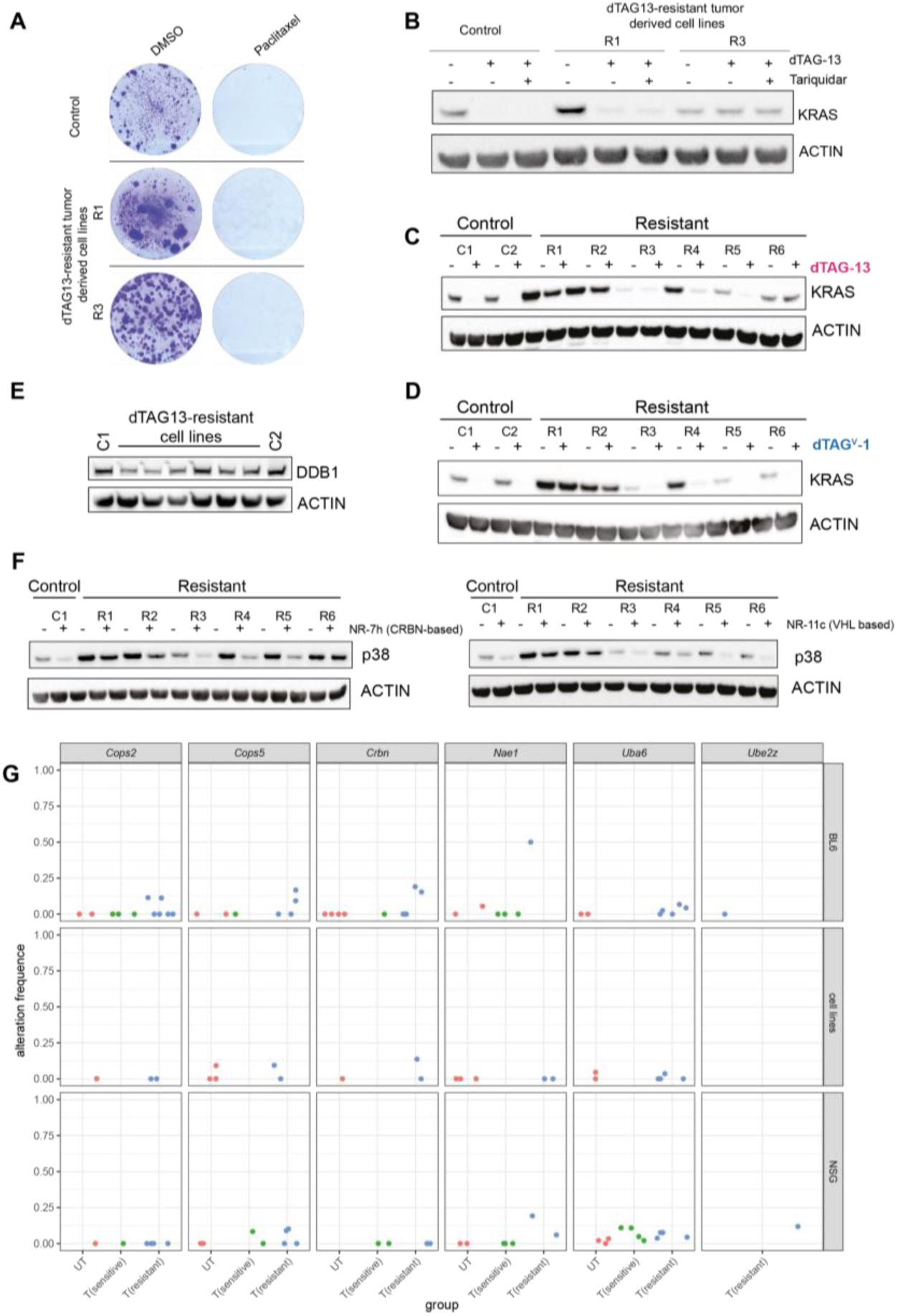
LUAD resistance to targeted degradation *in vivo* is mainly driven by defects in targeted proteolysis. **A,** Colony forming assay in cells derived from dTAG-13 degradation-resistant LUADs. Paclitaxel = 1 µM (minimum dose to affect control cells). **B,** Degradation of dTAG- KRAS^G12V^ in LUAD murine cells derived from mice treated with DMSO or the indicated compounds. (dTAG-13 = 0.3 µM, Tariquidar = 1 µM; 24h). ACTIN is shown as loading control. **C,** Degradation of dTAG-KRAS^G12V^ in LUAD murine cells treated with DMSO or dTAG-13 (0.3 µM, 24h). Cell lines established from treatment-naïve (Controls) and dTAG-13 resistant LUAD tumors. Quantification in Fig.4D. **D,** Degradation of dTAG-KRAS^G12V^ in LUAD murine cells treated with DMSO or dTAG^V^-1 (0.3 µM, 24h). Quantification in Fig.4F. **E,** DDB1 levels in cell lines established from treatment-naïve and dTAG-13 resistant LUAD tumors. ACTIN is shown as loading control. **F,** p38 degradation upon PROTAC treatment (1 µM; 16h) with NR-7h and NR-11c. Cell lines established from treatment-naïve and dTAG- 13 resistant LUAD tumors. ACTIN is shown as loading control. **G,** Frequency of alternative allele in SNPs or INDELs predicted to have a high or moderate impact in protein function. SNP calling with GATK Haplotype caller and annotation with SnpEff (see Methods). BL6: tumors from immunocompetent mice (C57BL/6J). NSG: tumors from immunocompromised mice. UT: untreated (red). T(sensitive): dTAG-13 sensitive tumors (green). T(resistant): dTAG-13 resistant tumors or derived cell lines (blue). BL6 UT tumors n = 4, sensitive n = 4, resistant n = 8. Cell lines UT n = 3, resistant n = 6. NSG UT tumors n = 4, sensitive n = 4, resistant n = 6. Only samples with variants with a minimum coverage of 10 reads are shown.

## METHODS

### Cell lines

*Culture conditions.* Murine LUAD cell lines were grown in DMEM (Gibco) supplemented with 10% FBS and 1% penicillin/streptomycin (pen/strep). KRAS^lox/lox^ MEFs and KRAS^lox/lox^ dTAG- KRAS^G12V^ were grown in DMEM (Gibco) supplemented with 15% FBS and 1% pen/strep. All cell lines were routinely tested for mycoplasma contamination.

*Proliferation and viability assays.* 250 LUAD cells were seeded per well (in tetraplicates) in 96- well plates for apoptotic evaluation. 3 days later, DMSO, dTAG-13 or dTAG^V^-1 degraders (0.5 µM) were added in combination with PE-Annexin-V (Immunostep, ANXVPE-200T) and cell death was monitored using the AI module of an Incucyte SX5 device (Sartorius) for both cell scanning and analysis.

### Plasmid cloning

The lentiviral vector pLEX305_dTAG-2xHA-KRAS^G12V^ was generated by Gateway cloning (Invitrogen). First, KRAS^G12V^ was cloned into a Gateway compatible donor vector (pDONR223) using the BP clonase after PCR with primers containing BP overhangs. Then, we performed a Gateway LR reaction with pENTR223_KRAS^G12V^ and pLEX305-N-dTAG (Addgene #91797), according to manufacturer’s protocol.

### Chemical compounds

dTAG-13, dTAG^V^-1, and IGP002 were synthesized as described in the Supplementary Information. Tariquidar (HY-10550, MedChemExpress). The p38 PROTACs NR-7h (CRBN-based) and NR-11C (VHL-based) were provided by A. Riera^63,64^.

### Lentiviral production for intranasal delivery of dTAG-KRAS^G12V^

*Lentivirus Production.* 293T cells were expanded to 20x p150 plates. On day 1, cells were transfected using a mix containing 407.5 µL DMEM, 2.125 µg of pLEX305_dTAG-2xHA- KRAS^G12V^, 4.25 µg pMD2.G (Addgene #12259), 8.5 µg psPAX2 (Addgene #12260), and 122 µL PEI (Polysciences, 24765-1) per p150 plate. The transfection mix was vortexed, incubated at room temperature (RT) for 10 min, and added to each plate containing 15 mL DMEM. After 72h hours, virus-containing media was collected and concentrated by ultracentrifugation.

*Ultracentrifugation.* Two p150 plates (30 mL total) were used per ultracentrifuge tube (#344058, Beckman Coulter). Samples were centrifuged at 24,000 rpm under vacuum for 1.5 hrs at 4°C (Allegra X-15R Centrifuge, Beckman Coulter). The viral pellet was resuspended in 300 µL PBS, achieving a 100-fold concentration. Concentrated virus was pooled and aliquoted into 300 µL and 50 µL volumes for storage and titration, respectively.

*Virus titration.* HAP1 cells (100,000 per condition) were prepared in 99 µL IMDM supplemented with polybrene. A seven-point 1:10 serial dilution of the virus was performed, ranging from 10 µL to 0.00001 µL. Cells were incubated with virus for 1 hr at 37°C, with gentle mixing every 15 min, then plated on p100 dishes. Puromycin selection was applied 48 hrs post-transduction. After additional 48 hrs, colonies were stained with crystal violet and counted to determine viral titer (1.5 x 10^6^ viral particles/ µL).

#### Western Blot analysis

Cell pellets were washed in cold PBS and lysed with 8 M urea 1% CHAPS with constant shaking at 4°C for 1 hr. The samples were centrifuged at 11,000×g for 5 min at 4°C and the supernatants were collected for protein quantification. For the electrophoresis, 20-25 μg of protein with loading buffer (NuPAGE LDS Sample Buffer, 10% β-mercaptoethanol) were denatured for 10 min at 70 °C, separated on 4-12% SDS-PAGE gels (Invitrogen) and transferred into nitrocellulose membranes using the Trans-Blot Turbo Transfer System. Membranes were blocked in 5% (w/v) non-fat powder milk/TBST for 1hr and later incubated at 4°C overnight with the primary antibodies in TBST. Next day, membranes were washed twice for 5 min in TBS-T and then incubated with the secondary antibodies at RT for 1 hr. Enhanced chemiluminescence (ECL) measurement was performed in a ChemiDoc MP Imaging system.

The primary antibodies used are the following: CUL4A (#2699), DDB1 (#5428), NAE1 (#14321), CRBN (#71810) and p38 (#9217) were purchased from Cell Signaling Technology and used at 1:1000 dilution. β-ACTIN (A5441, used at 1:20000 dilution) and pan-RAS (OP40, used at 1:1000) were purchased from Sigma-Aldrich. The secondary antibodies used are the following: anti-rabbit (115-035-003) and anti-mouse (111-035-003) are from Jackson ImmunoResearch and used at 1:5000.

#### Colony formation assays

Cells were seeded at 1000 cells per well in 12-well plates. Treatment was added the same day and changed every 3-4 days. After 12-15 days, cells were washed and subsequently fixed and stained with 0.33 % (w/v) methylene blue+methanol. After 30 min, the cells were washed in distilled water and air-dried.

#### Chemistry

Detailed synthetic procedures and characterizations are provided as **Supplementary information.**

### Proteomics

*Sample collection*. 15 x 10^6^ cells of each cell line were seeded in triplicates the day before collection. To collect, cells were washed with PBS and trypsinized until detached. After collection into a Falcon and centrifuged for 5 min at 300xg, pellets were washed three times with cold PBS and then snap-frozen in liquid nitrogen and stored at −80°C until processed.

*Cell Lysis and Digestion.* Cell pellets were lysed using Beatbox according to the manufacturer’s instructions. In brief, each cell pellet sample was introduced in an individual BeatBox tube. PreOmics’ LYSE buffer and Pierce^TM^ Universal Nuclease for Cell Lysis (Thermo Scientific) were added. Samples were homogenized into the BeatBox instrument where the settings were set at STANDARD for 2 cycles (10 min each). Cell debris was removed by centrifugation at 16000 g for 5 min at 20°C and supernatant was transferred to new tubes. Protein concentration was determined using the Pierce^TM^ BCA protein Assay (Pierce Biotechnology). An amount of 100 µg of total protein per sample was digested following the PreOmics digestion protocol (iST Sample Preparation Kit 96x, P.O.00027, PreOmics) using Trypsin/LysC mix as enzyme. Incubation was done at 500 rpm for 2h at 37°C. The tryptic peptides eluted were dried down in a speed vacuum centrifuge (Eppendorf).

*RP Liquid Chromatography Mass Spectrometry.* Mass spectrometry was performed on an Orbitrap Eclipse Tribrid mass spectrometer (ThermoFisher Scientific, San Jose, CA) coupled to a EVOSEP One (EVOSEP, Odense, Denmark) via nanoEasy Spray Source interface with a stainless steel emitter (EV-1086 EVOSEP). Tryptic peptides were loaded onto the EVOTIP following the manufacturer’s instructions. The analytical column was a 15 cm x 150 µm ID, with Dr Maisch C18 AQ, 1.9 µm beads. The eluents were 0.1% formic acid in water and 0.1% formic acid in acetonitrile. The Evosep One method was 15 SPD (88 min gradient) and the flow rate 220 nl/min. The mass spectrometer was operated in targeted mode, acquiring data with a tailored Parallel Reaction Monitoring (PRM) method. Unique peptides corresponding to 23 proteins of interest were selected for quantification in 21 samples. Full MS1 scans were acquired in the Orbitrap with a scan range of 350 - 1400 m/z and a resolution of 120,000 (at 200 m/z). Automatic gain control (AGC) was set to a target of 4 x 10^5^ and a maximum injection time of 50 ms. Only MS1 precursors corresponding to the peptides of interest were subjected to MS2 fragmentation and were acquired in the Orbitrap with a resolution of 15,000 (at 200 m/z). Automatic gain control (AGC) was set to a target of 1 x 10^5^ and a maximum injection time of 22 ms. Higher energy collision induced dissociation (HCD) was applied with a normalized collision energy (NCE) of 28 or 30% depending on the m/z (see Supplementary Table 3 with acquired m/z, retention time windows and collision energies applied). Orbitrap Eclipse Tune Application 3.5.3890 and Xcalibur version 4.6.67.17 were used to operate the instrument and to acquire data, respectively. The mass spectrometry proteomics data has been deposited at PRIDE with the identifier PXD058734.

*Data Analysis.* Acquired .raw data files were searched against Mouse SwissProt database (v. June 2024 containing the proteins of interest) using the Proteome Discoverer 3.0.1.27 software, with Sequest HT as the search engine. The Percolator validation node was used to remove false positives with a false discovery rate (FDR) of 1% at the peptide level. Data were searched with mass tolerances of ± 10 ppm and ± 0.02Da on the precursor and fragment ions, respectively. These results were used to build a spectrum library to facilitate the integration of the peaks for the quantitative analysis. For that, the .raw data files were analyzed in Skyline v.24.1.0.199. Peaks were picked automatically, using the default Skyline peak picking model and Savitzky–Golay smoothing was applied. Areas corresponding to the selected peptides were reviewed and after integration and manual validation of the available transition peak areas, total areas for each peptide were exported in .csv format to be further analyzed. Protein quantitation was done by performing MaxLFQ algorithm over the tracked peptide areas for a given protein, using diann_maxlfq from the diann package of the R statistical software (v4.3.0; R Core Team 2023). Comparison between groups was done (each resistant cell line vs. control). For each comparison, estimated log2 fold changes were calculated from the mean of each cell line triplicate.

### SNP calling from bulk RNAseq

*Custom Genome Construction.* A custom genome was generated by incorporating the dTAG- 2xHA-KRAS sequence into the Mus musculus version 10 genome assembly. *Gene Annotation.* Coding exon regions from genes of interest were extracted from the Ensembl database using the biomaRt package in R.

*Variant Discovery.* To identify new variants (SNPs and Indels) for each sample in the regions of interest, the RNA-seq short variant discovery pipeline from the GATK website was utilized with minor customizations.

RNA-seq reads were aligned using a two-step alignment strategy with STAR (v2.7.10a). First alignment was perfomed using outFilterMismatchNoverLmax = 0.05; outFilterMatchNmin = 25; genomeLoad = LoadAndRemove and outSAMstrandField = intronMotif as parameters. A genome generation step (--genomeGenerate) was performed with the parameter sjdbOverhang = 50. Second Alignment was performed using genomeLoad = LoadAndRemove and outSAMstrandField = intronMotif as parameters. Post-alignment data cleanup was performed using Picard Tools (v1.98). The following steps and parameters were applied: AddOrReplaceReadGroups with default settings; MarkDuplicates with REMOVE_DUPLICATES = TRUE; ASSUME_SORTED = TRUE and VALIDATION_STRINGENCY = LENIENT as parameters and ReorderSam with ALLOW_CONTIG_LENGTH_DISCORDANCE = TRUE. Variant calling was conducted using GATK (v4.5.0.0): Data was prepared for variant calling using the SplitNCigarReads function and nucleotide recalibration was conducted by applying BaseRecalibrator utilizing the dbSNP137 database for known variants and ApplyBQSR to apply base quality score recalibration. Variants were identified specifically within the exonic regions of the selected genes using the HaplotypeCaller function with parameters ERC = GVCF; dont-use-soft-clipped-bases = TRUE and standard-min-confidence-threshold-for-calling = 20.0. Then, a variant filtering step was performed using the VariantFiltration with parameters cluster-window-size = 35; cluster-size = 3; FS > 30.0 and QD < 2.0.

*Joint-call cohort analysis.* Joint-call cohort analysis was conducted using GATK (v4.5.0.0). Individual sample VCF files were transformed into a GenomicsDB using the GenomicsDBImport function. SNPs and indels were jointly called using the GenotypeGVCFs function.

*Variant annotation and filtering.* Variant annotation of SNPs and indels separately, was performed using the SelectVariants tool. Finally, SNPs and indels were filtered using the VariantFiltration function with parameters cluster-window-size = 35; cluster-size = 3; FS > 30.0 and QD < 2.0.

### *In vivo* tumor studies

*Generation of dTAG-KRAS^G12V^ cell lines.* 7.5×10^7^ pLEX305_dTAG-2xHA-KRAS^G12V^ viral particles in PBS were delivered into C57BL/6J mice by intranasal administration. Animals were monitored until signs of distress compatible with LUAD development were observed (average latency of 10 months). Mice were sacrificed and tumors were collected and orthotopically implanted in the lung of NSG recipient mice as an intermediate step for the development of primary cancer cell lines as previously described^49^. dTAG-KRAS^G12V^ LUAD cells used for *in vivo* experiments were infected with lentiviruses expressing ZsGreen (a kind gift from Héctor Peinado, CNIO, Spain) and luciferase (). dTAG-13 resistant cell lines and paired controls were derived from vehicle-treated and dTAG-13 refractory tumors respectively developed upon prolonged exposure to the degrader *in vivo* (see below). Those cell cultures established from resistant tumors were mantained in medium supplemented with dTAG-13 (0.5 µM).

*Tumor engraftment and KRAS^G12V^ targeted degradation.* All animal experiments were approved by the Bioethics Committee of Salamanca University and the animal experimentation authorities of the autonomous government of Castilla y León (Spain) -animal license number #736-. Mice were kept in ventilated rooms in pathogen-free facilities under controlled temperature (23 °C), humidity (50%) and illumination (12 hr light/12 hr dark cycle) conditions. After weaning, mice were fed a standard chow diet (#2918, Harlan Laboratories). Animals were treated humanely in accordance with standards described in the Guide for the Care and Use of Laboratory Animals considering relevant national and European guidelines. For orthotopic tumor induction, 10^5^ dTAG-KRAS^G12V^ LUAD cells co-expressing both immunofluorescence and luminescence reporters, were injected in 100 μl of PBS via the tail vein of 8-week-old C57BL/6 or NSG mice. Tumor growth was routinely monitored by measuring the bioluminescence signal. For image acquisition, mice were injected intraperitoneally with 100 μL of a 30 mg/ml D-Luciferin potassium salt solution (LUCK-100, Gold Biotechnology) 10 min before imaging and were then anesthetized using 2% isoflurane. The bioluminescent signal was quantified as total photon flux by measuring at the site of lung tumors using a region of interest (ROI), which directly correlates the signal with the size of the tumor. Bioluminescence imaging and data processing were performed using the IVIS Lumina S5 imager (Revvity) equipped with the Living Image 4.7.3 software, starting on day 7 after tumor cell injection. Prior to treatment initiation, when tumor size reaches a pre-determined bioluminescence level (between 7.5-8 x 10^8^ p/s), mice with similar photon counts were randomized to either the control or treatment groups. Then, mice assigned to each of the groups received a daily intraperitoneal 200 μl injection of either solvent solution (20% Polyoxyl 15 hydroxystearate USP – GX7347, Glentham Life Sciences-in PBS, control group) or dTAG-13 degrader (40 mg/kg, treatment group) during the first week, a period in which IVIS imaging was conducted at 3 and 7 days post-treatment. Afterwards, a 4-day/week degrader administration with *in vivo* imaging follow-up once a week was implemented. Undetectable bioluminescence signal in two consecutive measurements resulted in treatment discontinuation. Subsequently, the 4-day/week degrader administration was restarted upon detection of bioluminescence signal. When measurements revealed recurrent increasing bioluminescence counts in two consecutive weekly measurements, animals were considered refractory to the treatment and were sacrificed.

*Immunohistochemistry.* Tumor-bearing lungs were formalin-fixed for 24 hrs and paraffin embedded accroding to standard methods. Tissue sections were stained with hematoxylin and eosin for histophathological evaluation. Additional sections were used for immunohistochemical stainings using the following antibodies: HA-tag (Cell Signaling, 3724), ZsGreen (Takara Bio, 632474), phospho-Histone H3 pSer10 (Sigma, H0412), Cleaved Caspase-3 Asp175 (Cell Signaling, 9661), pERK1/2 (Cell Signaling, 9101), Phospho-S6 Ribosomal Protein Ser240/244 (Cell Signaling, 2215), CD8 (CNIO, OTO94A) and F4/80 (Bethyl, A700-209). A quantitative histology score (H-score) considering both the intensity of the staining and the area containing a positive signal was automatically calculated by QuPath 0.5.1 software.

### Tumor immunophenotyping studies

*Sample preparation.* Mice were euthanized by cervical dislocation and 100-200 μl of peripheral blood was collected via cardiac punctures in EDTA-coated tubes for subsequent hemocytometer and flow cytometry analysis. Tumor bearing lungs were dissected and minced by using sterile scissors and added into gentleMACS C tubes (Miltenyi, 130-093-237) with 2.5 mL of the digestion mix (Miltenyi, tumor dissociation kit 130-096-730). Samples were incubated at 37° C during 40 min and mechanically processed into single cell suspensions by gentleMACS dissociator (130-093-235, using program m_impTumor_04.01). After homogenization, samples were filtered through a 70 μm nylon cell strainer, erythrocytes were eliminated using a red-cell lysis buffer (15 mM NH_4_Cl, 1 mM KHCO_3_, 0.1 mM EDTA). Samples were washed and blocked on ice during 15-20 min in FACS buffer (2 mM EDTA and 0.5% bovine serum albumin in PBS). After blocking, a small volume of the cell suspension was separated for subsequent anti-CD45 staining (APC-CD45 Biolegend, 147708) required for estimating immune cell infiltration into the lung tissue by flow cytometry (designated as pre-depleted fraction). The remaining sample was incubated with magnetically labeled mouse anti-CD45 antibody conjugated microbeads (Miltenyi, 130-052-301), in order to separate both CD45^+^ and CD45^—^ fractions, representing immune and non-immune cell populations respectively. While the CD45+ cell fraction was immunophenotyped (see below), CD45^—^ cells were subjected to a second round of CD45 staining and FACs separation in order to enhance the elimination of immune contaminants. Finally, tumor cells positive for ZsGreen expression were isolated by FACS from this ultra pure CD45^+^ cell fraction and processed for transcriptomic analysis.

*Immunophenotyping protocols.* Characterization of tumor and blood immune cell components was conducted using 50 μL of EDTA-anticoagulated plasma-depleted blood, 2.5×10^5^ cells of the crude tumor suspension and 2×10^6^ cells from the CD45^+^/ZsGreen^+^-enriched tumor fraction. All samples were stained employing a stain-lyse-wash procedure. Briefly, samples were washed with PBS at pH 7.4 for 5 min at 540 x g and pre-incubated with anti-CD3 antibody (for C57BL6 samples), True-Stain Monocyte Blocker (Biolegend, 426103), TruStain FcX™ PLUS (Biolegend, 156604), and the Zombie NIR viability maker (Biolegend, 423106) at a 1:2000 dilution for 30 min at RT, protected from light. Subsequently, the remaining antibodies (see Supplementary Table 4).) were added, and the samples were incubated the presence of Brilliant Staining Buffer Plus (BD Biosciences, 566385) for 30 min at RT (protected from light) in a roller. To lyse red blood cells and fix the samples, 1x BD FACS Lysing Solution (BD Biosciences, 349202) was added and incubated for 10 min at RT (protected from light), followed by centrifugation at 540 x g for 5 min, washing with PBS containing 0.5% bovine serum albumin (BSA), 0.1% sodium azide, and 2 mM EDTA (pH 7.4), and resuspension in 400 µL of PBS prior to analysis.

Data acquisition was performed using an Aurora spectral flow cytometer (Cytek) equipped with five lasers (355 nm, 405 nm, 488 nm, 561 nm, 640 nm). Daily instrument setup and quality control were conducted according to manufacturer instructions prior to sample measurement. Single-stained reference controls for each fluorochrome in the antibody combination, as well as an unstained control sample, were processed identically to the multicolor-stained samples to ensure accurate spectral unmixing. The resulting unmixing matrix was generated using SpectroFlo software (v3.3.0; Cytek). Data analysis was performed using Infinicyt™ software (BD Biosciences, version 2.1.0.a.000). Absolute counts in blood samples were determined using a double-platform method based on nucleated cell quantification obtained from the Element HT5 automated hematological analyzer (Heska).

### Transcriptomic studies

*RNA sequencing.* ZsGreen+/CD45- LUAD cells were were sent to CeGaT GmbH for RNA isolation (MagMax, ThermoFisher), library preparation (SmartSeq stranded, Takara Bio) and sequencing (NovaSeq 6000, Ilumina). Bioinformatic analysis included demultiplexing using llumina bcl2fastq (version 2.20) and FastQC analysis (version 0.11.5-cegat) prior to completing in-house bioinformatics studies (see below).

*Bioinformatic pipeline.* RNA sequencing data analysis was performed using the following workflow: (1) the quality of the samples was verified using FastQC software (https://www.bioinformatics.babraham.ac.uk/projects/fastqc/); (2) the alignment of reads to the mouse genome (GRCm39) was performed using STAR^65^; (3) gene expression quantification using read counts of exonic gene regions was carried out with featureCounts^66^; (4) the gene annotation reference was Gencode M34^67^; and (5) differential expression statistical analysis was performed using R/Bioconductor. Data will be publicly available in GEO database. Gene expression data was normalized with edgeR^68^ and voom^69^. After quality assessment and outlier detection using R/Bioconductor, a filtering process was performed. Genes with read counts lower than 6 in more than the 50% of the samples of all the studied conditions were considered as not expressed in the experiment under study. LIMMA^70^ was used to identify the genes with significant differential expression between experimental conditions. Further functional and clustering analyses were performed and graphical representations were generated using clusterProfiler and R/Bioconductor. The functional analyses included Gene Set Enrichment Analysis (GSEA) with MSigDB Hallmarks collection of gene sets^71^. Additional GSEA analyses were performed using a recently reported KRAS-dependent transcriptome^50^.

## ACKNOWLEDGMENTS

We would like to thank the members of the Mayor-Ruiz lab and Santamaría lab for helpful discussion that helped shape this work and G.E. Winter (CeMM) for initial support and ideas. We apologize to those colleagues who could not be cited due to space limitations. We thank the Histopathology Facility, and the Mass Spectrometry & Proteomics Facility at IRB Barcelona (the latter granted in the 2014-2020 ERDF Operational Programme; IU16-015983). The Mayor-Ruiz lab acknowledges funding from the European Research Council (ERC) as a Starting Grant (ERC-2021-StG-101040046 TrickE3), AECC (Cancer Grant Challenges partnership, PROTECT team), the Spanish Ministry of Science and Innovation (PID2020-120110RA-I00, PID2023-147995OB-I00, and RYC2020-030061-I), FarmaIndustria, TRIP-Clinics (GenCat), and AGAUR-GenCat (2021SGR01279). I.G.-P. is supported by La Caixa INPhiNIT PhD fellowship (LCF/BQ/DR22/11950030). A.R. acknowledges grants from the Spanish Ministry of Science and Innovation (PID2023-147298NB-I00) and from GenCat (2021 SGR 00866). The contract of C.R.-M. has been funded by the European Union (Next Generation, EU). The Santamaría lab received funding from Spanish Ministry of Science and Innovation (PID2020-116824RB-I00) and by FCAECC (Excellence Program 2022 -EPAEC222641CICS- and Proyectos Generales AECC 2022 -PRYGN222960SANT-).

