## Supplementary figures and images for "Targeted KRAS^G12V^ degradation elicits efficient and durable lung adenocarcinoma regression *in vivo*"

### Supplementary Table 1

#### dTAG-KRAS DNA sequencing by Sanger

##### Control cell line

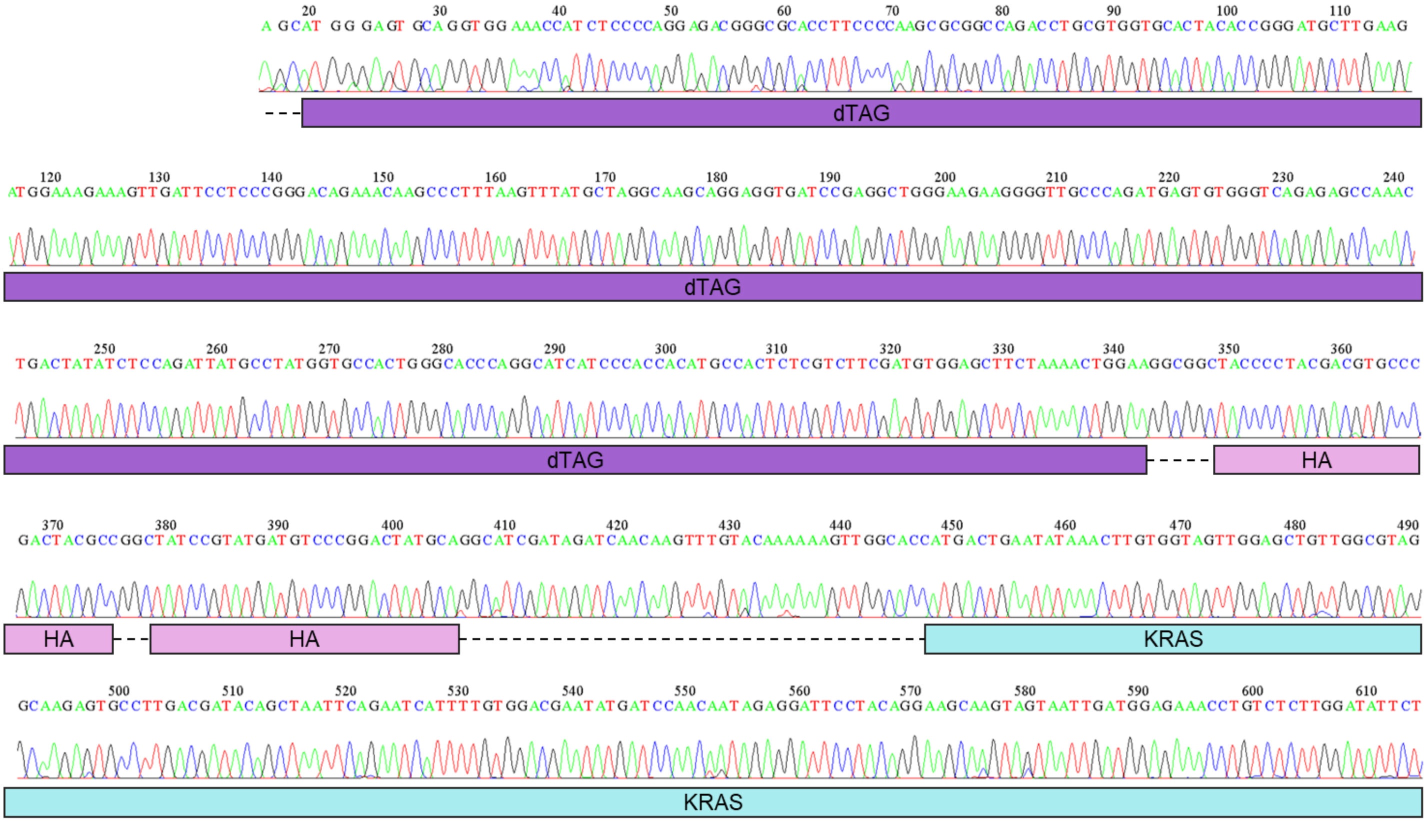


##### dTAG13 resistant cell lines

1

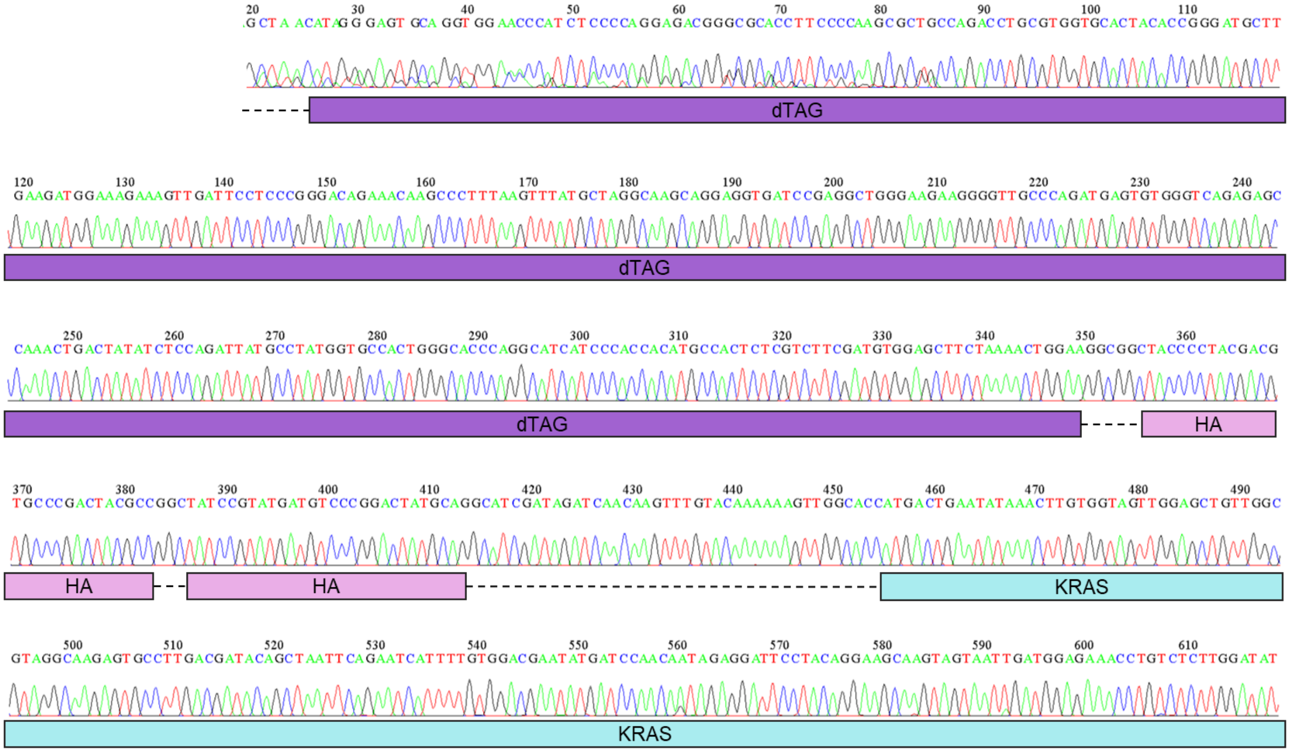

2

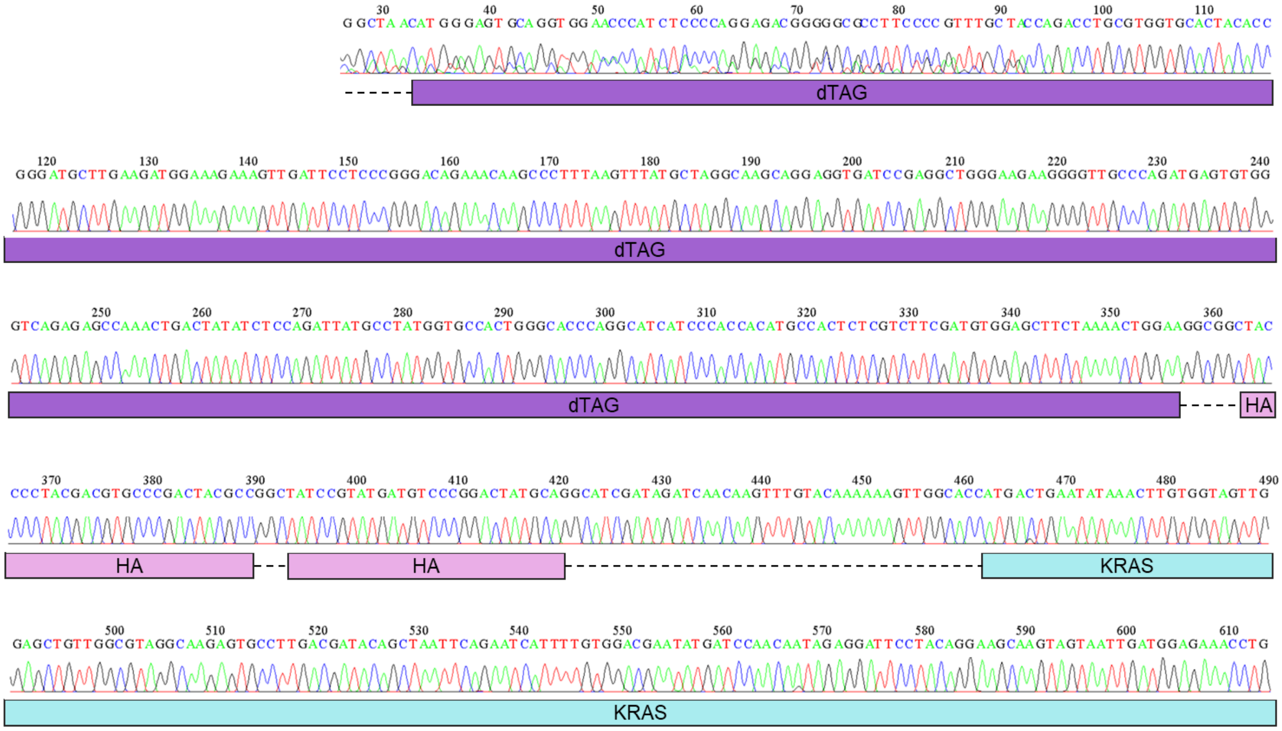


3

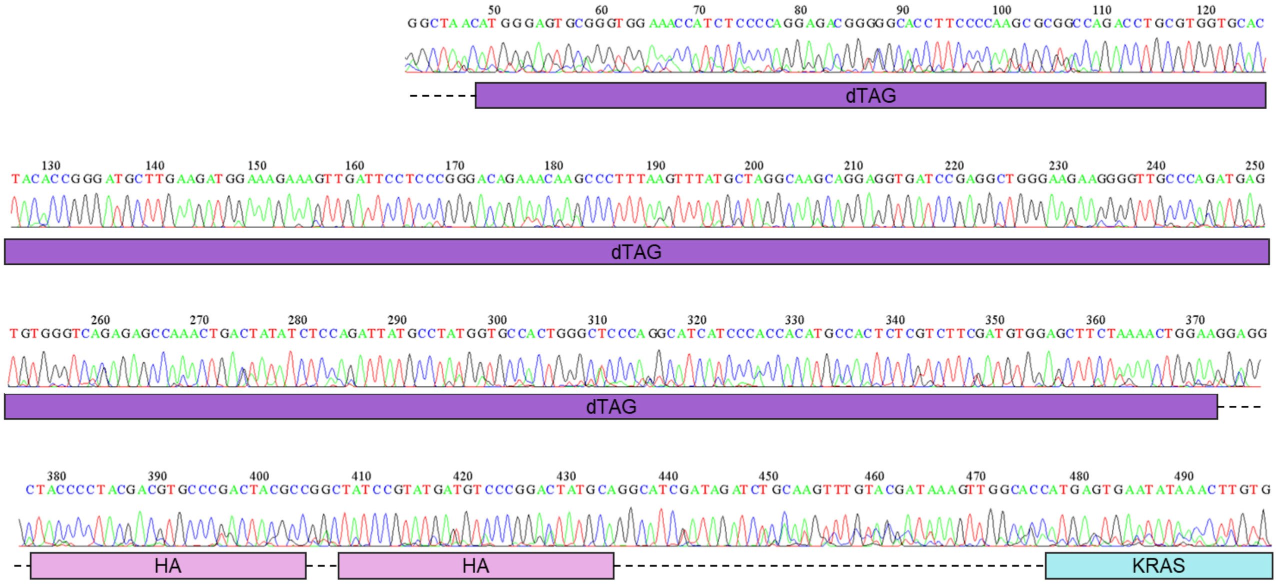

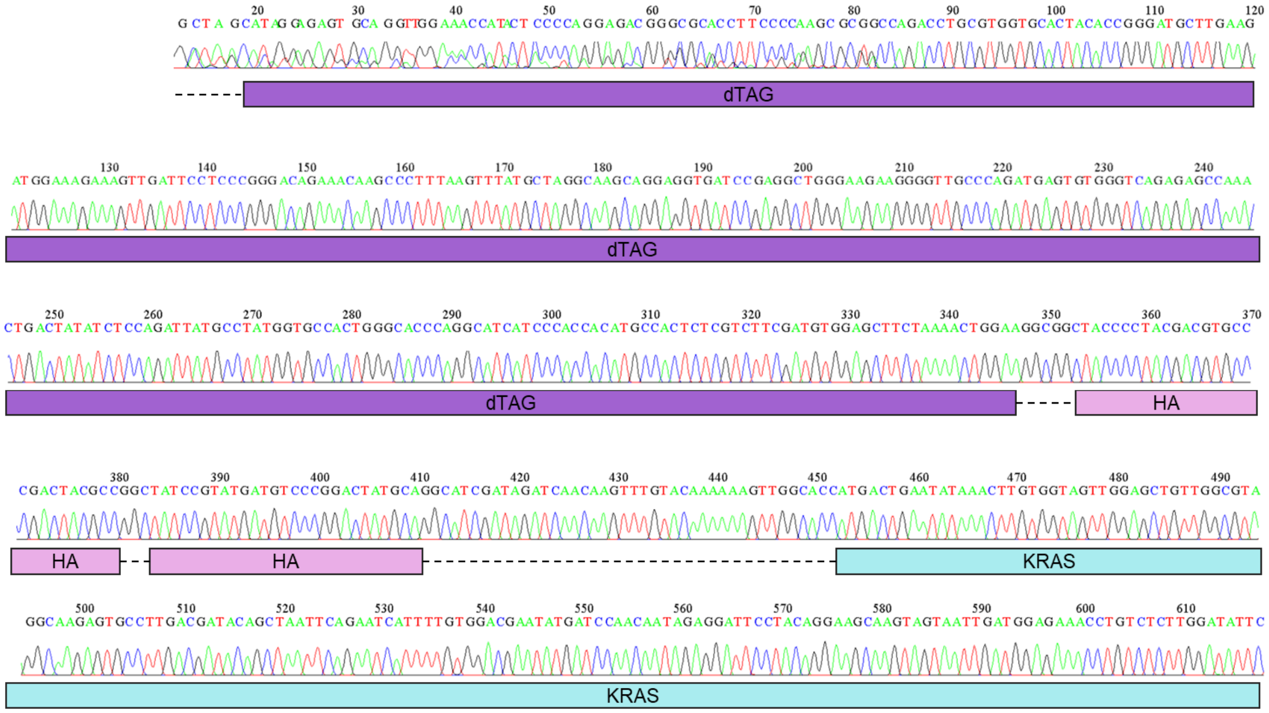

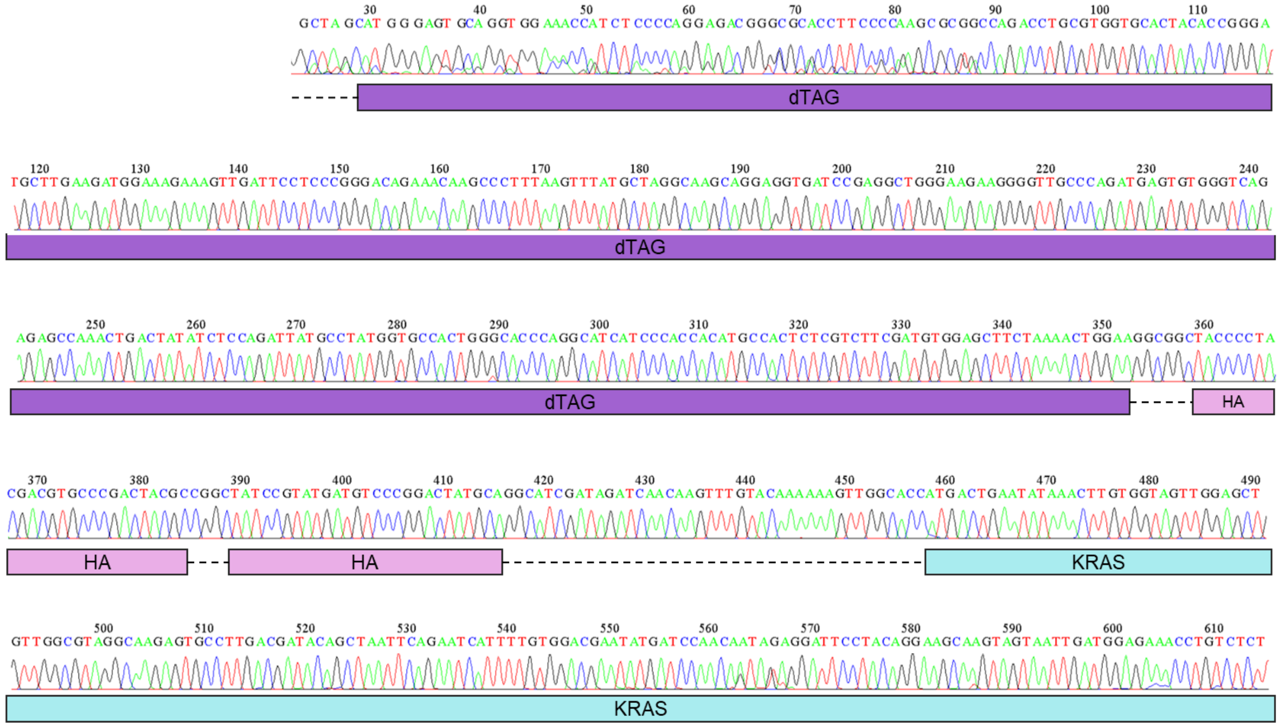
