## Supplementary Table 4 for "Targeted KRAS^G12V^ degradation elicits efficient and durable lung adenocarcinoma regression *in vivo*"

**Supplementary Table 4**. Antibodies and reagents employed in flow cytometry and cell sorting assays

| Antibody/Reagent | Clone | Source | Identifier | Panel |
|---|---|---|---|---|
| CD3 BUV496 | 17A2 | BD Biosciences | Cat# 569671, RRID:AB_3668952 | 1,2,3,4 |
| CD4 BV605 | GK1.5 | BD Biosciences | Cat# 743156, RRID:AB_2741309 | 3,4 |
| CD5 BV750 | 53-7.3 | BD Biosciences | Cat# 747114, RRID:AB_2871865 | 3,4 |
| CD8a BUV395 | 53-6.7 | BD Biosciences | Cat# 563786, RRID:AB_2732919 | 1,2,3,4 |
| CD11b BV421 | M1/70 | BioLegend | Cat# 101236, RRID:AB_11203704 | 1,2,3,4 |
| CD11c BUV615 | N418 | BD Biosciences | Cat# 751222, RRID:AB_2875243 | 1,2,3,4 |
| CD19 RB545 | 1D3 | BD Biosciences | Cat# 569727, RRID: AB_3668953 | 1,2,3,4 |
| CD21/CD35 APC | 7E9 | BioLegend | Cat# 123412, RRID:AB_2085160 | 3,4 |
| CD23 BUV737 | B3B4 | BD Biosciences | Cat# 749668, RRID:AB_2873931 | 3,4 |
| CD25 BV480 | PC61 | BD Biosciences | Cat# 566120, RRID:AB_2739522 | 3,4 |
| CD27 BB700 | LG.3A10 | BD Biosciences | Cat# 742135, RRID:AB_2871394 | 3,4 |
| CD43 BV510 | S7 | BD Biosciences | Cat# 563206, RRID:AB_2738069 | 1,2,3,4 |
| CD44 APC Vio770 | REA664 | Miltenyi Biotec | Cat# 130-118-695, RRID:AB_2733120 | 3,4 |
| CD45 PerCP Cy5.5 | I3/2.3 | BioLegend | Cat# 147705, RRID:AB_2563537 | 1,2 |
| CD45 APC | I3/2.3 | BioLegend | Cat# 147708, RRID:AB_2563540 | 5,6 |
| CD45.2 PerCP | 104 | BioLegend | Cat# 109826, RRID:AB_893349 | 3,4 |
| CD45R (B220) VioBlue | RA3-6B2 | Miltenyi Biotec | Cat# 130-118-321, RRID:AB_2733945 | 1,2,3,4 |
| CD62L BUV805 | MEL-14 | BD Biosciences | Cat# 569201, RRID:AB_3668954 | 3,4 |
| CD64 PE | X54-5/7.1 | BioLegend | Cat# 139303, RRID:AB_10613467 | 1,2,3,4 |
| CD69 BUV661 | H1.2F3 | BD Biosciences | Cat# 741478, RRID:AB_2870943 | 3,4 |
| CD80 BV786 | 16-10A1 | BD Biosciences | Cat# 740888, RRID:AB_2740537 | 2,4 |
| CD93 RY586 | AA4.1 | BD Biosciences | Cat# 753131, RRID:AB_3668955 | 3,4 |
| CD95 RB744 | Jo2 | BD Biosciences | Cat# 756863, RRID:AB_3668956 | 3,4 |
| CD103 PE Fire640 | QA17A24 | BioLegend | Cat# 156917, RRID:AB_2924487 | 1,2,3,4 |
| CD127 RB780 | A7R34 | BD Biosciences | Cat# 569066, RRID:AB_3668957 | 3,4 |
| CD138 PE Cy5 | 281-2 | BD Biosciences | Cat# 142539, RRID:AB_2894661 | 3,4 |
| CD163 PE Cy7 | TNKUPJ | Thermo Fisher Scientific | Cat# 25-1631-82, RRID:AB_2815112 | 2,4 |
| CD185 PE CF594 | 2G8 | BD Biosciences | Cat# 562856, RRID:AB_2737842 | 3,4 |
| CD279 (PD1) APC Fire810 | 29F.1A12 | BioLegend | Cat# 135251, RRID:AB_2910292 | 1,2,3,4 |
| CD304 BUV563 | V46-1954 | BD Biosciences | Cat# 752460, RRID:AB_2917455 | 3,4 |
| CD335 (NKp46) BV711 | 29A1.4 | BioLegend | Cat# 137621, RRID:AB_2563289 | 1,2,3,4 |
| F4/80 SparkNIR685 | BM8 | BioLegend | Cat# 123168, RRID:AB_2924461 | 1,2,3,4 |
| GL7 RB705 | GL7 | BD Biosciences | Cat# 570602, RRID:AB_3668959 | 3,4 |
| I-A/I-E (MHC-II) BV650 | M5/114.15.2 | BD Biosciences | Cat# 563415, RRID:AB_2738192 | 1,2,3,4 |
| IgD BV605 | 11-26c.2a | BD Biosciences | Cat# 563003, RRID:AB_2737944 | 3,4 |
| IgM BUV395 | R6-60.2 | BD Biosciences | Cat# 564025, RRID:AB_2738550 | 3,4 |
| Ly6C PE Fire810 | HK1.4 | BioLegend | Cat# 128061, RRID:AB_3083119 | 1,2,3,4 |
| Ly6G Ly6G | 1A8 | BD Biosciences | Cat# 567039, RRID:AB_2916402 | 1,2,3,4 |
| NK1.1 BV570 | PK136 | BioLegend | Cat# 108733, RRID:AB_10896952 | 3,4 |
| Siglec F AF647 | E50-2440 | BD Biosciences | Cat# 562680, RRID:AB_2687570 | 1,2,3,4 |
| TCRβ RB545 | H57-597 | BD Biosciences | Cat# 756204, RRID:AB_3668961 | 3,4 |
| TruStain FcX PLUS | S17011E | BioLegend | Cat# 156604, RRID:AB_2783138 | 1,2,3,4,5 |
| Brilliant Stain Buffer Plus | N/A | BD Biosciences | Cat# 566385 | 1,2,3,4 |
| True-Stain Monocyte Blocker | N/A | BioLegend | Cat# 426102 | 1,2,3,4 |
| BD FACS™ Lysing Solution (10x) | N/A | BD Biosciences | Cat# 347691 | 1,2,3,4 |
| Zombie NIR™ Fixable Viability Kit | N/A | BioLegend | Cat# 423105 | 1 2,3,4,5 |
| DAPI | N/A | PENDING | PENDING | 6 |

*Panel:* 1, Blood NSG; 2, Tumor CD45+ cells NSG; 3, Blood C57BL6; 4, Tumor CD45+ cells C57BL6; 5; pre-depleted tumor cell suspension; 6, cell sorting.

*Abbreviations:* AF, Alexa fluor; APC, allophycocyanin; BB; Brilliant Blue; BUV, Brilliant Ultraviolet; BV, Brilliant Violet; Cat#, catalog number; CF, Cyanin-based Fluorescent dye; Cy7, Cyanin7; DAPI, 4',6-diamidino-2-phenylindole, dihydrochloride; NIR, near-infrared; PE, Phycoerythrin; PerCP Cy5.5, Peridinin-chlorophyll-protein-cyanin 5.5; RB, RealBlue; RY, RealYellow; R, Red; RRID, Research Resource Identifier
