## Supplementary Material Chemistry for "Targeted KRAS^G12V^ degradation elicits efficient and durable lung adenocarcinoma regression *in vivo*"

**SUPPLEMENTARY INFORMATION. CHEMISTRY**

**Table of contents.**

1.-Chemistry. General methods and instrumentation................................................S2

2. Experimental procedures for the synthesis of dTAG-13.........................................S3

3. Experimental procedure for the synthesis of dTAG-v1 .......................................S13

4. Experimental procedure for the synthesis of IGP002...........................................S16

5. $^{1}$H and $^{13}$C NMR spectra ..........................................................................................S18

#### 1.-Chemistry. General methods and instrumentation

Unless otherwise specified, starting materials were obtained from commercial sources and used without further purification. Unless otherwise noted, air- and moisture-sensitive reactions were carried out in oven-dried (55 °C) glassware capped with a rubber septum under a positive pressure of nitrogen. Air- and moisture- sensitive reagents, solvents, and solutions were transferred via syringe or stainless-steel cannula under a dry nitrogen atmosphere. Reactions were stirred using over-dried Teflon-coated magnetic stir bars. R.t. indicates a temperature in the span of 15–25 °C. Overnight (o.n.) indicates a time span of 16 to 22 h. Solvents were removed using rotary evaporators. Commercial reagents and solvents were used as received without further purification unless otherwise noted. Anhydrous and degassed dichloromethane and tetrahydrofuran were taken from a solvent purification system (SPS PS-MD-3). Other anhydrous solvents were purchased from Sigma Aldrich. Normal phase chromatography was performed using an automated chromatography system (PuriFlash® 430, Interchim). Reverse phase chromatography was performed using an automated chromatography system (CombiFlash® NextGen 300+, Teledyne).

**NMR spectroscopy:** $^{1}$H and $^{13}$C were recorded on the NMR spectrometers of the Centres Científics i Tecnològics de la Universitat de Barcelona. The employed spectrometers were a Varian Mercury 400 MHz, a Bruker 400 MHz, or a Bruker 500 MHz. Chemical shifts (δ) were referenced to internal solvent resonances and reported relative to TMS (tetramethylsilane). The coupling constants (*J*) are reported in Hertz (Hz).

The following abbreviations are used to define multiplicities: s (singlet), d (doublet), t (triplet), q (quartet), dd (doublet of doublets), ddd (doublet of doublet of doublets), ddd (doublet of doublet of triplets), dddd  doublet of doublet of doublet of doublets), dt (doublet of triplets), td (triplet of doublets), ddq (doublet of doublet of quartets), dtd (doublet of triplet of doublets), dq (doublet of quartets), tt (triplet of triplets), qt (quartet of triplets), m (multiplet), br s (broad signal).

**HPLC analysis** were performed on an Agilent Technologies Series 1100 chromatograph with UV detector by Enantia S.L. The conditions for each analysis are specified in every case.

**Melting points** were determined using a Büchi M-540 apparatus.

#### 2. Experimental procedures for the synthesis of dTAG-13.

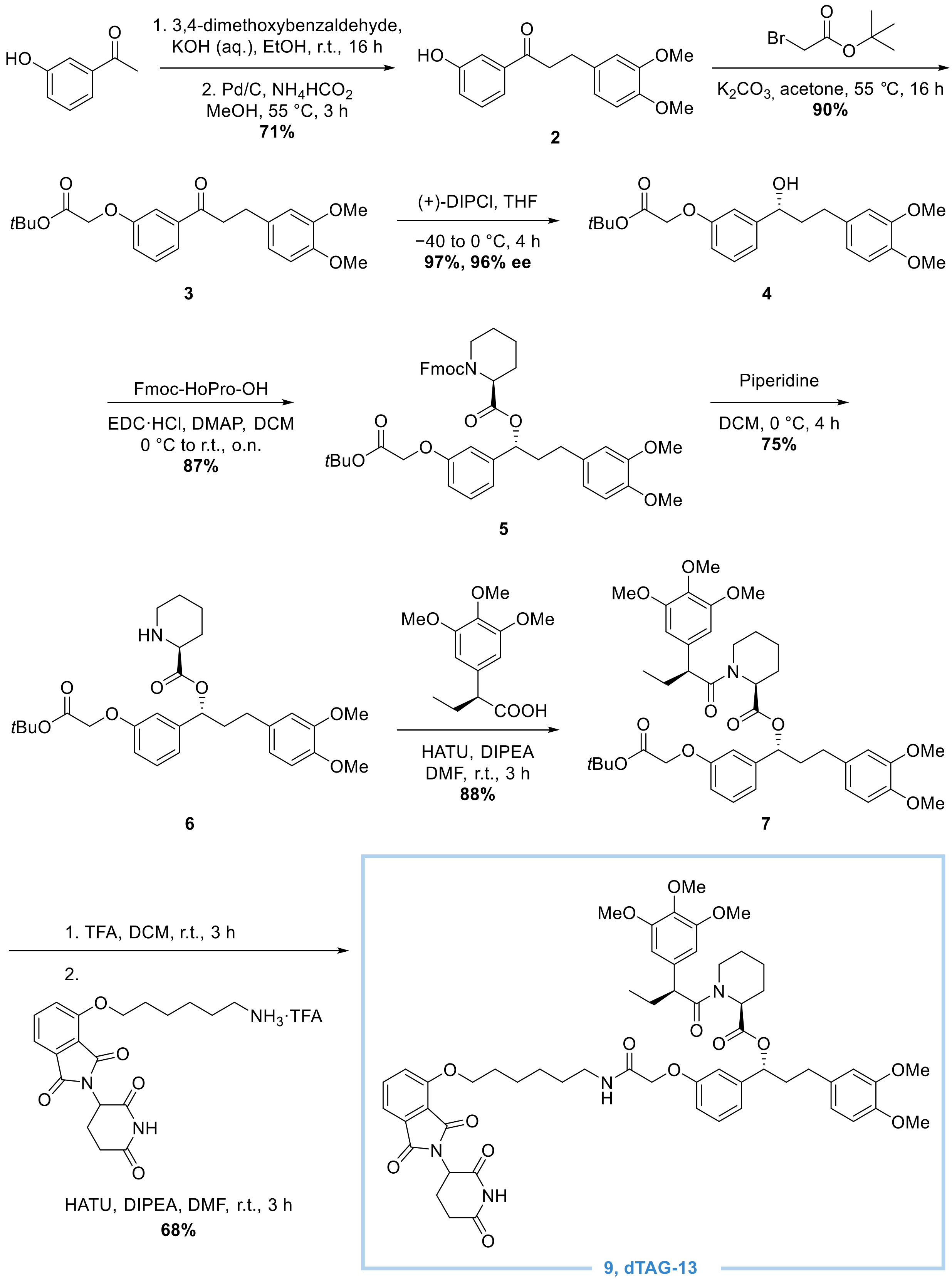


**Supplementary Figure S1.** General synthetic scheme of the dTAG-13 synthesis.

2.1 Synthesis of 3-(3,4-dimethoxyphenyl)-1-(3-hydroxyphenyl)prop-2-en-1-one (**1**)

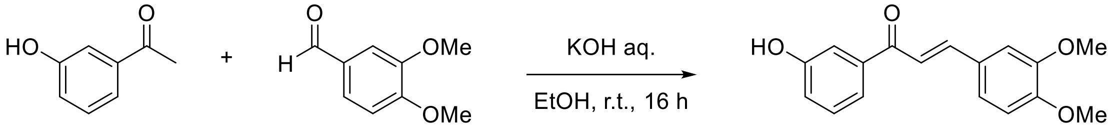


**1**

3-Hydroxyacetophenone (50.0 g, 367.2 mmol) and 3,4-dimethoxybenzaldehyde (61.0 g, 367.2 mmol) were placed in a 2 L round-bottom flask and dissolved in ethanol (490 mL, 0.75M). The reaction was cooled down to 0 °C, a solution of KOH pellets (97.0 g, 1.47 mol, 4 eq.) in water (76 mL) was charged in an addition funnel and added dropwise over 1 h. Then, the ice bath was removed, and the reaction was left to stir at room temperature overnight. The reaction was cooled down to 0 °C and HCl 37% was added until pH 3. Then, water was added (6.3V in relation to 3,4-dimethoxybenzaldehyde) and the solution was left stirring at room temperature for 3 h. The product was filtered, washed with water (3 x 2V) and dried in the oven at 40 °C for 48 h to constant mass. Compound **1** was obtained as a yellow solid (96.4 g, 92%). Spectroscopic data were in excellent agreement the reported in the literature.[1] **$^{1}$H NMR (400 MHz, DMSO)** δ 9.77 (s, 1H), 7.62 (dt, *J* = 7.9, 1.2 Hz, 1H), 7.53 (d, *J* = 2.0 Hz, 1H), 7.45 (dd, *J* = 2.6, 1.6 Hz, 1H), 7.39 – 7.34 (m, 2H), 7.05 (dq, 1H), 7.02 (d, *J* = 8.4 Hz, 1H), 3.86 (s, 3H), 3.82 (s, 3H) ppm.

2.2 Synthesis of 3-(3,4-dimethoxyphenyl)-1-(3-hydroxyphenyl)propan-1-one (**2**).

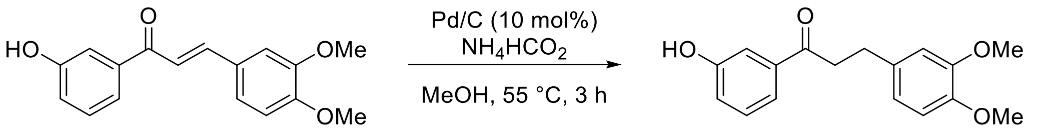


**1** **2**

Compound **1** (30.0 g, 105.5 mmol) and 10 mol% Pd/C 50% wet (3.0 g, 10% wt) were placed in a two-necked round-bottom flask equipped with a stir bar under $N_2$ atmosphere. The reactants were dissolved in anhydrous MeOH (150 mL, 0.7M) and ammonium formate (10.3 g, 158.3 mmol, 1.5 eq.) was added with positive nitrogen current. The reaction was heated up to 55 °C and stirred for 1.5 h. After this time,

[1] Liu, P.; Calderon, A.; Konstantinidis, G.; Hou, J.; Voss, S.; Chen, X.; Li, F.; Banerjee, S.; Hoffmann, J.-E.; Theiss, C.; Dehmelt, L.; Wu, Y.-W.: A Bioorthogonal Small-Molecule-Switch System for Controlling Protein Function in Live Cells. *Angew. Chem. Int. Ed.* **2014**, *53*, 10049-10055

the reaction progress was analysed by TLC, and additional batches of 3.3% wt of Pd/C were added until the observed conversion was almost complete. After that, the reaction was filtered through a celite plug and washed with abundant MeOH. The filtrate was concentrated to dryness under reduced pressure and redissolved in hot ethyl acetate. Then, it was filtered through a $SiO_2$ plug and washed with hot EtOAc. The crude was recrystallized in MeOH (2V), by heating up to 80-90 °C and gradually cooling down to 0 °C. Then, the product was filtered and washed once with a minimal amount of cold MeOH. Compound **2** was obtained as a white solid (23.3 g, 77%). Spectroscopic data were in excellent agreement the reported in the literature.[1] **$^1$H NMR (400 MHz, DMSO)** δ 9.74 (s, 1H), 7.43 (dt, *J* = 7.6, 1.2 Hz, 1H), 7.33 – 7.28 (m, 2H), 7.01 (ddd, *J* = 8.0, 2.5, 1.0 Hz, 1H), 6.88 (d, *J* = 2.0 Hz, 1H), 6.83 (d, *J* = 8.1 Hz, 1H), 6.75 (dd, *J* = 8.2, 2.0 Hz, 1H), 3.73 (s, 3H), 3.70 (s, 3H), 3.27 (dd, *J* = 8.1, 7.0 Hz, 2H), 2.85 (t, *J* = 7.5 Hz, 2H) ppm.

2.3 Synthesis of *tert*-butyl 2-(3-(3-(3,4-dimethoxyphenyl)propanoyl)phenoxy)acetate (**3**).

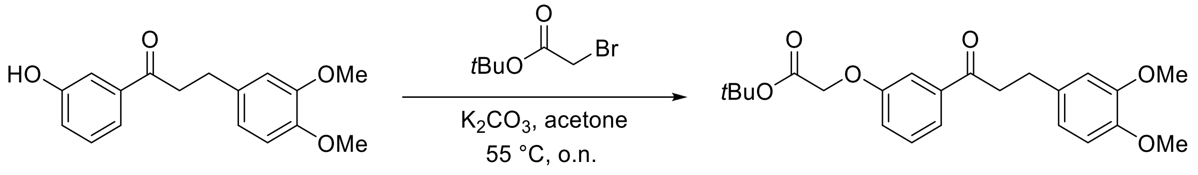


**2** **3**

A two-necked round-bottom flask was charged with **2** (13.4 g, 46.8 mmol) and anhydrous $K_2CO_3$ (9.7 g, 70.2 mmol, 1.5 eq.). Anhydrous acetone (133 mL, 0.35M) was added to the open flask, a reflux was installed, and the reactor was purged with vacuum-$N_2$ cycles. The solution was cooled down to 0 °C and *tert*-butyl bromoacetate (7.6 mL, 51.5 mmol, 1.1 eq.) was added dropwise. Then, the reaction was heated up to 55 °C and was left to stir overnight. The reaction was cooled down to room temperature, filtered and washed twice with acetone (2 x 1.2V). The solvent was removed under reduced pressure; the residue was dried in the vacuum line and afterwards stripped once with methanol. The crude was dissolved in 3.5V of methanol and the solution was heated to 85 °C, until complete dissolution. Then, water was added dropwise until the solution became turbid. The solution was warmed up again to 95 °C until complete dissolution, and then slowly cooled down to 0 °C. The obtained solid was filtrated, washed twice with cyclohexane (2 x 1.3V), and dried in a desiccator overnight. Compound **3** was obtained as a white solid (16.4 g, 90%). Spectroscopic data were in excellent agreement the reported in the literature.[1] . **$^1$H NMR (400 MHz, $CDCl_3$)** δ 7.56 (ddd, *J* = 7.7, 1.5, 0.9 Hz, 1H), 7.46 (dd, *J* = 2.7, 1.5 Hz, 1H), 7.36 (t, *J* = 7.9 Hz, 1H), 7.12 (ddd, *J* = 8.2, 2.7, 0.9 Hz, 1H), 6.82 – 6.74 (m, 3H), 4.56 (s,

2H), 3.86 (s, 3H), 3.85 (s, 3H), 3.25 (dd, *J* = 8.3, 6.9 Hz, 2H), 3.00 (dd, *J* = 8.3, 6.9 Hz, 2H), 1.48 (s, 9H) ppm. **$^{13}$C NMR (101 MHz, CDCl$_3$)** δ 199.0, 167.8, 158.3, 149.0, 147.5, 138.3, 133.9, 129.8, 121.6, 120.3, 120.2, 113.2, 111.9, 111.4, 82.7, 65.8, 56.0, 55.9, 40.9, 29.9, 28.1 ppm. **M.p.** = 51 – 54 °C.

2.4 Synthesis of *tert*-butyl *(R)*-2-(3-(3-(3,4-dimethoxyphenyl)-1-hydroxypropyl)phenoxy)acetate (4)

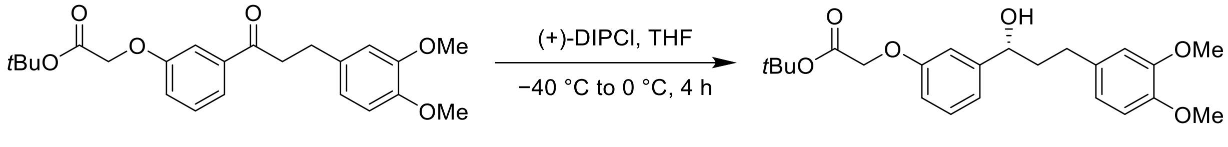


**3** **4**

A two-necked round-bottom flask was charged with **3** (16.3 g, 40.6 mmol), equipped with an addition funnel, and purged with vacuum-$N_2$ cycles. Then, anhydrous THF (87 mL, 0.47M) was added and the solution was cooled down to −40 °C. (+)-DIP-Chloride (44.0 mL, 70.3 mmol, 1.75 eq.) was charged to the addition funnel and added dropwise over 15 minutes. The solution was kept under stirring at the same temperature for 2 h. The reaction was gradually warmed up to 0 °C over 2 h. Then, water (25 mL, 1.5V) was added to the reaction and it was left to stir for 15 minutes. The solution was diluted with water and EtOAc, the layers were separated, and the aqueous phase was extracted thrice with EtOAc. The combined organic layers were dried over $MgSO_4$, filtered and evaporated. The crude was purified by column chromatography, with gradient to 7:3 cyclohexane:EtOAc. Compound **4** was obtained as a colorless oil (16.1 g, 97%). The enantiomeric excess was determined by chiral HPLC (96% ee). **$^{1}$H NMR (400 MHz, CDCl$_3$)** δ 7.27 – 7.21 (m, 1H), 6.94 (ddt, *J* = 7.4, 1.4, 0.7 Hz, 1H), 6.91 (dd, *J* = 2.7, 1.5 Hz, 1H), 6.81 – 6.75 (m, 2H), 6.72 (d, *J* = 2.0 Hz, 1H), 6.70 (d, *J* = 0.6 Hz, 1H), 4.64 (ddd, *J* = 7.6, 5.4, 2.5 Hz, 1H), 4.50 (s, 2H), 3.84 (s, 6H), 2.73 – 2.53 (m, 2H), 2.08 – 1.91 (m, 2H), 1.47 (s, 9H) ppm. **$^{13}$C NMR (101 MHz, CDCl$_3$)** δ 168.1, 158.2, 148.9, 147.2, 146.6, 134.4, 129.6, 120.3, 119.2, 113.6, 112.3, 111.8, 111.3, 82.4, 73.7, 65.7, 56.0, 55.9, 40.7, 31.7, 28.1 ppm. **HPLC:** Chiralpak IA, heptane/EtOH 80:20, 0.5 mL/min, λ = 210 nm. $t_{maj}$ = 15.2 min, $t_{min}$ = 16.7 min.

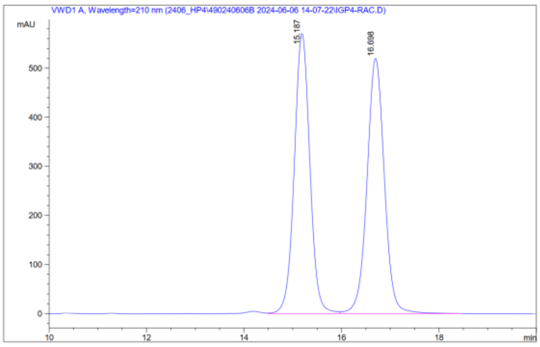


| # | Meas. R | Pea | Width | Area | Height | Area % |
|---|---|---|---|---|---|---|
| 1 | 15.187 | VV | 0.344 | 12701.440 | 570.923 | 49.636 |
| 2 | 16.698 | VB | 0.381 | 12887.743 | 519.986 | 50.364 |

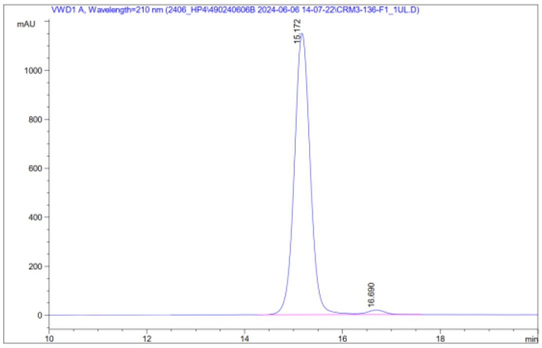


| # | Meas. R | Pea | Width | Area | Height | Area % |
|---|---|---|---|---|---|---|
| 1 | 15.172 | VV | 0.348 | 25837.814 | 1.150e3 | 98.076 |
| 2 | 16.690 | VB | 0.399 | 506.889 | 18.910 | 1.924 |

**Supplementary Figure S2.** Chiral HPLC of racemic and enantiomerically enriched alcohol **4**.

2.5 Synthesis of 1-((9H-fluoren-9-yl)methyl)-2-((*R*)-1-(3-(2-(*tert*-butoxy)-2-oxoethoxy)phenyl)-3-(3,4-dimethoxyphenyl)propyl (*S*)-piperidine-1,2-dicarboxylate (**5**)

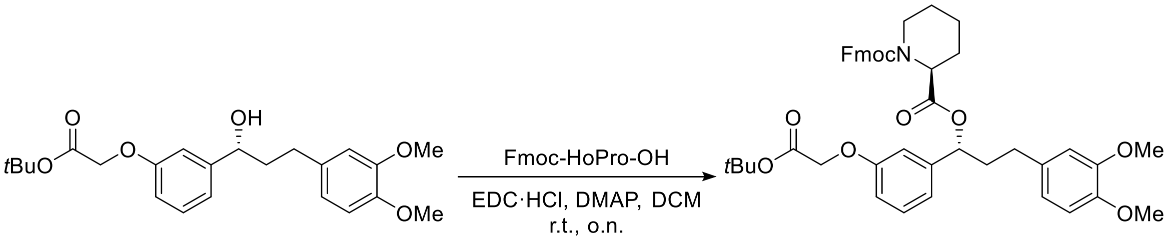


**4** **5**

A round bottom flask was charged with **4** (15.1 g, 37.5 mmol), Fmoc-HoPro-OH (14.5 g, 41.3 mmol, 1.1 eq.) and purged with vacuum-$N_2$ cycles. Then, anhydrous DCM (90 mL) was added. The solution was cooled down to 0 °C and a solution of EDC·HCl (8.6 g, 45.0 mmol, 1.2 eq.) in DCM (20 mL) was added dropwise. The reaction was kept at 0 °C for 15 min, then warmed up to room temperature and left to stir overnight. The reaction was diluted with DCM (2.4V) and washed with HCl 1.0M (3.2V). The aqueous phase was extracted thrice with DCM, the combined organic layers were dried with $MgSO_4$, filtered and evaporated. The crude was purified by column chromatography, with gradient to 8:2 cyclohexane:EtOAc. Compound **5** was obtained as a white solid (24.0 g, 87%). **$^1$H NMR (400 MHz, $CDCl_3$)** Mixture of rotamers, δ 7.80 – 7.74 (m, 1H), 7.71 (d, *J* = 7.6 Hz, 1H), 7.59 (t, *J* = 8.0 Hz, 1H), 7.51 – 7.14 (m, 6H), 6.98 – 6.92 (m, 1H), 6.89 (t, *J* = 2.0 Hz, 1H), 6.84 – 6.75 (m, 1H), 6.73 (d, *J* = 8.2 Hz, 1H), 6.67 – 6.55 (m, 2H), 5.82 – 5.69 (m, 1H), 5.02* (d, *J* = 5.5 Hz, 1H major rotamer), 4.88* (d, *J* = 5.6 Hz, 1H minor rotamer), 4.49 (s, 1H), 4.47 – 4.23 (m, 3H), 4.11 (q, *J* = 11.8, 9.4 Hz, 1H), 3.84 (s, 3H), 3.80 (d, *J* = 7.8 Hz, 3H), 3.14* (td, *J* = 13.1, 3.0 Hz, 1H major rotamer), 2.99* (td, *J* = 13.3, 3.0 Hz, 1H minor rotamer), 2.62 – 2.39 (m, 1H), 2.38 – 2.27 (m, 1H), 2.25 – 2.12 (m, 1H), 2.08 – 1.93 (m,

1H), 1.72 (dtd, *J* = 17.9, 9.6, 8.3, 4.9 Hz, 3H), 1.46 (s, 9H), 1.33 – 1.15 (m, 1H) ppm. **$^{13}$C NMR (101 MHz, $CDCl_3$)** δ 171.1, 168.0, 158.2, 156.5, 148.9, 147.4, 144.2, 144.0, 141.8, 141.4, 133.6, 129.8, 127.8, 127.2, 125.2, 120.1, 119.8, 114.0, 113.4, 111.8, 111.4, 82.5, 76.3, 67.9, 65.8, 56.0, 55.9, 54.7, 47.3, 42.1, 38.2, 31.2, 28.1, 26.9, 24.9, 20.9 ppm. **M.p.** = 49 – 52 °C

2.6 Synthesis of (*R*)-1-(3-(2-(*tert*-butoxy)-2-oxoethoxy)phenyl)-3-(3,4-dimethoxyphenyl)propyl (*S*)-piperidine-2-carboxylate (**6**).

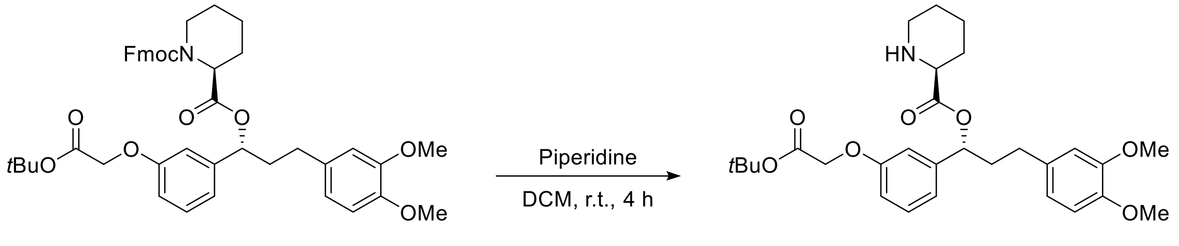


**5** **6**

A round-bottom flask was charged with **5** (7.5 g, 10.2 mmol), purged with vacuum-$N_2$ cycles and dissolved in anhydrous DCM (0.08M, 120 mL). The solution was cooled down to 0 °C and piperidine (7.0 mL, 71.2 mmol, 7 eq.) was added dropwise. The reaction was kept at 0 °C for 15 min, then warmed up to room temperature and left to stir for 4 h. The reaction was cooled down to 0 °C and 120 mL of a solution of saturated $NH_4Cl_{(aq.)}$ were slowly added. The solution was stirred at room temperature for 10 min. Then, the layers were separated, and the organic phase was washed five times with saturated $NH_4Cl_{(aq.)}$, once with $H_2O$ and once with brine. Afterwards, the organic phase was dried with $MgSO_4$, filtered and concentrated. Upon dried, the reaction was left the minimum amount of time possible heating in the rotavapor to minimize epimerization (4% of epimer was observed), and then further dried in the vacuum line. The crude was purified by column chromatography, with gradient to 2:8 cyclohexane:EtOAc, effectively removing the epimer traces. The product was obtained as a colorless oil (3.9 g, 75%). **$^1$H NMR (400 MHz, $CDCl_3$)** δ 7.23 (t, 1H), 6.93 (dt, *J* = 7.7, 1.2 Hz, 1H), 6.88 (t, *J* = 2.1 Hz, 1H), 6.81 – 6.74 (m, 2H), 5.75 (dd, *J* = 7.9, 5.6 Hz, 1H), 4.49 (d, *J* = 0.8 Hz, 2H), 3.84 (s, 3H), 3.83 (s, 3H), 3.35 (dd, *J* = 9.9, 3.2 Hz, 1H), 3.05 (dt, *J* = 11.9, 3.7 Hz, 1H), 2.69 – 2.46 (m, 3H), 2.29 – 2.13 (m, 1H), 2.10 – 1.97 (m, 3H), 1.84 – 1.73 (m, 1H), 1.66 – 1.51 (m, 2H), 1.47 (s, 9H), 1.24 (td, *J* = 7.2, 1.1 Hz, 1H) ppm. **$^{13}$C NMR (101 MHz, $CDCl_3$)** δ 172.9, 168.0, 158.1, 148.9, 147.4, 142.0, 133.7, 129.7,

120.2, 119.9, 113.8, 113.2, 111.7, 111.4, 82.5, 75.5, 65.8, 58.8, 56.0, 55.9, 45.8, 38.1, 31.4, 29.4, 28.1, 25.9, 24.3 ppm.

2.7 Synthesis of (*R*)-1-(3-(2-(tert-butoxy)-2-oxoethoxy)phenyl)-3-(3,4-dimethoxyphenyl)propyl (*S*)-1-((*S*)-2-(3,4,5-trimethoxyphenyl)butanoyl)piperidine-2-carboxylate (7)

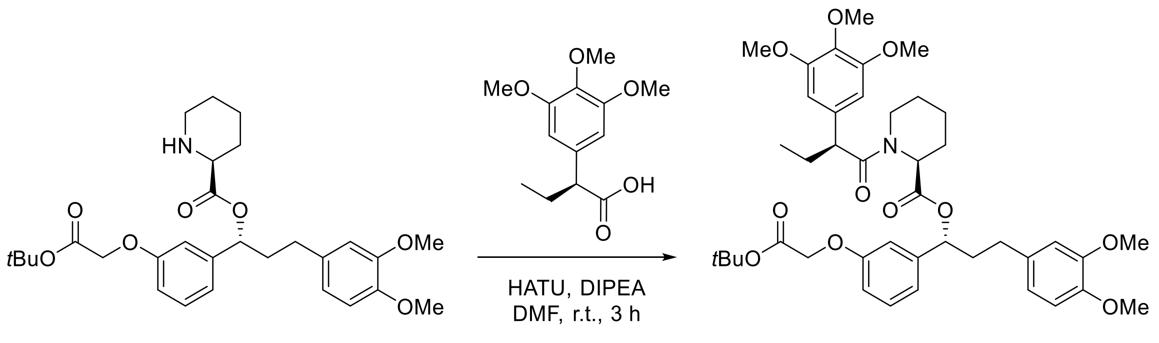


**6** **7**

Compound **6** (2.55 g, 4.96 mmol) and (*S*)-2-(3,4,5-trimethoxyphenyl)butanoic acid (1.33 g, 5.21 mmol, 1.05 eq.) were charged in a round-bottom flask, purged with vacuum-$N_2$ cycles and dissolved in anhydrous DMF (0.1M, 50 mL). The solution was cooled down to 0 °C and DIPEA (3.5 mL, 19.9 mmol, 4.0 eq.) was added dropwise. After stirring for 10 minutes, the flask was opened to air and HATU (2.08 g, 5.46 mmol, 1.5 eq.) was added in a solid manner. The reaction was purged again with vacuum-$N_2$ cycles and was left to stir at room temperature for 3 h under a $N_2$ atmosphere. The solution was diluted with DCM and cold water, the layers were separated, and the aqueous phase was extracted thrice with DCM. The combined organic layers were dried with $MgSO_4$, filtered and evaporated to dryness under reduced pressure. The crude was purified by column chromatography, with gradient to 7:3 cyclohexane:EtOAc. Compound **7** was obtained as a white solid (3.29 g, 88%). **$^1$H NMR (400 MHz, $CDCl_3$)** δ 7.15 (t, *J* = 7.8 Hz, 1H), 6.79 – 6.73 (m, 3H), 6.67 – 6.58 (m, 3H), 6.41 (d, *J* = 2.3 Hz, 2H), 5.61 (dd, *J* = 7.9, 5.6 Hz, 1H), 5.49 – 5.44 (m, 1H), 4.52 (d, *J* = 1.3 Hz, 2H), 3.86 – 3.82 (m, 10H), 3.78 (s, 3H), 3.69 (s, 6H), 3.58 (dd, *J* = 7.8, 6.5 Hz, 1H), 2.80 (ddd, *J* = 14.9, 13.0, 3.0 Hz, 1H), 2.63 – 2.38 (m, 3H), 2.15 – 2.00 (m, 2H), 1.98 – 1.85 (m, 1H), 1.78 – 1.64 (m, 2H), 1.47 (s, 8H), 0.90 (t, *J* = 7.3 Hz, 3H) ppm. **$^{13}$C NMR (101 MHz, $CDCl_3$)** δ 172.7, 170.7, 168.1, 158.0, 153.3, 149.0, 147.4, 142.0, 136.8, 135.5, 133.6, 129.7, 120.3, 119.4, 113.9, 113.2, 111.8, 111.4, 105.1, 82.4, 75.9, 65.9, 60.9, 56.4, 56.1, 56.0, 56.0, 52.2, 50.9, 43.6, 38.3, 31.3, 28.5, 28.2, 27.0, 26.9, 25.5, 21.1, 12.7 ppm. **M.p.** = 42 °C – 45 °C

2.8 Synthesis of 2-(3-((*R*)-3-(3,4-dimethoxyphenyl)-1-(((*S*)-1-((*S*)-2-(3,4,5-trimethoxyphenyl)butanoyl)piperidine-2-carbonyl)oxy)propyl)phenoxy)acetic acid (**8**)

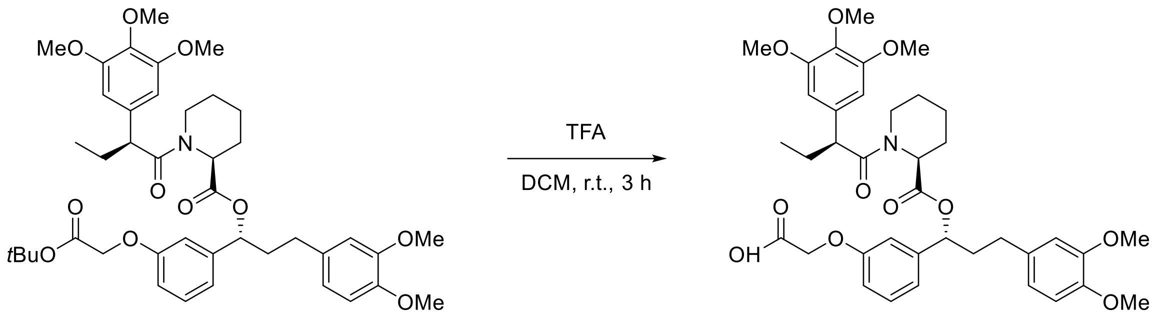


**7** **8**

Compound **7** (1.40 g, 1.87 mmol) was placed in a round-bottom flask and dissolved in anhydrous DCM (25 mL, 0.07M), under $N_2$ atmosphere. Then, TFA (4.3 mL, 30.0 eq.) was added dropwise. The reaction mixture was left to stir at room temperature for 3 h. After that time, the solvent was removed under reduced pressure, and the crude was purified by column chromatography, with gradient to 97:3 DCM:MeOH. Compound **8** was obtained as a white solid (1.27 g, 97%). **$^1$H NMR (400 MHz, $CDCl_3$)** δ 7.19 (dd, *J* = 8.3, 7.5 Hz, 1H), 6.84 (ddd, *J* = 8.3, 2.7, 0.9 Hz, 1H), 6.80 – 6.73 (m, 2H), 6.71 – 6.68 (m, 2H), 6.66 (s, 2H), 6.26 (s, 2H), 5.55 – 5.45 (m, 2H), 4.73 (d, *J* = 5.5 Hz, 7H), 3.85 (d, *J* = 3.6 Hz, 9H), 3.56 (s, 7H), 2.87 (td, *J* = 13.4, 3.2 Hz, 1H), 2.66 (ddd, *J* = 14.6, 9.6, 5.3 Hz, 1H), 2.55 (dt, *J* = 9.4, 6.9 Hz, 1H), 2.38 – 2.25 (m, 1H), 2.21 – 2.10 (m, 1H), 2.09 – 1.92 (m, 1H), 1.83 – 1.57 (m, 3H), 1.26 (d, *J* = 2.4 Hz, 4H) ppm. **$^{13}$C NMR (101 MHz, $CDCl_3$)** δ 173.8, 171.2, 170.2, 158.2, 153.3, 149.1, 147.5, 142.8, 136.6, 134.7, 133.4, 129.6, 120.3, 119.5, 115.9, 111.7, 111.4, 108.8, 104.9, 65.7, 60.9, 56.1, 56.0, 56.0, 52.4, 51.2, 43.6, 38.6, 31.7, 29.8, 28.3, 27.4, 25.3, 20.9, 12.6 ppm.

2.9 Synthesis of (1*R*)-3-(3,4-dimethoxyphenyl)-1-(3-(2-((6-((2-(2,6-dioxopiperidin-3-yl)-1,3-dioxoisoindolin-4-yl)oxy)hexyl)amino)-2-oxoethoxy)phenyl)propyl (2*S*)-1-((*S*)-2-(3,4,5-trimethoxyphenyl)butanoyl)piperidine-2-carboxylate, (**9, dTAG-13**)

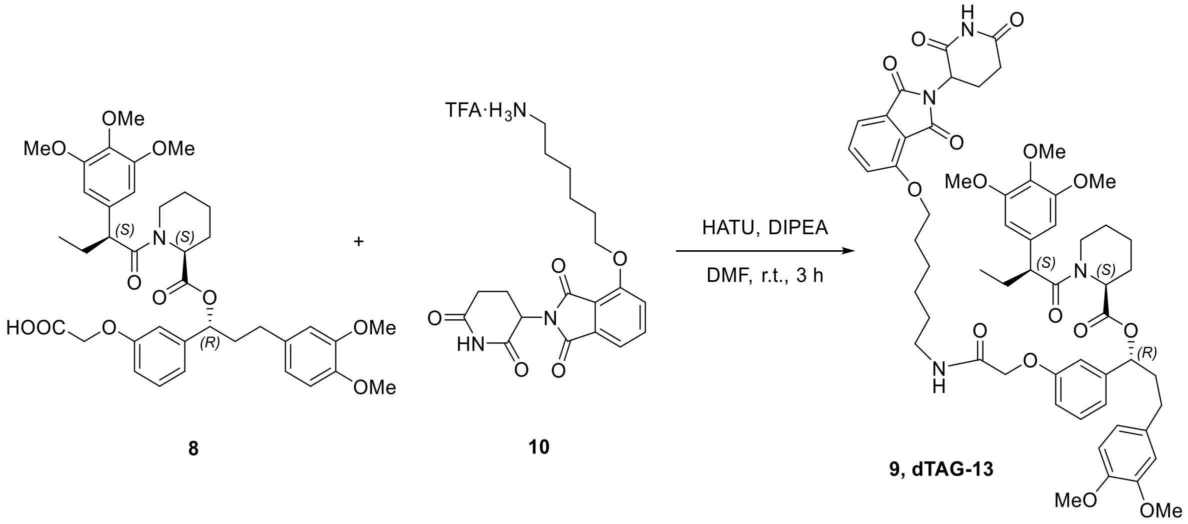


Compound **8** (1.0 g, 1.44 mmol, 1.0 eq.) and compound **10** (0.7 g, 1.44 mmol, 1.0 eq.)(prepared according to the reported procedure[2]) were charged in a round-bottom flask, purged with vacuum-$N_2$ cycles and dissolved in anhydrous DMF (0.96M, 15 mL). The solution was cooled down to 0 °C and DIPEA (1.0 mL, 5.77 mmol, 4.0 eq.) was added dropwise. After stirring for 10 minutes, the flask was opened to air and HATU (0.82 g, 2.16 mmol, 1.5 eq.) was added as solid The reaction was purged again with vacuum-$N_2$ and was left to stir at room temperature for 3 h under $N_2$ atmosphere. The solution was diluted with cold water and diethyl ether, the layers were separated, and the aqueous phase was extracted thrice with diethyl ether. The combined organic layers were washed once with brine, dried with $MgSO_4$, filtered and evaporated to dryness under reduced pressure. The crude was purified by reverse phase column chromatography, with gradient to 2:8 $NH_4HCO_2$: ACN. Compound **9** was obtained as a white solid (1.05 g, 70% yield, 94% purity by HPLC-MS). The product was further purified by reverse phase column chromatography with gradient to 2:8 $NH_4HCO_2$:ACN, obtaining dTAG-13 as a white solid (744 mg, 50% yield, 98% purity by HPLC-MS)[3]. **$^1$H NMR (400 MHz, $CDCl_3$)** δ 7.66 (t, *J* = 7.8 Hz, 1H), 7.44 (d, *J* = 7.3 Hz, 1H), 7.20 (d, *J* = 8.4 Hz, 2H), 6.82 – 6.74 (m, 3H), 6.70 – 6.61 (m, 3H), 6.42

[2] a) Harki, D. A.; Tang, J.; Moorthy, R.: Preparation of heteroaryl compounds that degrade kinases and uses thereof. WO2020247537, 2020. b) Bradner, J.; Buckley, D.; Winter, G.: Methods to induce targeted protein degradation through bifunctional molecules, WO2016105518, 2016.

[3] Nabet, B.; Roberts, J. M.; Buckley, D. L.; Paulk, J.; Dastjerdi, S.; Yang, A.; Leggett, A. L.; Erb, M. A.; Lawlor, M. A.; Souza, A.; Scott, T. G.; Vittori, S.; Perry, J. A.; Qi, J.; Winter, G. E.; Wong, K.-K.; Gray, N. S.; Bradner, J. E.: The dTAG system for immediate and target-specific protein degradation. *Nat. Chem. Biol.* **2018**, *14*, 431-441

(d, *J* = 3.8 Hz, 2H), 5.63 (dd, *J* = 8.2, 5.3 Hz, 1H), 5.47 (d, *J* = 5.5 Hz, 1H), 4.94 (dd, *J* = 12.1, 5.3 Hz, 1H), 4.48 (d, *J* = 5.5 Hz, 2H), 4.16 (tt, *J* = 6.4, 2.7 Hz, 2H), 3.85 (q, *J* = 3.5, 2.9 Hz, 8H), 3.78 (s, 2H), 3.68 (s, 5H), 3.36 (dq, *J* = 13.1, 6.7, 5.2 Hz, 2H), 2.90 – 2.65 (m, 4H), 2.51 (dddd, *J* = 38.6, 14.1, 9.1, 5.7 Hz, 2H), 2.31 (d, *J* = 12.6 Hz, 1H), 2.10 (tdd, *J* = 17.2, 14.3, 12.3, 7.6 Hz, 3H), 1.89 (dp, *J* = 21.0, 7.8, 6.7 Hz, 3H), 1.72 (p, *J* = 6.7, 6.2 Hz, 2H), 1.57 (dt, *J* = 23.9, 7.4 Hz, 3H), 1.43 (dt, *J* = 15.2, 6.9 Hz, 3H), 1.32 – 1.20 (m, 1H), 0.90 (t, *J* = 7.3 Hz, 2H) ppm. **$^{13}$C NMR (101 MHz, CDCl$_3$)** δ 172.8, 171.2, 170.7, 168.3, 168.2, 167.2, 165.8, 157.4, 156.7, 153.3, 148.9, 147.4, 142.4, 136.7, 136.6, 135.4, 133.9, 133.4, 129.9, 120.3, 119.9, 119.9, 119.0, 117.2, 115.8, 113.6, 113.2, 111.8, 111.4, 105.1, 75.7, 69.4, 67.5, 60.9, 56.4, 56.1, 56.0, 55.9, 52.1, 50.9, 49.2, 43.6, 39.1, 38.3, 31.5, 31.3, 29.5, 28.8, 28.4, 26.9, 26.6, 25.7, 25.4, 22.7, 21.0, 12.8, 12.6 ppm. **M.p.** = 95–97 °C.

Kinetex EVO C18 50x 4.6mm, 2.6um; Mobile phase: 10mM NH4HCO3 pH8 / ACN (95:5)--0.5min---(95:5)---6.5min----(0:100)---2min----(0:100) post run 1.5 min; Tª 40°C

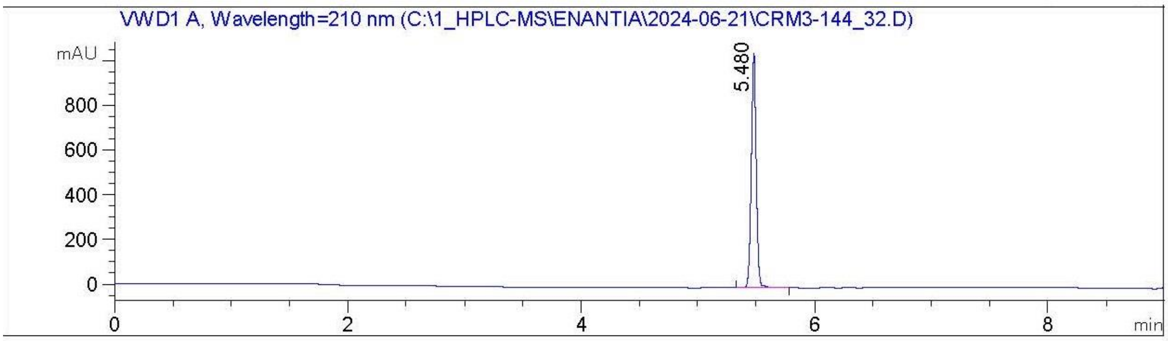


Signal: 2-> : VWD1 A, Wavelength=210 nm

| # | Meas. Ret. Time | Area | Area % |
|---|---|---|---|
| 1 | 5.480 | 3089.74 | 100.00 |

5.51 min

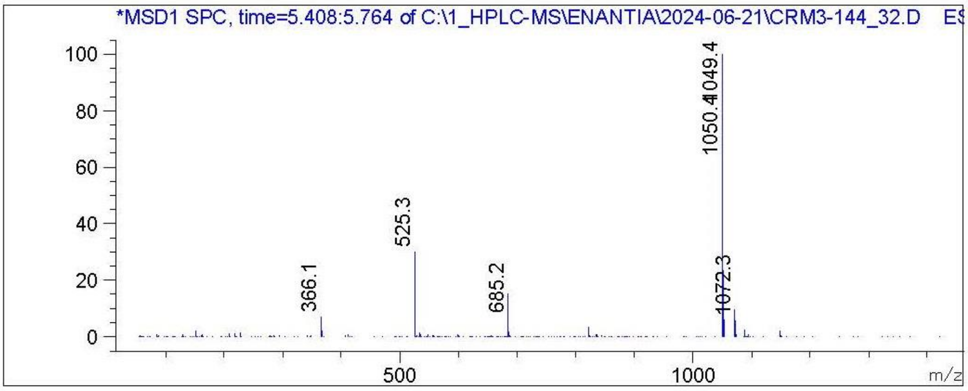


**Supplementary Figure S3.** Purity of compound **9** (**dTag-13**) by HPLC/Ms.

#### 3. Experimental procedures for the synthesis of dTAG-v1.

3.1 (R)-3-(3,4-dimethoxyphenyl)-1-(3-(2-((7-(((S)-1-((2S,4R)-4-hydroxy-2-(((S)-1-(4-(4-methylthiazol-5-yl)phenyl)ethyl)carbamoyl)pyrrolidin-1-yl)-3,3-dimethyl-1-oxobutan-2-yl)amino)-7-oxoheptyl)amino)-2-oxoethoxy)phenyl)propyl (S)-1-((S)-2-(3,4,5-trimethoxyphenyl)butanoyl)piperidine-2-carboxylate (**12, dTAG$^{v}$-1**).

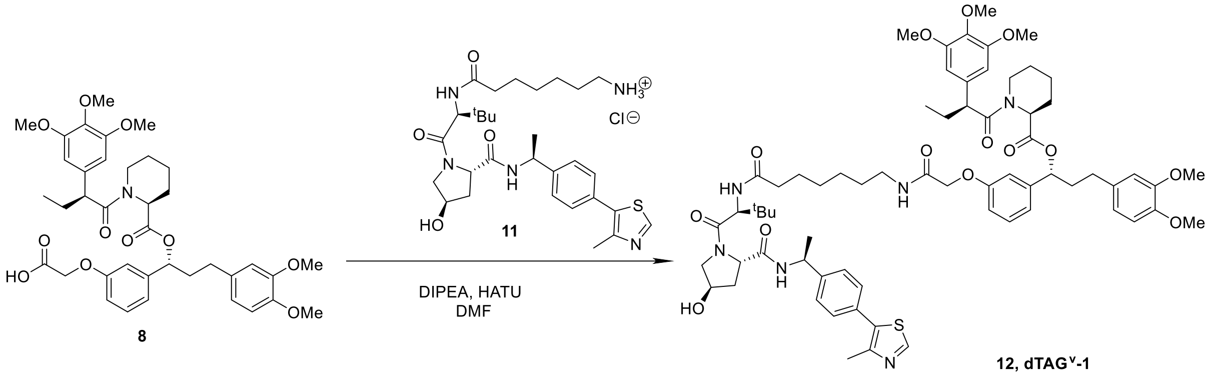


A round bottomed flask was charged with compound **8** (912.5 mg, 1.31 mmol, 1.0 eq.), compound **11** (800 mg, 1.31 mmol, 1.0 eq.)(prepared according to the reported procedure[4,5]), DIPEA (4 eq) and DMF (0,25M). The mixture was stirred at 0°C and HATU (550 mg, 1,1 eq) was added. The reaction was let to warm up to r.t. for 2-5 h (reaction completion usually achieved after 3h). The reaction was followed by TLC. When reaction completion was achieved, water and EtOAc were added and the aqueous layer was extracted with EtOAc (x3). The organic layers were combined, washed with brine (x2) and $CuSO_4$ (x2), dried over $MgSO_4$, filtered and the solvent was evaporated *in vacuo*. The crude was purified by flash column chromatography ($CH_2Cl_2$/MeOH 96:4). Compound **12** was obtained as a white solid (1.01 g, 62 % yield). The purity by HPLC was 95%. Compound **12** was dissolved in ACN (0,5M). Once the compound was dissolved, water was added in a proportion 2:1 Water/ACN. Trifluoroacetic acid (1,1 eq) was added to the solution and the mixture was lyophilized overnight.

**$^1$H NMR (400 MHz, DMSO)** Mixture of rotamers, the major rotamer is reported, δ: 8.98 (s, 1H), 8.38 (d, J = 7.8 Hz, 1H), 8.05 (q, J = 6.0, 4.5 Hz, 1H), 7.79 (d, J = 9.2 Hz, 1H), 7.43 (d, J = 8.0 Hz, 2H), 7.38 (d, J = 8.2 Hz, 2H), 7.21 – 7.14 (m, 1H), 7.00 – 6.89 (m, 1H), 6.88 – 6.83 (m, 2H), 6.72 (d, J = 2.0 Hz, 1H), 6.66 – 6.56 (m, 2H), 6.53 (s, 2H), 5.75 (s, 1H), 5.56 – 5.48 (dd, 1H), 5.31 – 5.25 (m, 1H), 5.10

---

[4] a) Galdeano, C.; Gadd, M. S.; Soares, P.; Scaffidi, S.; Van Molle, I.; Birced, I.; Hewitt, S.; Dias, D. M.; Ciulli, A.: Structure-Guided Design and Optimization of Small Molecules Targeting the Protein–Protein Interaction between the von Hippel–Lindau (VHL) E3 Ubiquitin Ligase and the Hypoxia Inducible Factor (HIF) Alpha Subunit with in Vitro Nanomolar Affinities. *J. Med. Chem.* **2014**, *57*, 8657-8663. B)

[5] Nabet, B., Ferguson, F.M., Seong, B.K.A. *et al.* Rapid and direct control of target protein levels with VHL-recruiting dTAG molecules. *Nat. Commun.*, **2020**, *11*, 4687.

(d, J = 3.5 Hz, 1H), 4.92 (p, J = 7.7 Hz, 1H), 4.56 – 4.38 (m, 4H), 4.28 (d, J = 3.4 Hz, 1H), 4.06 – 3.96 (m, 1H), 3.90 – 3.83 (m, 1H), 3.74 (s, 3H), 3.72 (s, 1H), 3.71 (s, 2H), 3.70 (s, 3H), 3.63 (s, 2H), 3.59 (s, 6H), 3.57 (s, 3H), 3.10 (q, J = 6.8 Hz, 2H), 2.70 – 2.59 (m, 1H), 2.50 (p, J = 1.9 Hz, 2H), 2.45 (s, 3H), 2.43 – 2.11 (m, 2H), 2.07 – 1.85 (m, 2H), 1.84 – 1.74 (m, 1H), 1.70 – 1.41 (m, 2H), 1.37 (d, J = 7.0 Hz, 3H), 1.21 (d, 6H), 0.93 (s, 9H), 0.81 (t, J = 7.2 Hz, 3H) ppm. **$^{13}$C NMR (101 MHz, DMSO)** Mixture of rotamers, the major rotamer is reported, δ 172.2, 172.0, 170.6, 170.3, 169.6, 167.3, 157.6, 152.7, 151.5, 148.6, 147.8, 147.1, 144.7, 141.9, 135.9, 135.5, 133.1, 131.1, 129.7, 129.5, 128.8, 126.4, 126.3, 120.0, 118.6, 113.8, 112.9, 112.1, 111.8, 105.1, 104.8, 75.0, 68.8, 67.0, 60.0, 59.8, 58.6, 56.3, 56.3, 55.9, 55.5, 55.5, 55.3, 51.6, 48.8, 47.7, 42.9, 38.3, 37.7, 37.6, 35.2, 34.9, 30.6, 29.1, 28.4, 28.1, 27.8, 26.4, 26.4, 26.1, 25.4, 24.9, 22.5, 20.5, 16.0, 12.4, 12.3 ppm. The analytical data for this compound was in excellent agreement with the reported data.[5]

```
HPLC8, Kinetex C18 50 x 4.6 mm, 2.6 um (COL-HP-136)
HCOOH0.1%:ACN (95:5)---0.5 min---(95:5)---6.5 min--(0:100)---2
min---(0:100)
Aprox: 1mg/ml  ACN
```

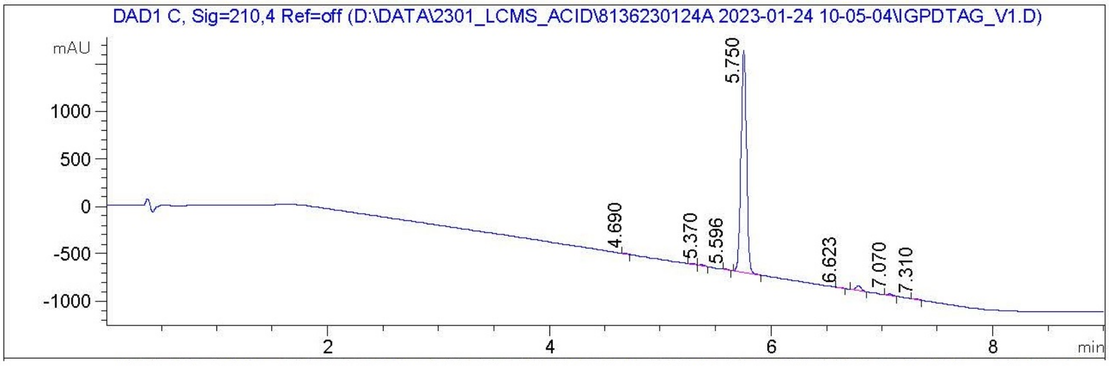


Signal:    1-> DAD1 C, Sig=210,4 Ref=off

| # | Meas. Ret. Time | Peak Type | Area | Area % |
|---|---|---|---|---|
| 1 | 4.690 | MM | 11.03 | 0.13 |
| 2 | 5.290 | MF | 23.60 | 0.28 |
| 3 | 5.370 | FM | 38.35 | 0.45 |
| 4 | 5.596 | MM | 12.81 | 0.15 |
| 5 | 5.750 | MM | 8111.11 | 95.81 |
| 6 | 6.623 | MM | 7.49 | 0.09 |
| 7 | 6.790 | MM | 188.29 | 2.22 |
| 8 | 7.070 | MM | 49.68 | 0.59 |
| 9 | 7.310 | MM | 23.53 | 0.28 |

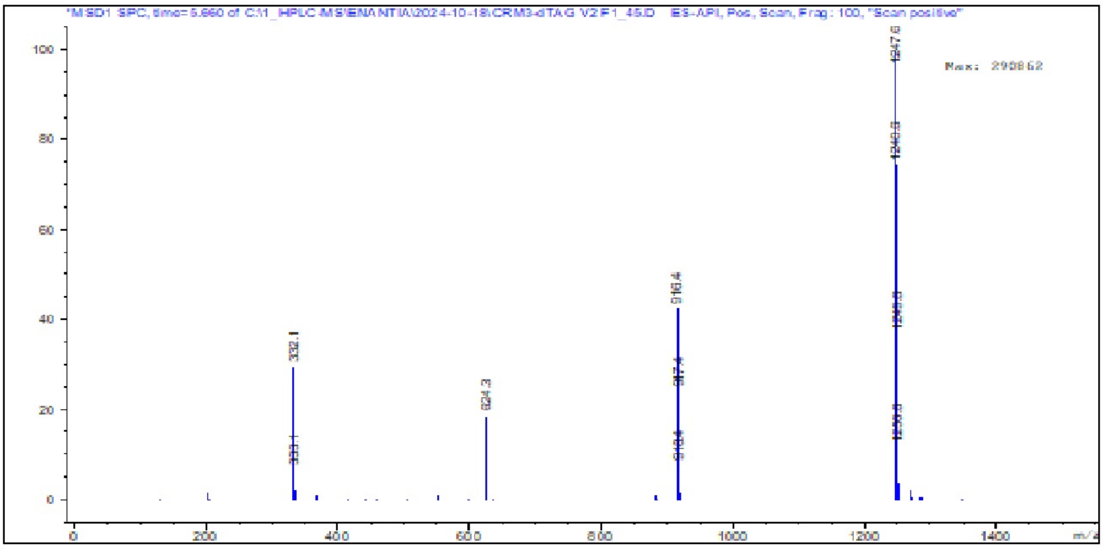


**Supplementary Figure S4.** Purity of compound **12** by HPLC/Ms.

#### 4. Experimental procedures of the synthesis of IGP002.

4.1 (1R)-3-(3,4-dimethoxyphenyl)-1-(3-(2-((6-((2-(2,6-dioxopiperidin-3-yl)-1,3-dioxoisoindolin-4-yl)amino)hexyl)amino)-2-oxoethoxy)phenyl)propyl (2S)-1-((S)-2-(3,4,5-trimethoxyphenyl)butanoyl)piperidine-2-carboxylate (**14, IGP002**).

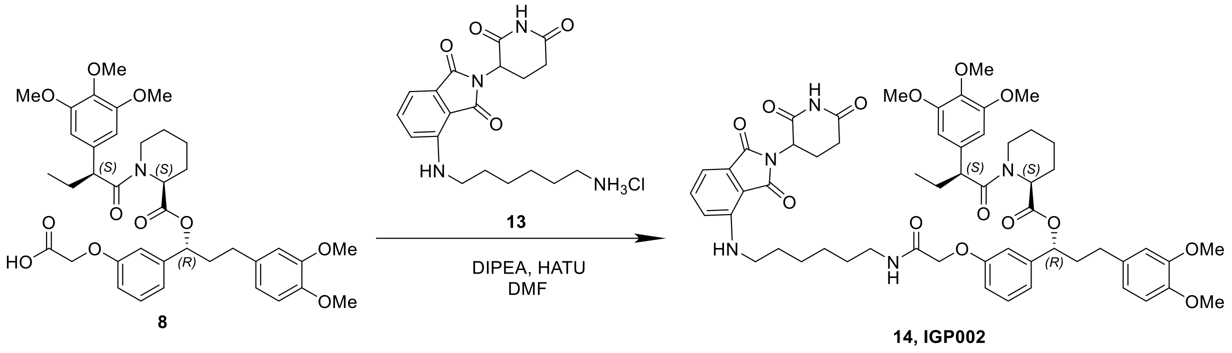


A round bottomed flask was charged with compound **8** (763.6 mg, 1.1 mmol, 1.0 eq.), compound **13** (450 mg, 1.1 mmol, 1.0 eq.)(prepared according to the reported procedure[6]), DIPEA (4 eq) and DMF (0,25M). The mixture was stirred at 0ºC and HATU (460 mg, 1,1 eq) was added. The reaction was let to warm up to r.t. for 2-5 h (reaction completion usually achieved after 3h). The reaction was followed by TLC. When reaction completion was achieved, water and EtOAc were added and the aqueous layer was extracted with EtOAc (x3). The organic layers were combined, washed with brine (x2) and $CuSO_4$ (x2), dried over $MgSO_4$, filtered and the solvent was evaporated *in vacuo*. The crude was purified by flash column chromatography ($CH_2Cl_2$/MeOH 97:3). Compound **14** was obtained as a yellow solid (751.3 mg, 65 % yield). The purity by HPLC was 91%.

**$^1$H NMR (400 MHz, DMSO)** Mixture of rotamers. The major rotamer is reported, δ 11.09 (s, 1H), 8.04 (t, J = 5.8 Hz, 1H), 7.56 (dd, J = 8.5, 7.0 Hz, 1H), 7.17 (t, J = 8.2 Hz, 1H), 7.06 (d, J = 8.6 Hz, 1H), 7.01 (d, J = 7.0 Hz, 1H), 6.85 – 6.79 (m, 3H), 6.72 (d, J = 2.0 Hz, 1H), 6.65 – 6.56 (m, 2H), 6.52 (s, 3H), 5.51 (dd, J = 8.4, 5.0 Hz, 1H), 5.27 (d, J = 5.6 Hz, 1H), 5.04 (dd, J = 12.8, 5.5 Hz, 1H), 4.45 (s, 2H), 4.06 – 3.94 (m, 1H), 3.86 (t, J = 7.1 Hz, 1H), 3.57 (d, J = 7.5 Hz, 8H), 3.29 – 3.19 (m, 2H), 3.11 (q, J = 6.7 Hz, 2H), 2.88 (ddd, J = 17.3, 13.5, 5.4 Hz, 1H), 2.65 – 2.54 (m, 1H), 2.41 – 2.30 (m, 1H), 2.16 (d, J = 12.4 Hz, 1H), 1.98 – 1.83 (m, 2H), 1.63 (d, J = 9.3 Hz, 1H), 1.56 (dq, J = 13.9, 7.1, 6.7 Hz, 3H), 1.48 – 1.32 (m, 2H), 1.26 (d, J = 19.4 Hz, 5H), 0.80 (t, J = 7.3 Hz, 3H) ppm. **$^{13}$C NMR (101 MHz, DMSO)** Mixture of rotamers. The major rotamer is reported, δ 172.8, 172.2, 170.3, 170.1, 169.0, 167.4, 167.3, 157.6, 152.6, 148.6, 147.1, 146.4, 141.9, 136.3, 135.9, 135.5, 133.1, 132.2, 129.5, 120.0, 118.6,

[6] Sun, Z.; Deng, B.; Yang, Z.; Mai, R.; Huang, J.; Ma, Z.; Chen, T.; Chen, J.: Discovery of pomalidomide-based PROTACs for selective degradation of histone deacetylase 8. *Eur. J. Med. Chem.* **2022**, *239*, 114544.

117.2, 113.8, 112.8, 112.1, 111.8, 110.4, 109.0, 105.0, 75.0, 67.0, 59.8, 55.9, 55.5, 55.5, 55.3, 51.8, 48.8, 48.5, 42.9, 41.8, 38.3, 37.6, 31.0, 30.6, 29.1, 28.6, 28.1, 26.4, 26.1, 26.0, 24.9, 22.2, 20.5, 12.3 ppm.

HPLC8, Kinetex C18 50 x 4.6 mm, 2.6 um (COL-HP-136)
HCOOH0.1%:ACN (95:5)---0.5 min---(95:5)---6.5 min--(0:100)---2min---(0:100)
1mg-mL ACN

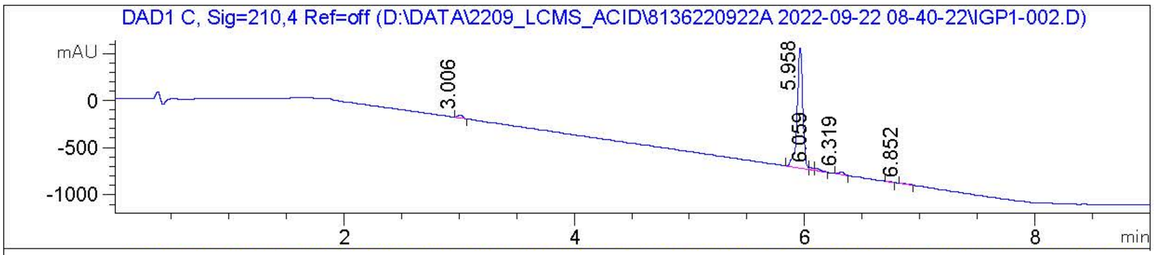


Signal: 1->: DAD1 C, Sig=210,4 Ref=off

| # | Meas. Ret | Peak | Width | Area | Response |
|---|---|---|---|---|---|
| 1 | 3.006 | MM | 0.045 | 93.461 | 2.019 |
| 2 | 5.958 | MF | 0.055 | 4.258e3 | 91.957 |
| 3 | 6.059 | MF | 0.046 | 55.993 | 1.209 |
| 4 | 6.106 | FM | 0.062 | 83.098 | 1.795 |
| 5 | 6.319 | MM | 0.051 | 87.579 | 1.892 |
| 6 | 6.732 | MM | 0.047 | 15.456 | 0.334 |
| 7 | 6.852 | MM | 0.091 | 36.824 | 0.795 |

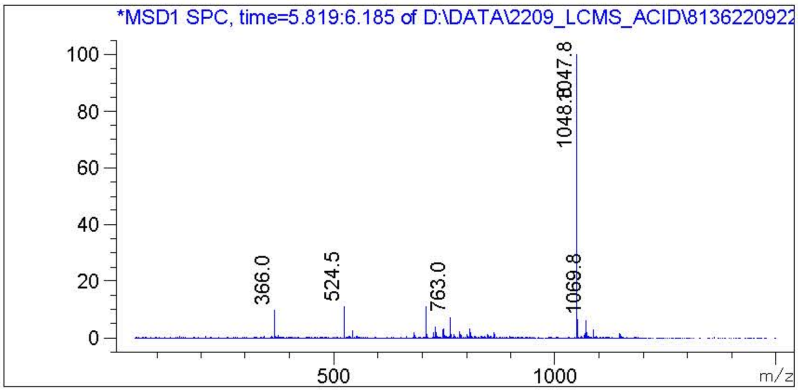


**Supplementary Figure S5. Purity of compound 14 by HPLC/Ms.**

### 5. $^{1}$H and $^{13}$C NMR spectra

#### 5.1. $^{1}$H and $^{13}$C NMR spectra of 1

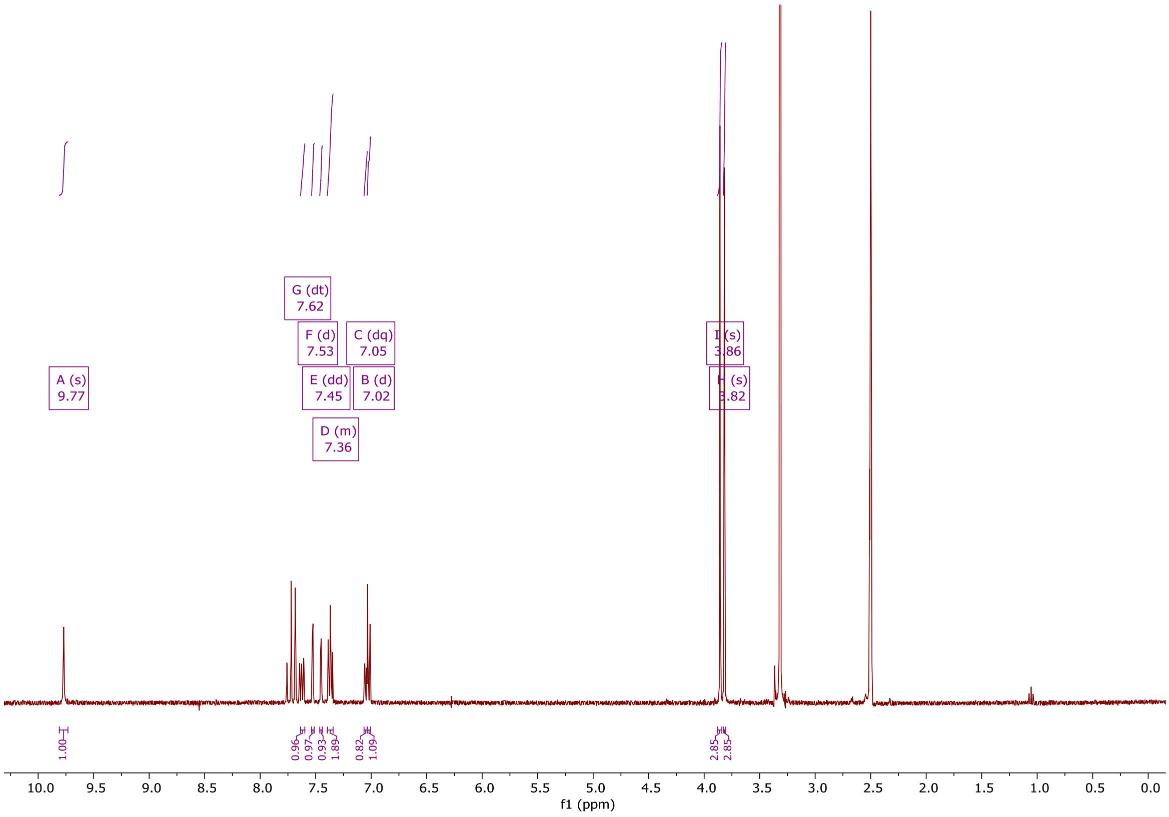


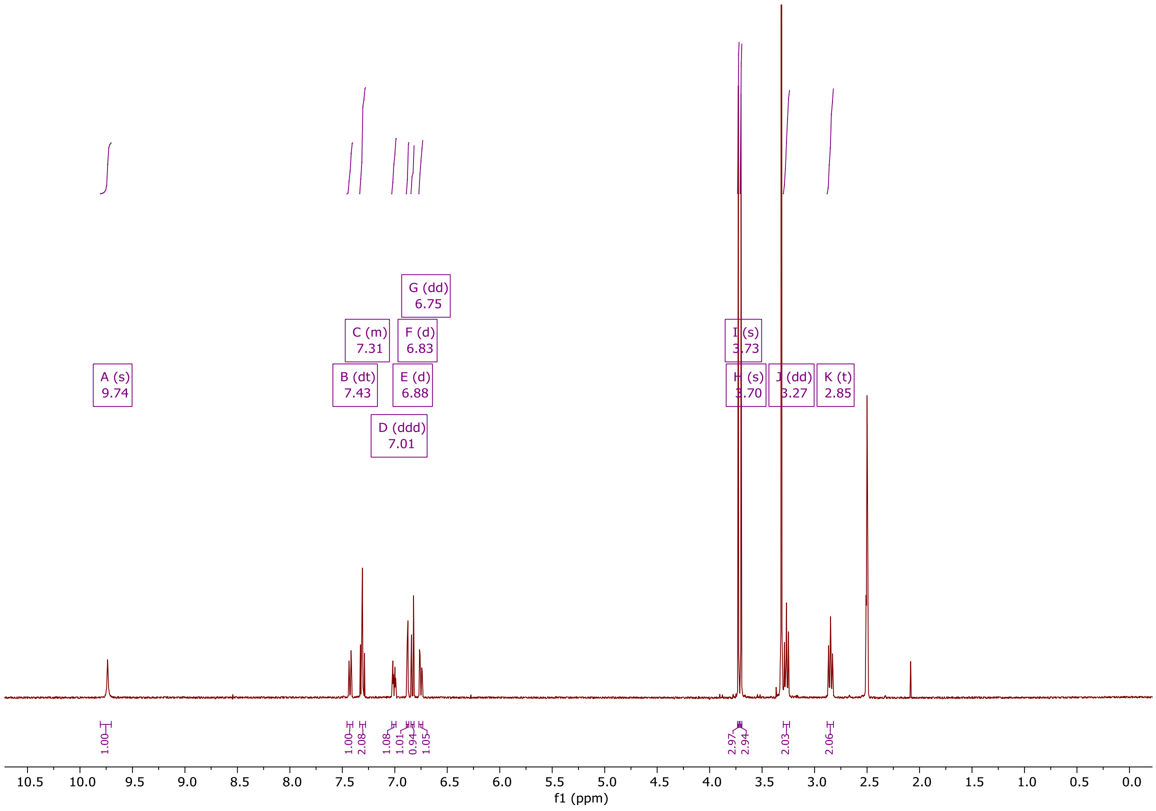

#### 5.2. $^{1}H$ and $^{13}C$ NMR spectra of 3

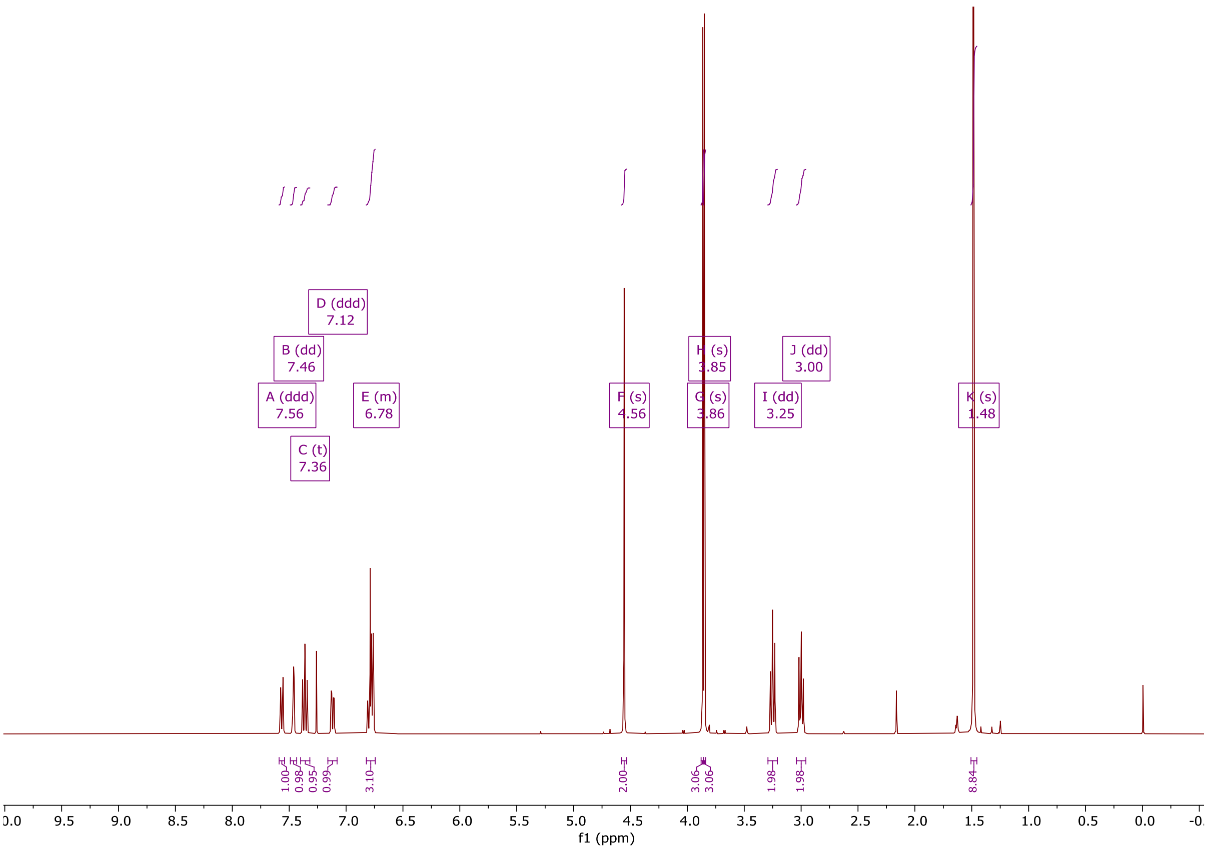


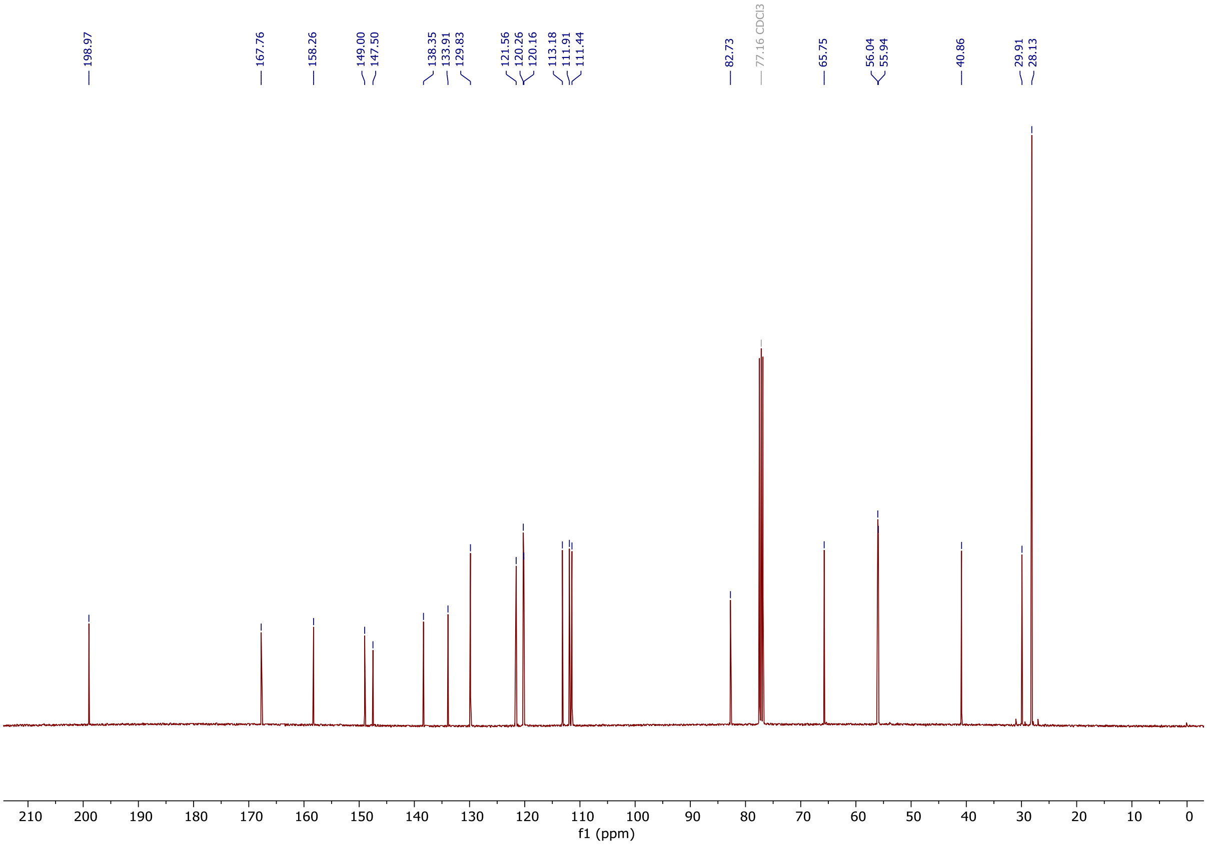

#### 5.3. [1]H and [13]C NMR spectra of 4

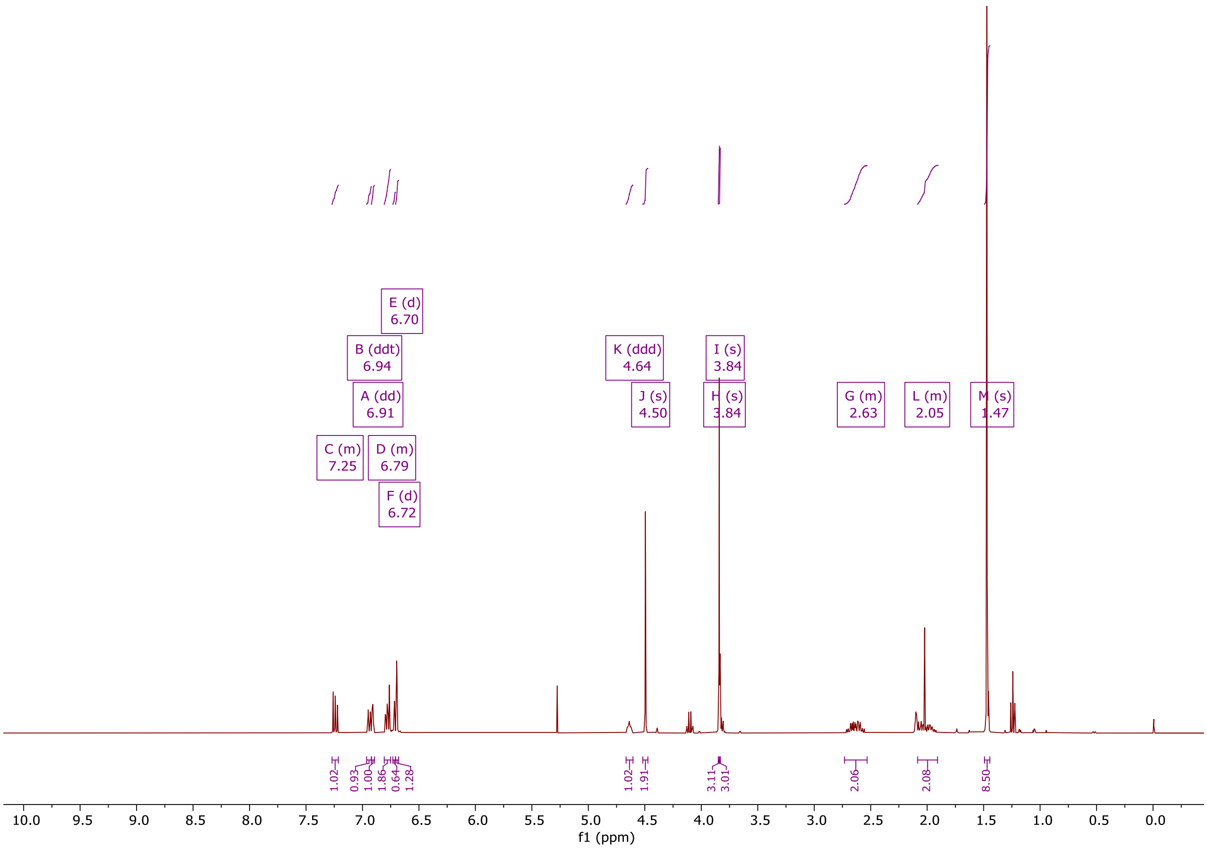


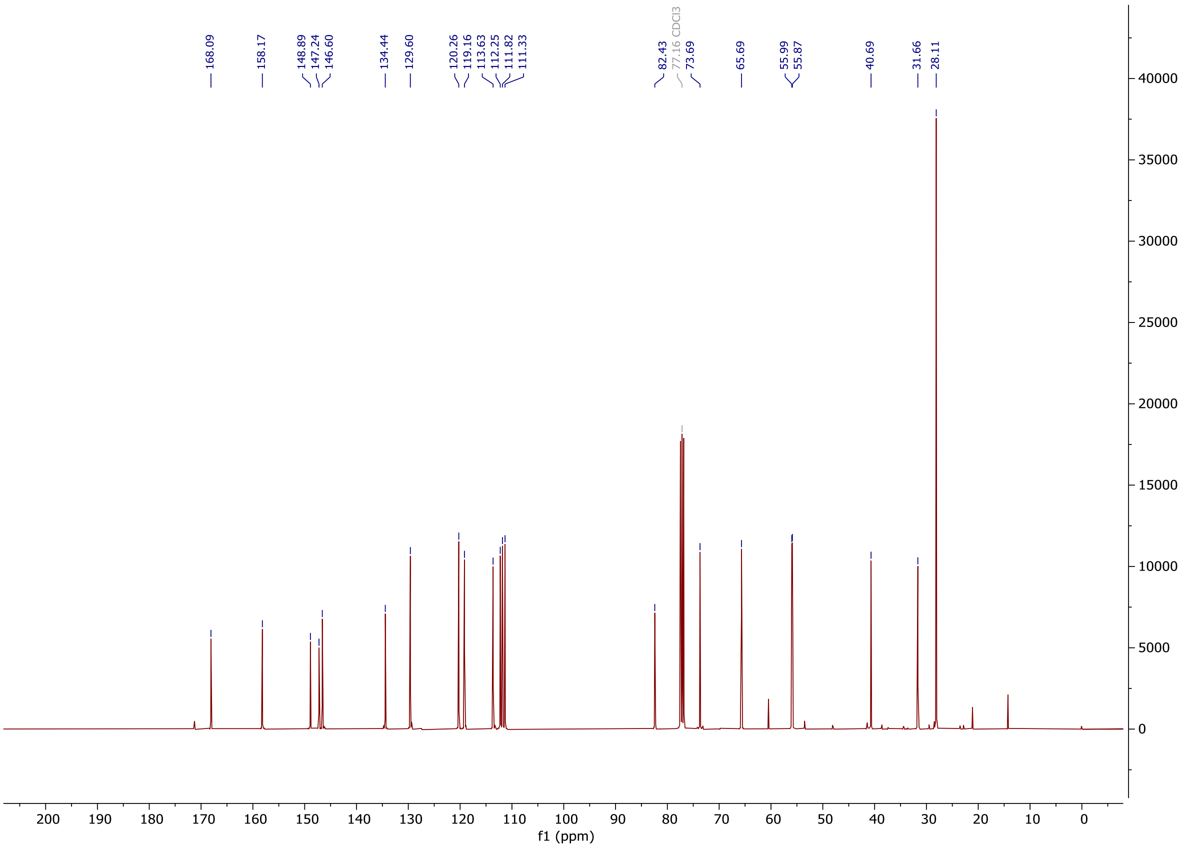

5.4. $^{1}$H and $^{13}$C NMR spectra of 5

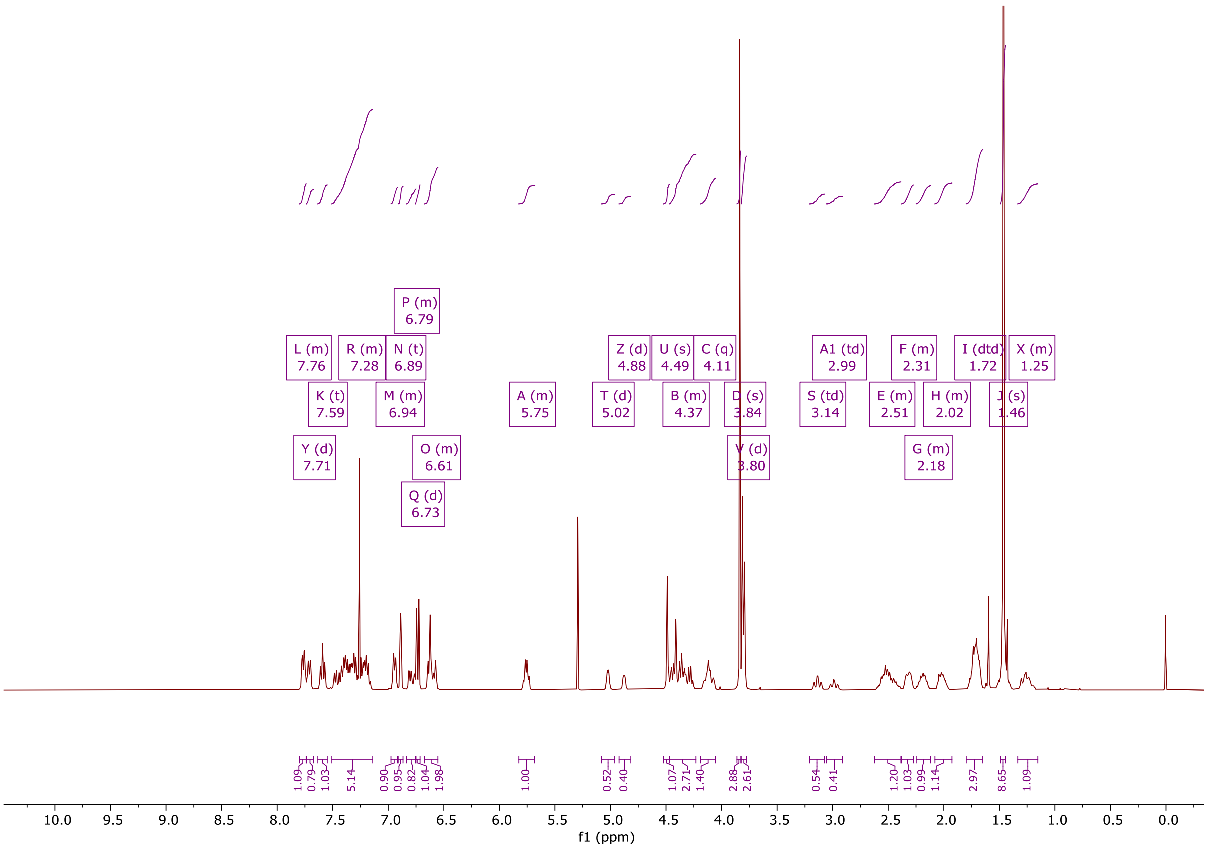


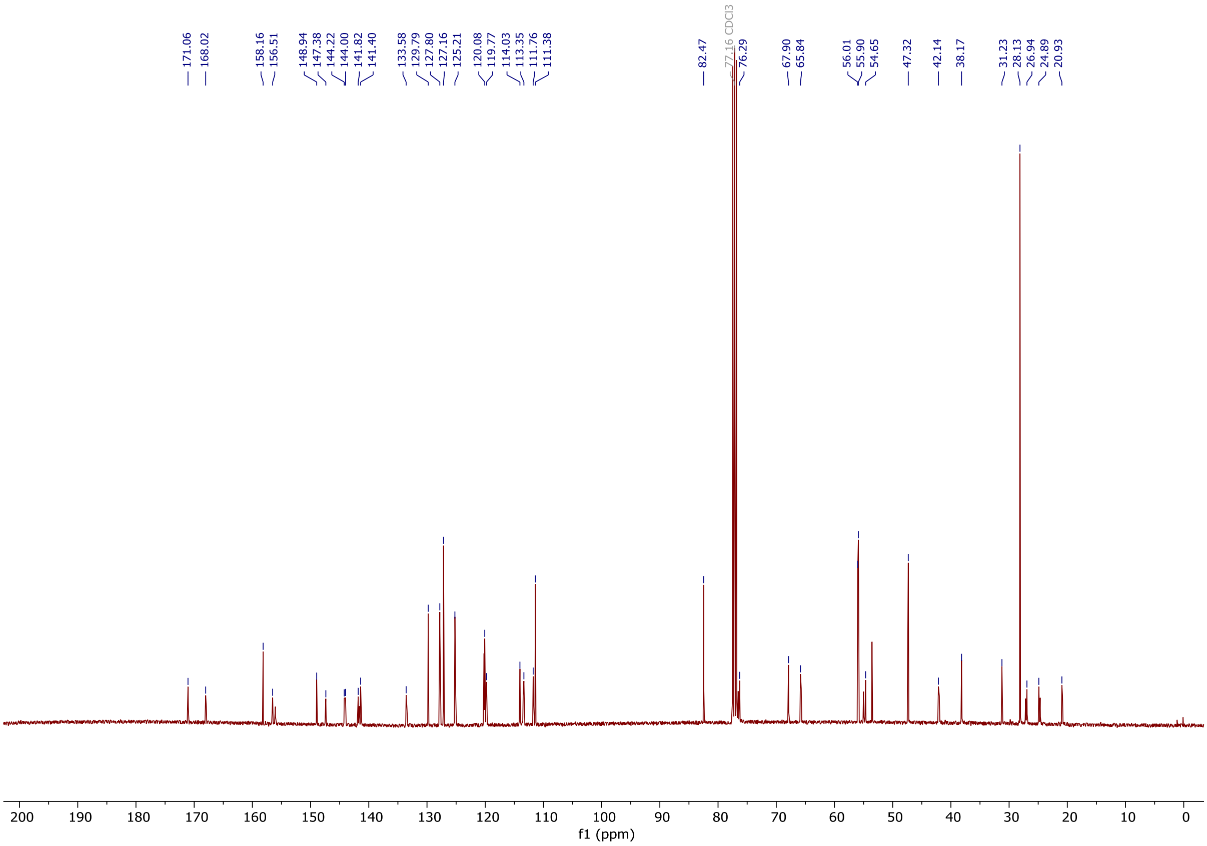

5.5. $^{1}$H and $^{13}$C NMR spectra of 6

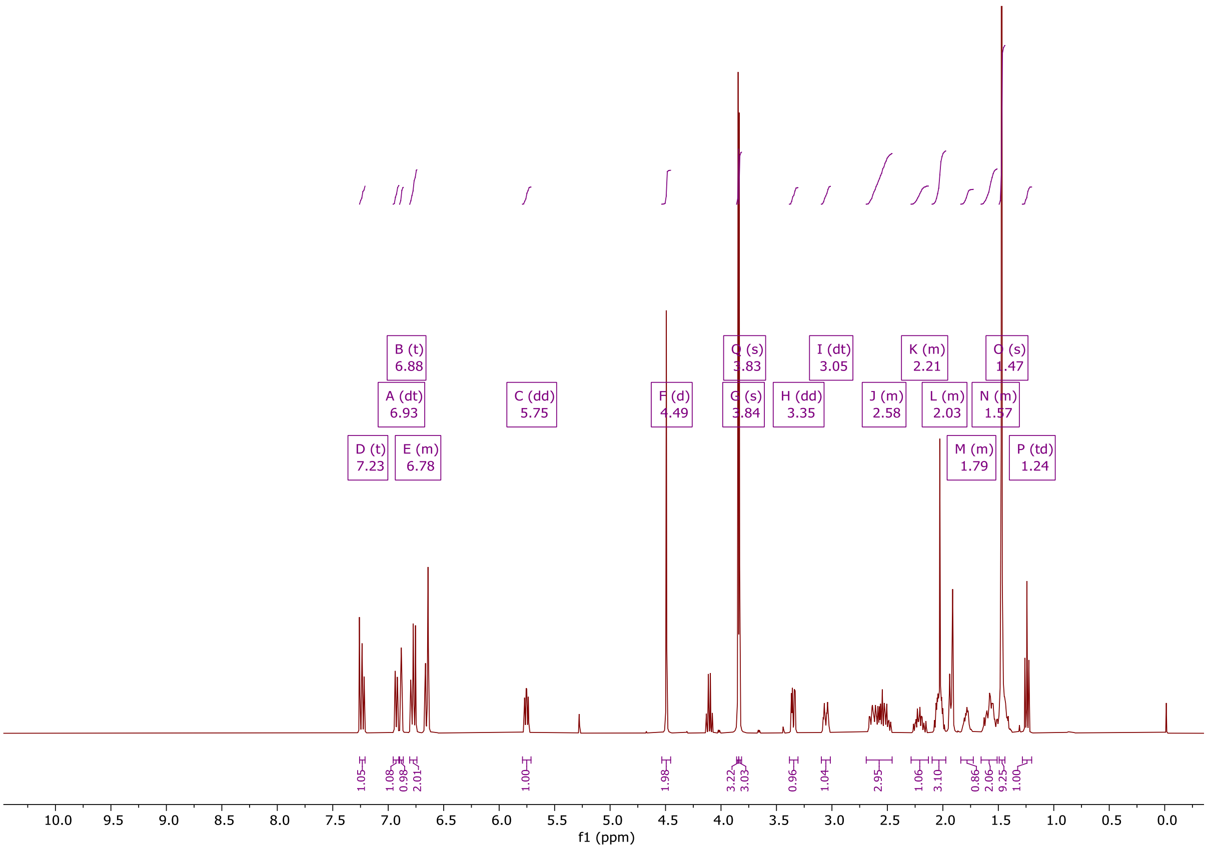


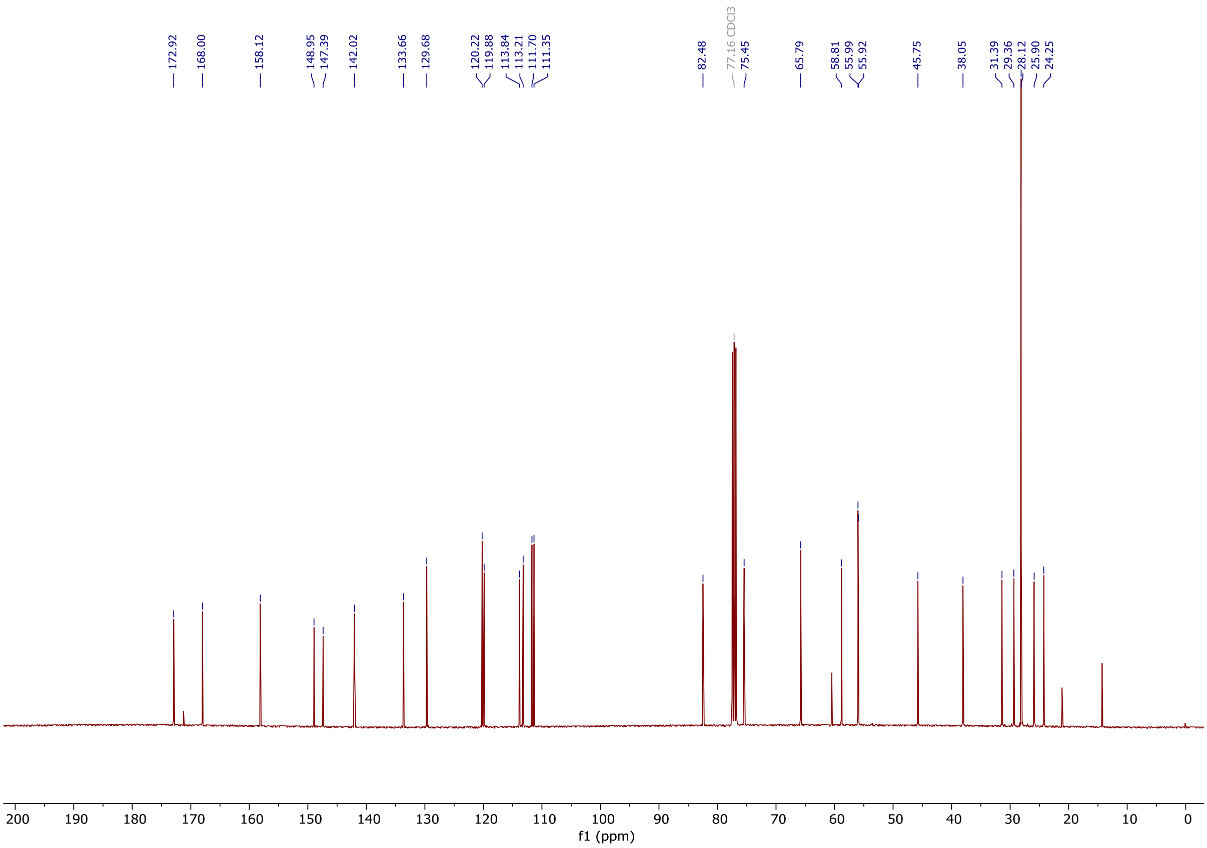

5.6. [1]H and [13]C NMR spectra of 7

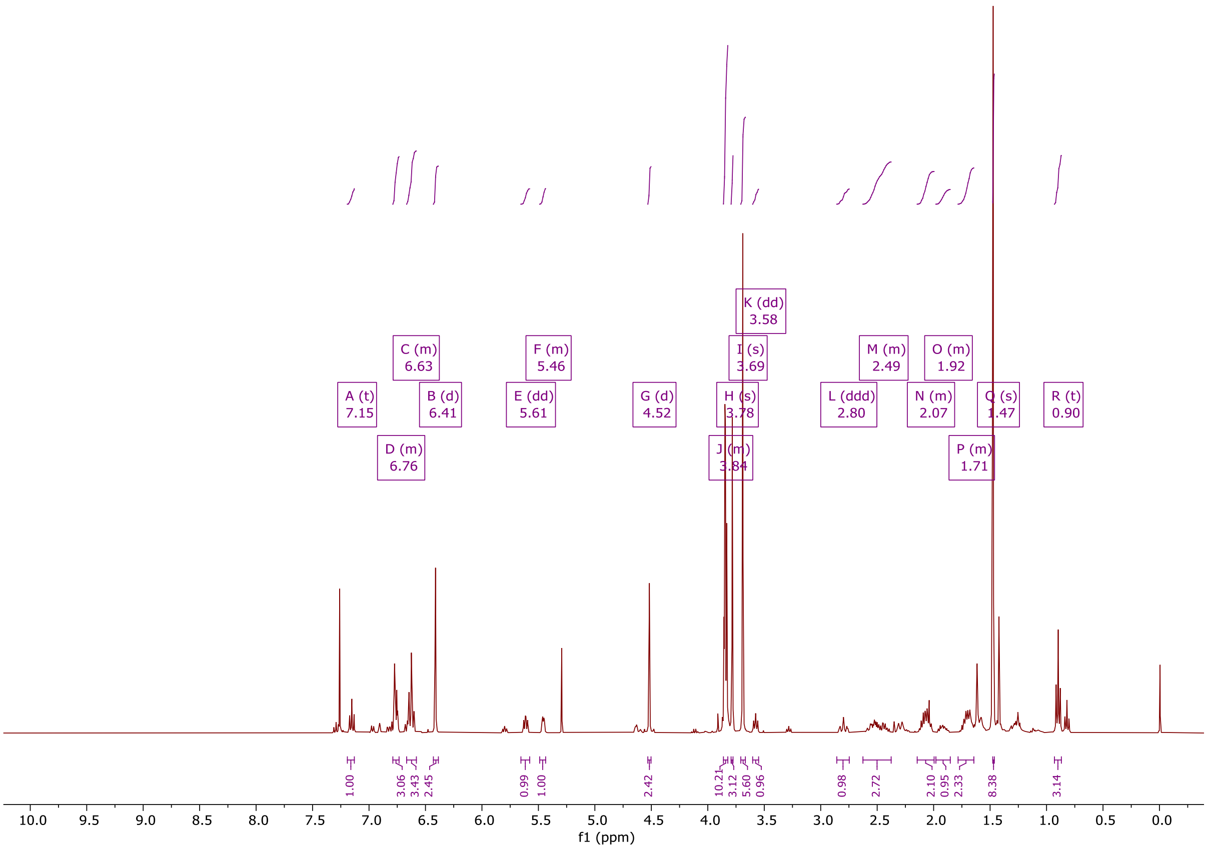


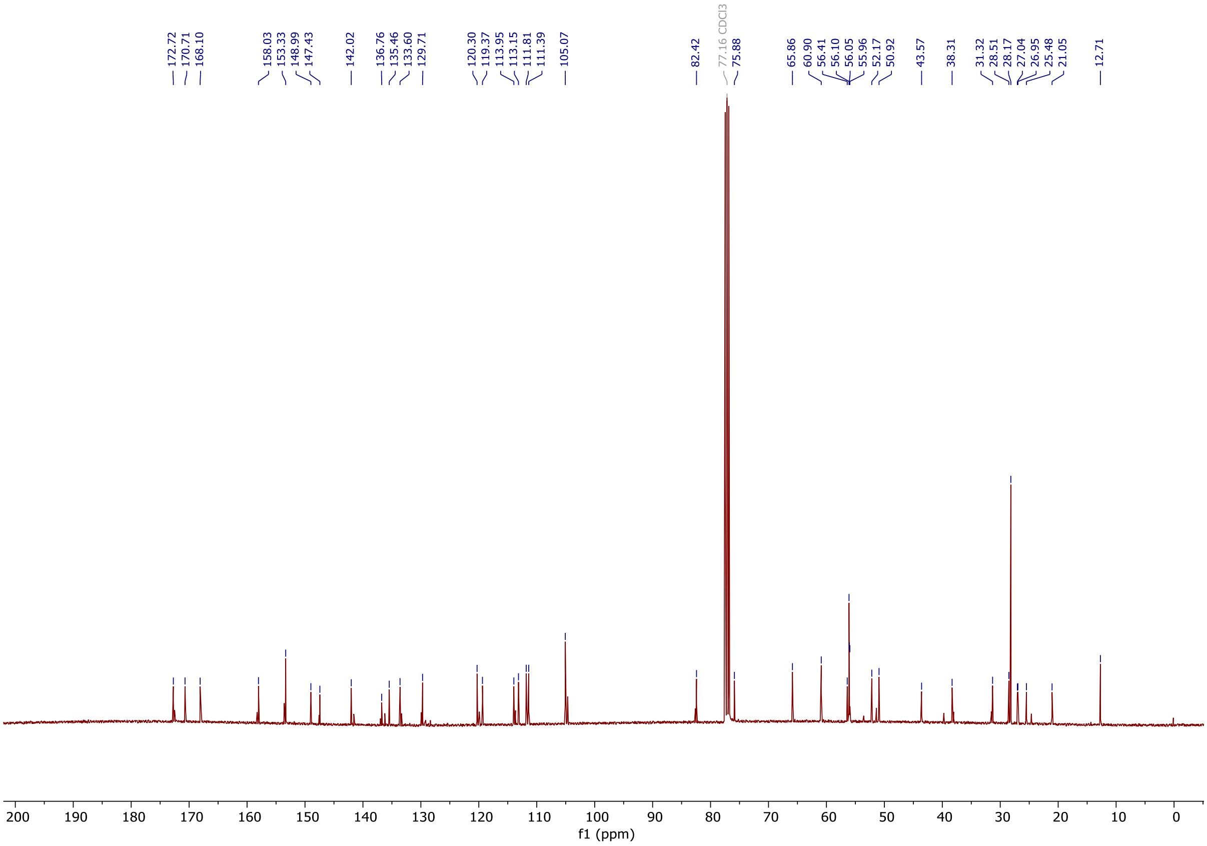

#### 5.7. ¹H and ¹³C NMR spectra of **8**

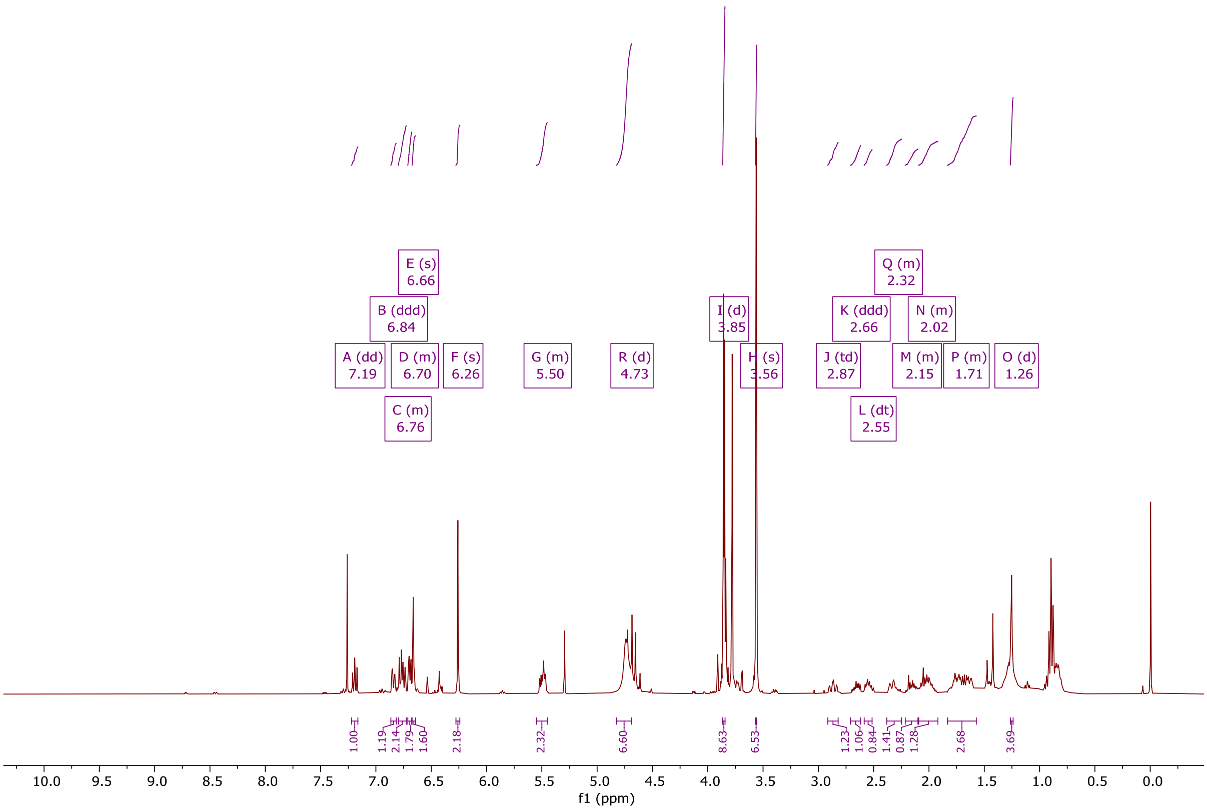


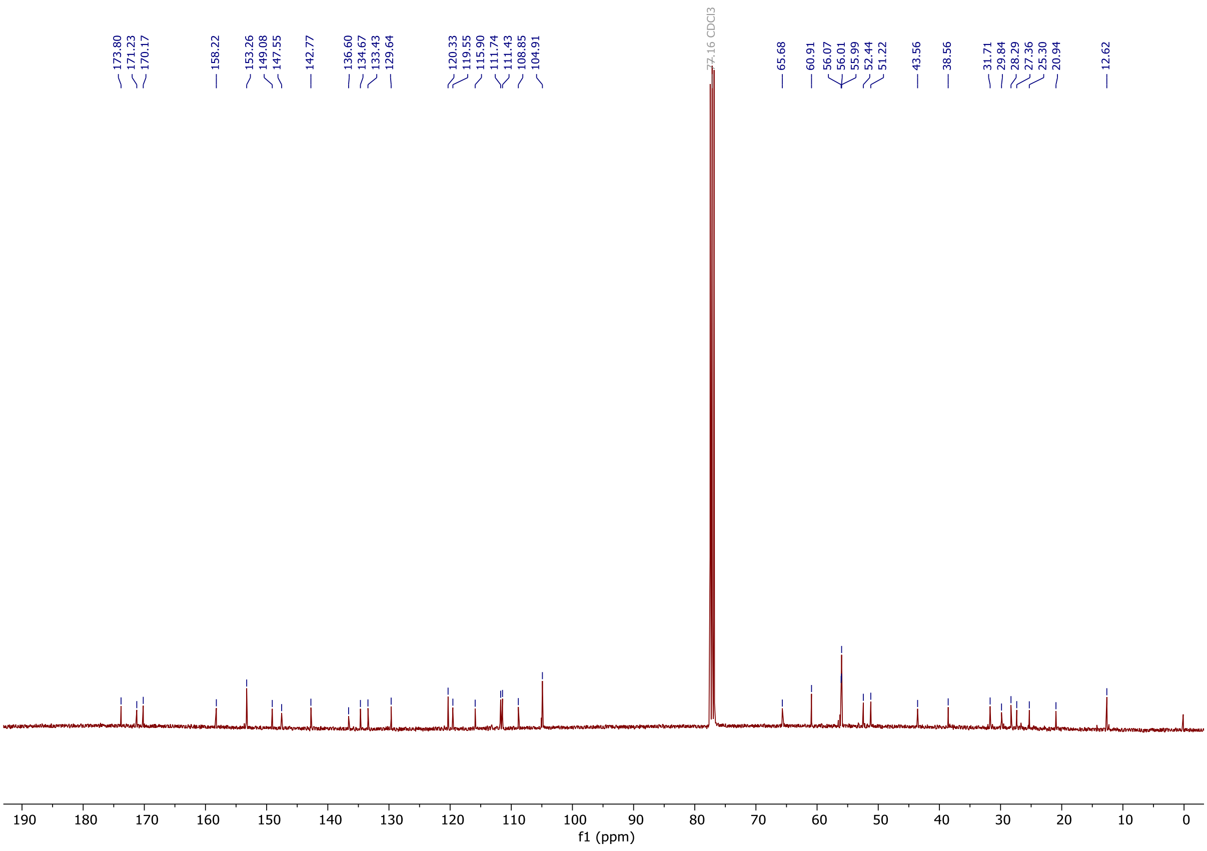

5.8. ¹H and ¹³C NMR spectra of **9** (dTAG-13)

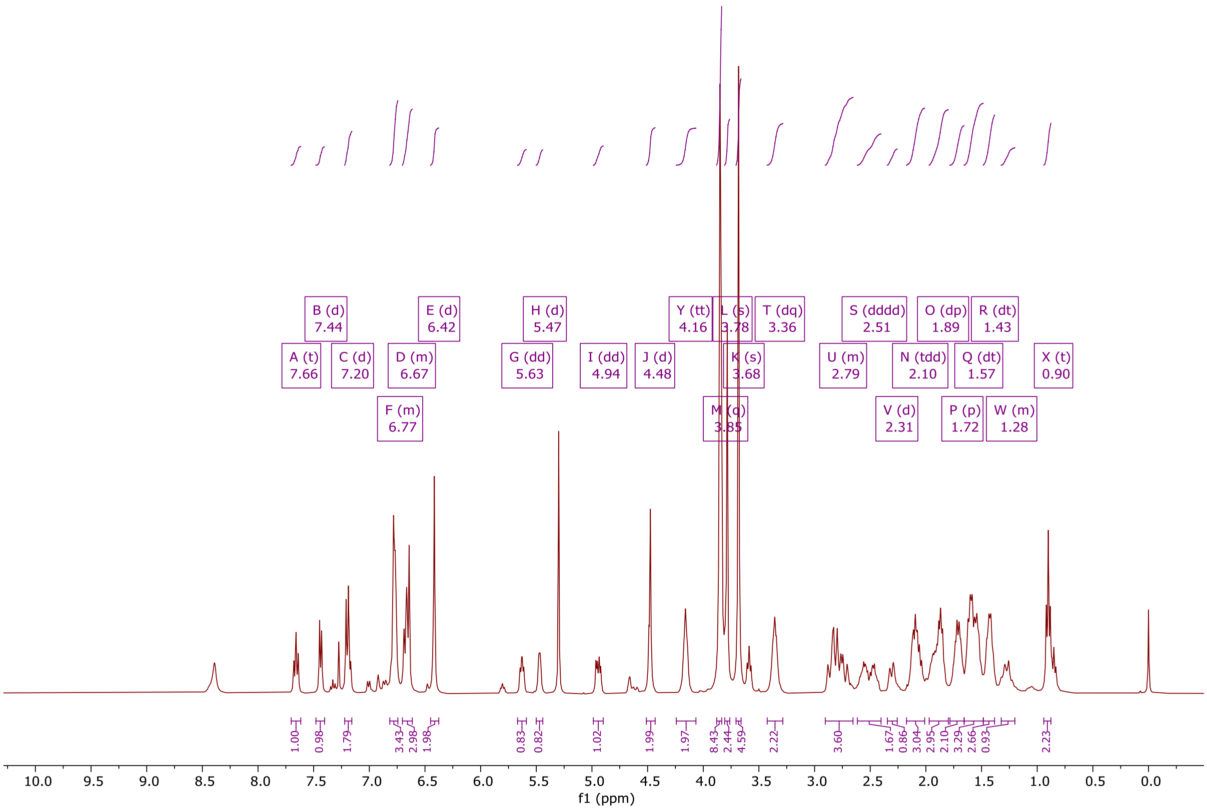


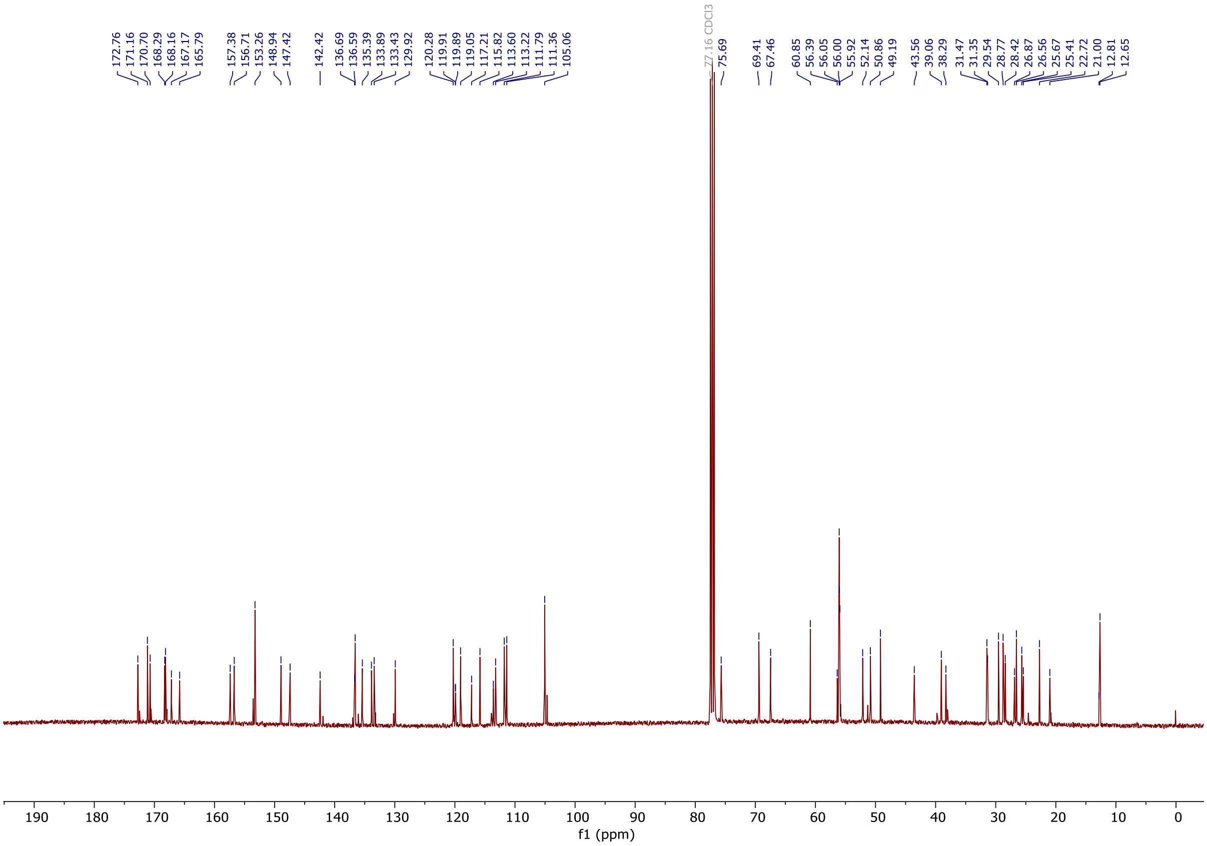

5.9. $^{1}H$ and $^{13}C$ NMR spectra of **12** (dTAG-v1)

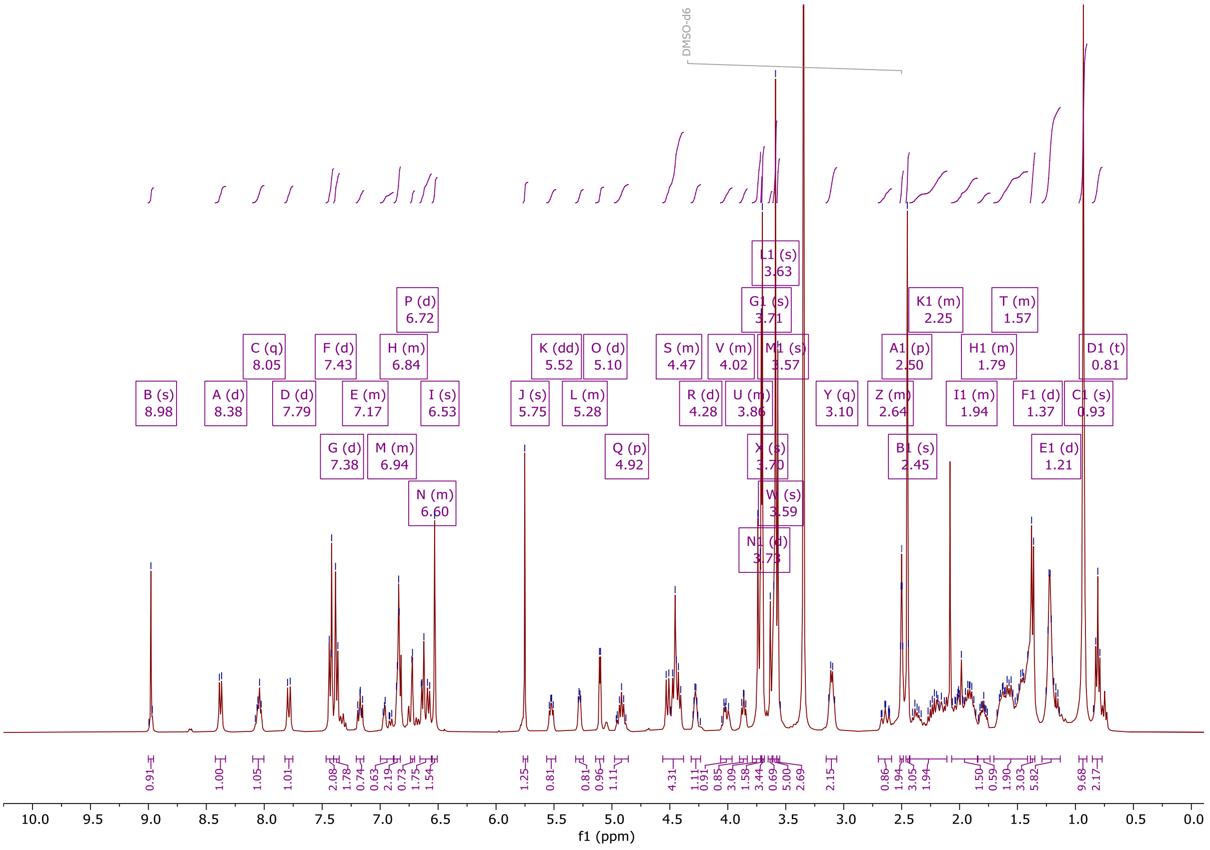

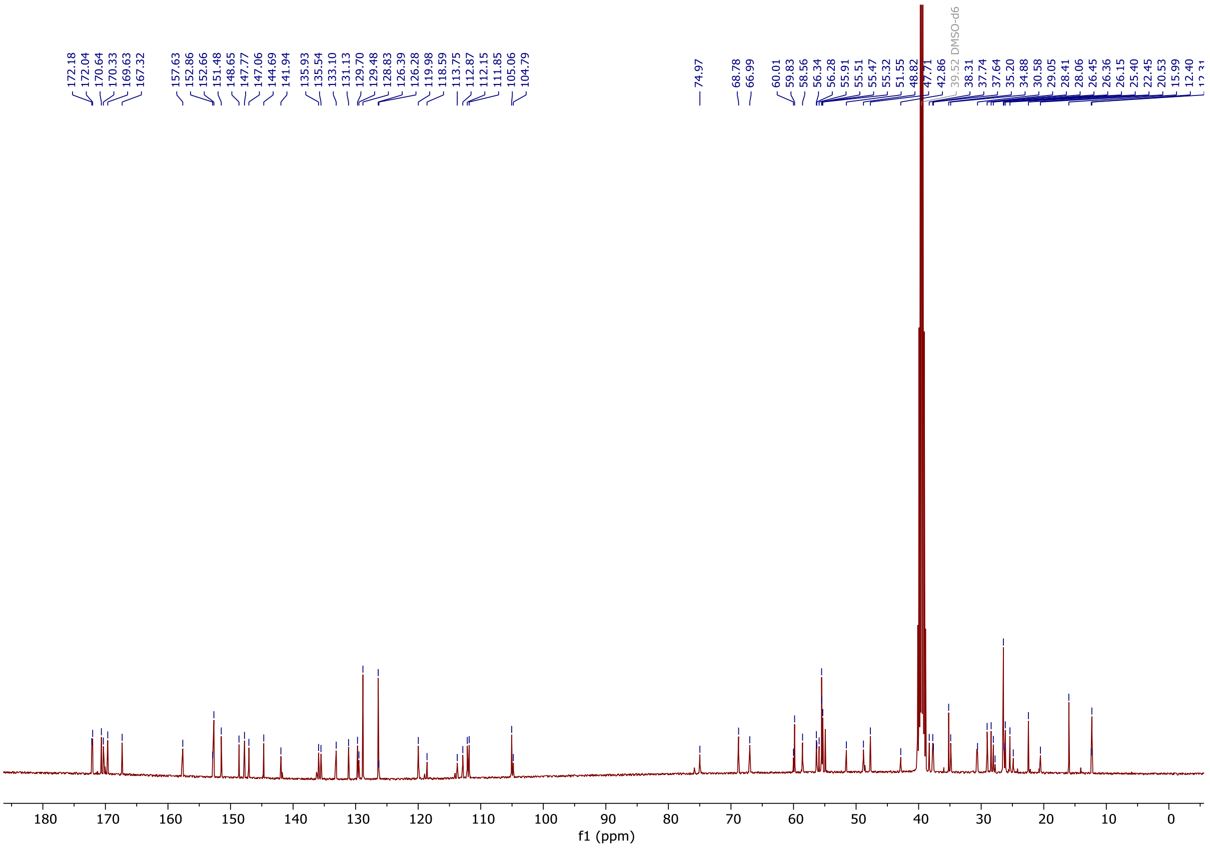

5.10. $^1$H and $^{13}$C NMR spectra of **14** (IGP002)

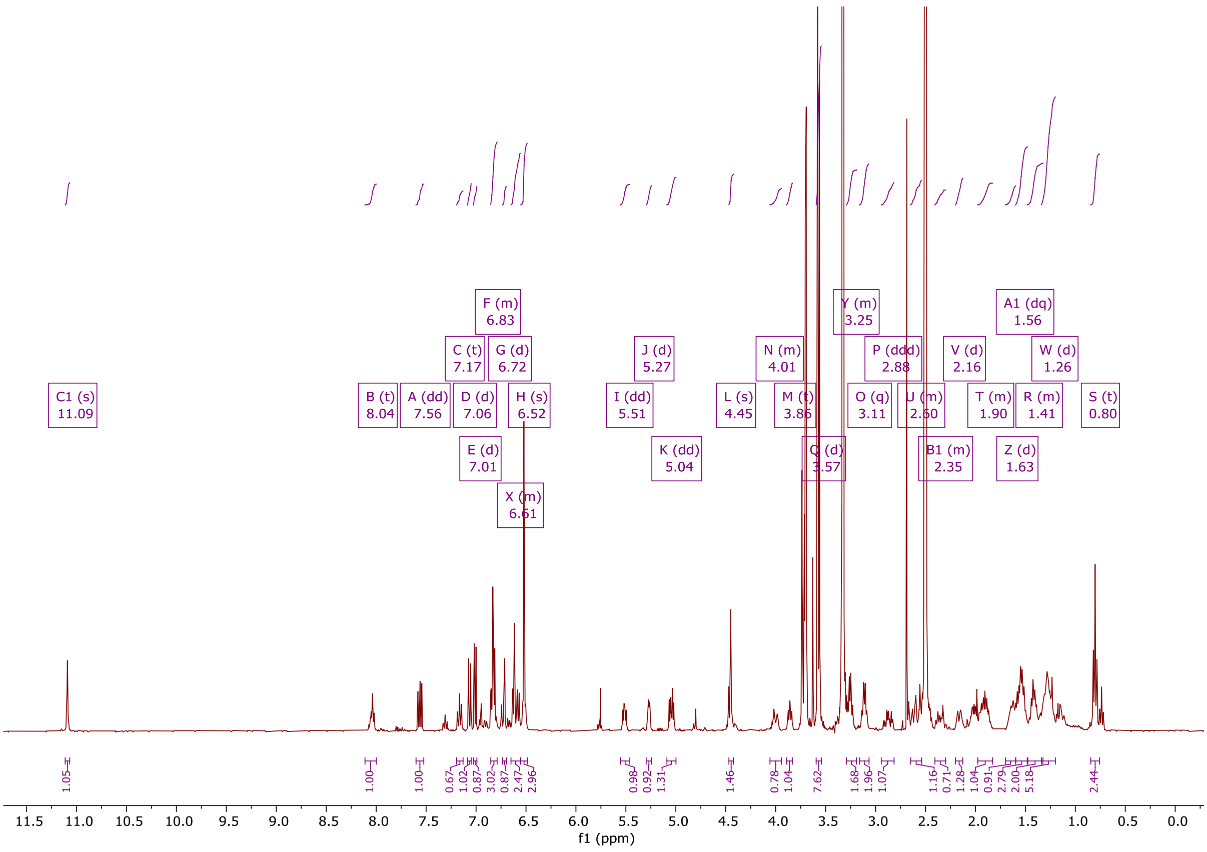

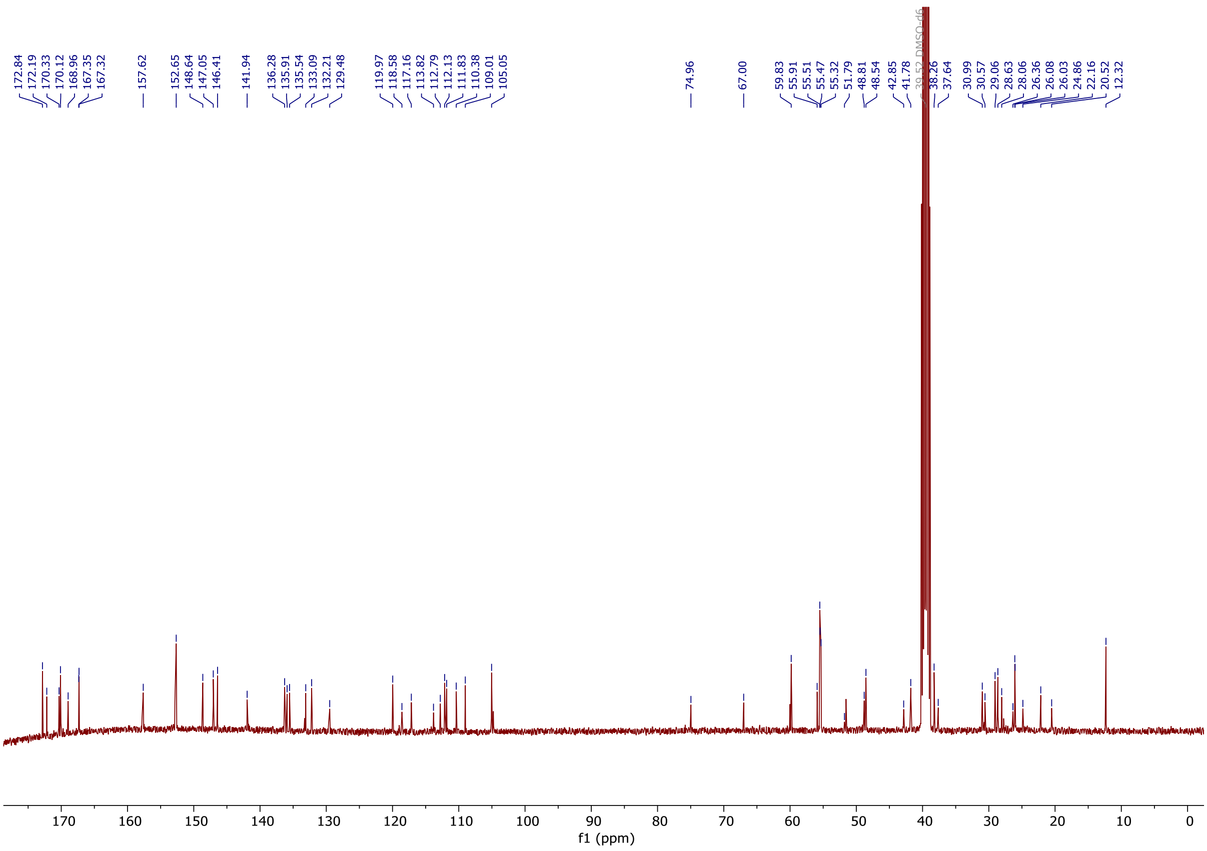
